# Evolutionary insights into glucose production in vertebrate development: new findings from Arctic lamprey (*Lethenteron camtschaticum*)

**DOI:** 10.64898/2026.05.24.727455

**Authors:** Marino Shimizu, Wataru Takagi, Fumiya Furukawa

## Abstract

Glucose has important roles in the development of the hematopoietic stem cells and the brain in vertebrate embryos; however, in most oviparous animals, the amount of glucose in the yolk is scarce. In zebrafish, gluconeogenesis takes place in the yolk syncytial layer (YSL), an extraembryonic tissue that surrounds the yolk. Gluconeogenic activity have also been observed in extraembryonic YSL-like tissue or endoderm-derived tissues in cloudy catshark, sterlet, and western clawed frog during development. However, it remains unclear when such ability was acquired or how it changed over the evolution of vertebrates. In this study, we used the Arctic lamprey, a cyclostome sister group of jawed vertebrates, to compare changes in metabolite levels and gluconeogenic gene expression patterns during development. Also, gluconeogenic activity was assessed using ^13^C-labeled substrates. Our metabolite analysis revealed that glucose levels increased during development and that glycerol was actively metabolized to produce glucose. In addition, many gluconeogenic genes were expressed in the muscle, notochord, and epithelium, making a striking contrast to previous observations in the above-mentioned vertebrates. Genomic DNA sequence motif analysis using HOMER and MEME identified common transcription factors binding motifs in the upstream regions of *g6pc1/2* and *fbp1* across vertebrate lineages. Among them, interestingly, the binding motif for HNF4A was not detected in *g6pc1/2* and *fbp1* genes of cyclostomes, suggesting distinct transcriptional regulation of gluconeogenesis in cyclostomes. These results indicate that gluconeogenesis is an essential process during development across vertebrate lineages, including cyclostomes, although the tissues and regulatory mechanisms for this function vary among lineages.

## Introduction

In oviparous animals, yolk functions as the sole nutritional source until the onset of feeding. The major components of yolk are proteins and lipids, whereas carbohydrates, which serve as substrates for energy production and biosynthetic pathways, are present only in small amounts. Carbohydrate metabolism during early development has traditionally received little attention. However, studies in the zebrafish, a vertebrate model organism, have suggested that glucose is essential for the normal development of the brain and hematopoietic stem cells (Harris et al., 2013; Jensen et al., 2006). Recently, it has been demonstrated that zebrafish supplement glucose through gluconeogenesis in the yolk syncytial layer (YSL), an extraembryonic tissue surrounding the yolk, utilizing yolk-derived substrates as precursors (Furukawa et al., 2024). These findings suggest that, during early stages when pivotal metabolic organs, such as the liver, are not yet fully developed, the YSL functions to take over their roles of synthesizing and supplying glucose.

The YSL is formed during the meroblastic cleavage characteristic of teleosts (Kimmel and Robert, 1985). Cartilaginous fishes, which share a similar cleavage pattern, also possess a YSL-like structure (Hamlett et al., 1984). In cartilaginous fishes, this tissue has likewise been shown to participate in yolk degradation, transport, and gluconeogenesis (Lechenault et al.,1993; Honda et al., 2022; Shimizu et al., 2024), indicating functional similarities to YSL in teleosts. In contrast, vertebrates exhibiting holoblastic cleavage cannot utilize yolk through such YSL-mediated mechanisms. In a chondrostean sturgeon, *Acipenser ruthenus*, or sterlet, yolk degradation and absorption occur in the embryonic endoderm that develops around the yolk mass (Shah et al., 2024), and this tissue has also been shown to perform gluconeogenesis (Shimizu et al., 2025). However, this mechanism appears to be associated with the unique mode of gut formation in sturgeons. Furthermore, in an anuran amphibian, *Xenopus tropicalis*, gluconeogenesis has been suggested to occur in the epidermis and in endodermal tissues that later differentiate into the liver (Aoki et al., 2026). In amphibians, all three germ layers contribute directly to embryonic body formation, and extraembryonic tissues are absent. Such a developmental mode is suggested to have been secondarily established through the incorporation of extraembryonic vegetal cell masses into the embryonic endoderm (Takeuchi et al., 2009).

Taking together, these findings suggest that gluconeogenesis during early development is a broadly conserved and important phenomenon among oviparous gnathostomes (jawed vertebrates). However, whether and how developmental gluconeogenesis occurs in cyclostomes, the only extant sister group of gnathostomes, remains unknown. Such knowledge may provide key insights into the evolutionary origin of developmental gluconeogenesis across vertebrates. Cyclostomes include the classes Myxini (hagfishes) and Petromyzontida (lampreys). These two lineages diverged more than 400 million years ago (Kuraku et al., 2009) and exhibit substantial differences in morphology and physiology. Their modes of early development also differ markedly: hagfishes possess yolk-rich eggs and undergo meroblastic cleavage, whereas lampreys exhibit holoblastic cleavage. In particular, because the common ancestor of vertebrates is believed to have exhibited holoblastic cleavage (Chea et al., 2005), elucidating the mechanisms of yolk utilization and gluconeogenesis in lampreys may provide important insights into how metabolic systems evolved during early vertebrate development.

As summarized by Richardson and Wright (2003), lampreys were intensively studied in embryological research from the late nineteenth to the early twentieth century. Although organogenesis during the prolarval stage—prior to yolk absorption and before reaching the ammocoete larval stage—has been described in detail with extensive illustrations (Tahara, 1988; Richardson and Wright, 2003; Richardson et al., 2010), relatively few studies have focused on the physiological aspects of this period. In particular, there is little information on how yolk is utilized during this stage.

In this study, we analyzed changes in metabolite levels during development in Arctic lamprey *Lethenteron camtschaticum*. Because an increase in glucose levels was detected during embryogenesis, the prolarvae were cultured with ¹³C-labeled metabolites, and their metabolic dynamics were traced using liquid chromatography–tandem mass spectrometry (LC-MS/MS). In addition, gene expression analyses were conducted to support gluconeogenic activity and to determine its spatial localization. Based on these results, we discuss the physiological and evolutionary significance of gluconeogenic tissues in developing vertebrates.

## Materials and methods

### Animals

Embryonic and adult Arctic lampreys, *Lethenteron camtschaticum* collected from the Ishikari River, were maintained at the Atmosphere and Ocean Research Institute, the University of Tokyo, in dechlorinated freshwater at 11°C under a 12 h light/12 h dark photoperiod until artificial fertilization experiments were initiated in May. Fertilized eggs were primarily maintained at 12°C in the dark. Although minor differences in developmental progression have been reported among lamprey species, embryonic stages are considered broadly comparable across species (Richardson et al., 2010; Li et al.,2019). In this study, embryonic stages of Arctic lampreys were identified according to the detailed staging table described by Tahara (1988), in which stage (st.) 24 approximately corresponds to the hatching stage. All experiments were approved by the Animal Care and Use Committee of the Atmosphere and Ocean Research Institute, the University of Tokyo (P22-6), and were conducted in accordance with the ARRIVE guidelines.

### Metabolomic analysis

Metabolomic analysis was performed by LC–MS according to previously reported methods (Shimizu et al., 2024; 2025) using samples from the 1-cell stage, 2-cell stage, sts.7–8, 16–17, 17–18, 21, 22–23, 24, 27, and 30. From st. 24, the hatched prolarvae were sampled. For each developmental stage, 30 individuals were pooled into a 1.5 mL tube (*n* = 6). Distilled water (100 μL), 100% methanol (200 μL), and 5 μL of internal standard solution [L-methionine sulfone, 2-(*N*-morpholino) ethanesulfonic acid, D-camphor-10-sulfonic acid, 5 mM each] were added, and the samples were homogenized. Subsequently, 200 μL of chloroform was added and mixed thoroughly. After incubation on ice for 10 min, the samples were centrifuged at 4°C and 13,000 rpm for 10 min. The aqueous phase was collected and subjected to liquid chromatography-mass spectrometry (LC-MS) analysis.

LC-MS analysis was performed using an LC system (Prominence; Shimadzu, Kyoto, Japan) connected to a TripleTOF 5600+ mass spectrometer (AB Sciex, Framingham, MA, USA). A Shodex HILICpak VG-50 2D column (Resonac, Tokyo, Japan) was used for separation. The mobile phases consisted of solvent A (acetonitrile) and solvent B (0.5% aqueous ammonia). The initial condition was 8.8% B, and the gradient program was as follows: 8.8% B (8 min), 95% B (14 min), 95% B (17 min), 8.8% B (19 min), and 8.8% B (24 min). The flow rate was set at 0.2 mL/min, and the column temperature was maintained at 60°C.

For the measurement of phosphorylated sugars and amino acids, a Shodex HILICpak VT-50 2D (Resonac, Tokyo, Japan) column was used. The mobile phase consisted of acetonitrile (A) and 25 mM ammonium formate (B) at a constant ratio of A:B = 20:80. The flow rate was 0.3 mL/min, and the column temperature was maintained at 60°C. LC-MS parameters used for analysis are summarized in Table S1A. Metabolite concentrations were calculated based on parallel measurements of serially diluted standard solutions.

### Isotope tracing

After anesthesia, st. 24–25 prolarvae were trisected. Glycerol-¹³C□ (Taiyo Nippon Sanso, Tokyo, Japan), L-lactate sodium salt-¹³C□ (Cambridge Isotope Laboratories, Tewksbury, MA, USA), L-alanine-¹³C□ (Merck, Darmstadt, Germany), and glutamine-^13^C□ (Cambridge Isotope Laboratories) were diluted to 5 mM in Steinberg’s solution (in mM: 5.81 NaCl, 0.271 HCl, 0.0835 MgCl□, 0.033 CaCl□, 0.067 KNO□, 0.0835 H□SO□, 0.46 Tris, pH 7.4; Palmer and Slack., 1970). Twenty trisected prolarvae were pooled per dish and incubated for 3 hours in Steinberg’s solution containing the labeled tracers or without tracers as a control (*n* = 8). Metabolites were subsequently extracted as described above and analyzed using an LCMS-8045 system (Shimadzu Corporation, Kyoto, Japan) under the same chromatographic conditions described for metabolomic analysis. LC-MS/MS parameters used for isotope tracing analysis are summarized in Table S1B. Relative enrichment of tracer-derived metabolites (isotopologues) was determined by measuring the mass (M)-shifted isotopologues and expressed as %M+0, a percentage relative to the unlabeled form (M+0).

### Histological analyses

To examine morphological changes during lamprey development, hematoxylin and eosin (H&E) staining was performed. Sagittal sections from developmental sts 24, 25, 28, 29, and 30 were used for the analysis. After deparaffinization, sections were stained with hematoxylin and eosin. Following staining, the sections were mounted with Permount and coverslipped. Images were acquired using a BX53 microscope (Olympus, Tokyo, Japan).

### RNA probe synthesis and *in situ* hybridization

DIG-labeled RNA probes were generated for *in situ* hybridization analysis. Target cDNA fragments were first amplified using the primers listed in Table S2. The purified cDNA fragments were used as templates for *in vitro* transcription in the presence of digoxigenin (DIG)-UTP with either T7 or SP6 RNA polymerase (Roche Diagnostics, Basel, Switzerland). The resulting cRNA probes were treated with DNase I (Roche Diagnostics) and subsequently purified using the RNeasy MinElute Cleanup Kit (Qiagen, Hilden, Germany).

*In situ* hybridization was performed according to Shimizu et al. (2025). Prolarvae at sts.24 and 27 were fixed overnight at 4°C in 4% paraformaldehyde (PFA) in PBS-DEPC, embedded in paraffin, and sectioned at 5 µm thickness. The sections were affixed to glass slides. After deparaffinization, sections were treated with 2 µg/mL proteinase K for 20 minutes at room temperature. Prehybridization was carried out using hybridization mix (HM+; 50% formamide, 5× SSC, 0.01% Tween 20, 500 µg/mL yeast tRNA, 50 µg/mL heparin). Hybridization was performed overnight at 70°C in 200 µL HM+ containing 65 ng of either antisense or sense (negative control) DIG-labeled RNA probe per 200 µL. Excess probe was removed by washing with 0.2× SSC at 70°C. The sections were then blocked with 5% skim milk in PBS-DEPC for 1 hour at 4°C and incubated for 1 hour at 4°C with anti-DIG antibody (Roche Diagnostics) diluted 1:10,000 in the blocking solution. mRNA signals were visualized by BCIP/NBT color development and photographed with the BX53 microscope.

### Periodic acid–schiff (PAS) staining

After deparaffinization, sections of st. 24 lampreys were immersed in 0.5% acetic acid for 10 min. For glycogen digestion, negative control sections were treated with 3 U/mL glucoamylase in 0.5% acetic acid for 2 h at 37°C, whereas adjacent sections were incubated in acetic acid alone. The sections were subsequently oxidized in 0.5% periodic acid for 10 min, rinsed with water, and stained with Schiff’s reagent (Muto Pure Chemicals, Tokyo, Japan) for 3 min. After washing in sulfite solution, the sections were counterstained with hematoxylin for 3 min, mounted with Permount, and observed under the BX53 microscope. PAS-positive glycogen signals were evaluated by comparison with corresponding glucoamylase-treated control sections.

### Tissue distribution analysis

Total RNA was extracted from nine tissues (brain, gill, epidermis, muscle, notochord, heart, kidney, liver, and intestine) of adult lampreys (*n* = 6) using TRI Reagent (Molecular Research Center, Cincinnati, OH, USA). After DNase I treatment, RNA was reverse-transcribed using the ReverTra Ace qPCR RT Kit (Toyobo, Osaka, Japan). Quantitative PCR (qPCR) was performed using a Thermal Cycler Dice Real-Time System II (Takara Bio, Shiga, Japan) and Luna Universal qPCR Master Mix (New England Biolabs, Ipswich, MA, USA). Primer sequences are listed in Table S2. PCR reactions were carried out in parallel with plasmid standard solutions containing each target gene fragment at known concentrations (10¹–10□ copies/µL). Gene expression levels in each sample were calculated based on the corresponding standard curves. After amplification, dissociation curve analysis was performed to confirm the specificity of the PCR products.

### Molecular phylogenetic analysis

Amino acid sequences of orthologs of G6PC1/2/3 from 58 species (3 cephalochordate, 6 cyclostomes, and 49 gnathostomes) and HNF4A/B/G (also referred to as HNF4α/β/γ) from 56 species (3 cephalochordate, 5 cyclostomes, and 48 gnathostomes) were obtained from NCBI and Ensembl. Accession numbers are listed in Tables S3 and S4. Amino acid sequences were aligned using MUSCLE implemented in MEGA11 (Tamura et al., 2021) and manually trimmed to remove poorly aligned terminal regions. Maximum-likelihood phylogenetic analyses were conducted using IQ-TREE v2.1.2 (Minh et al., 2020). The best-fit amino acid substitution model for each dataset was selected using ModelFinder (Kalyaanamoorthy et al., 2017) according to the Bayesian Information Criterion (BIC), resulting in JTT+F+R6 for the G6PC dataset and JTT+R5 for the HNF4 dataset. Branch support was assessed using 1,000 SH-aLRT replicates and 1,000 ultrafast bootstrap replicates (Hoang et al., 2018). The resulting trees were rooted using cephalochordate (amphioxus) sequences as outgroups and visualized using MEGA11 (Tamura et al., 2021).

### Motif analysis

To compare transcription factors regulating gluconeogenesis in cyclostomes and gnathostomes, upstream regions (3,000 bp) of the *g6pc1/2/3* and *fbp1*genes from 51 species (1 cephalochordate, 5 cyclostomes, and 46 gnathostomes) were obtained from NCBI and Ensembl. Common transcription factor binding motifs in the upstream regions of *g6pc1/2* or *fbp1* homologs were identified using HOMER (Hypergeometric Optimization of Motif EnRichment, v5.1; Heinz et al., 2010) and MEME (Multiple EM for Motif Elicitation, v4.11.2; Bailey et al., 2015; Bailey and Gribskov., 1988; Grant et al., 2011), and their positions were predicted.

## Results

### Metabolite analysis

The LC-MS/MS analysis revealed that glucose, gluconeogenic intermediates such as glucose-6-phosphate (G6P), fructose-6-phosphate (F6P), and phosphoenolpyruvate (PEP), as well as tricarboxylic acid (TCA) cycle–related metabolites were present at low levels until around the pre- to post-hatching stages (sts. 23–24) (Fig. 1). These metabolites increased markedly from st. 24 to 27. In contrast, glycogen, the intracellular storage form of glucose, together with its precursor UDP-glucose, tended to increase at earlier developmental stages compared with the above metabolic intermediates.

**Fig. 1.**
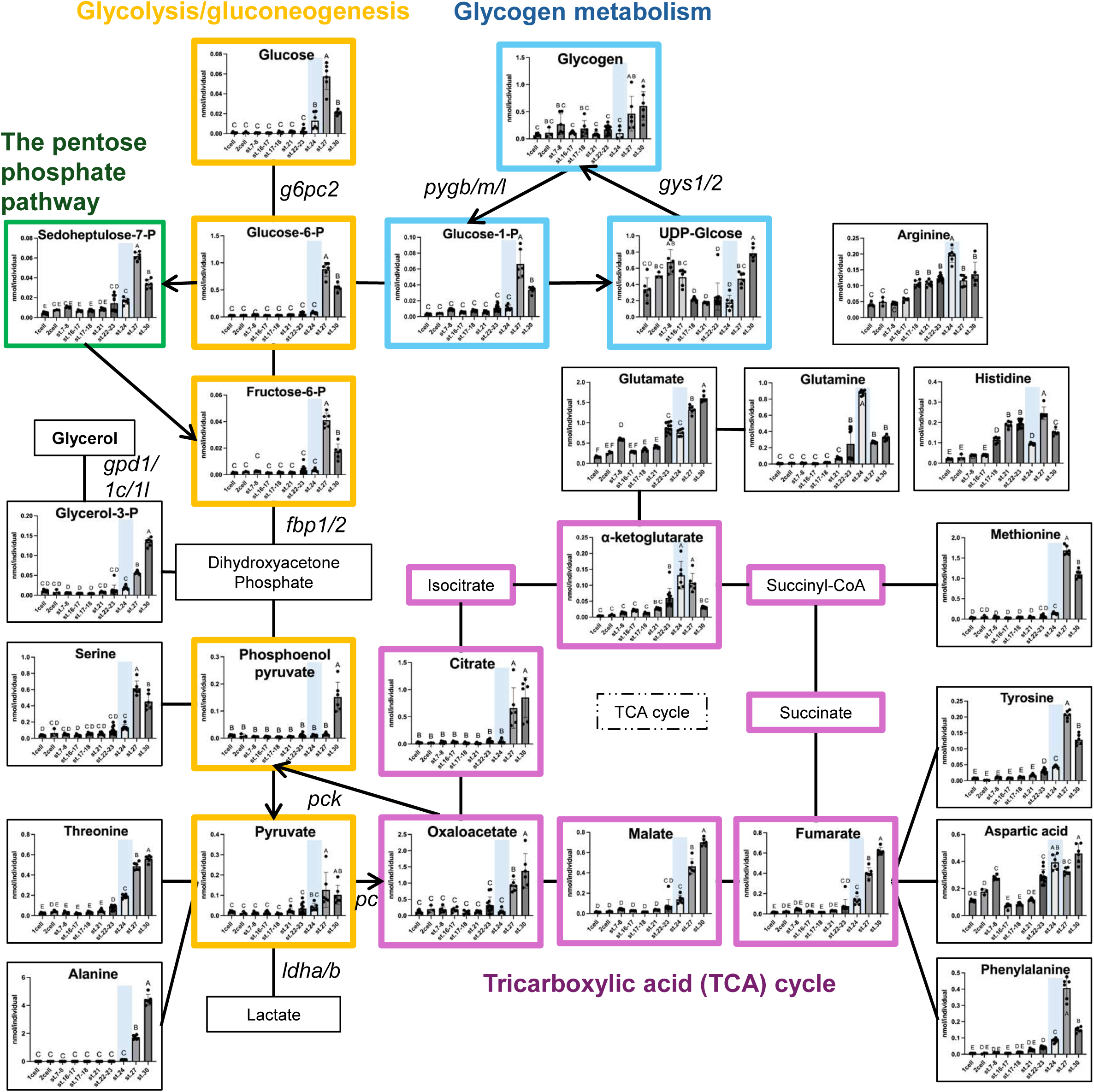
Changes in metabolite levels during the development of Arctic lamprey (*Lethenteron japonicum*). The graphs showing levels of metabolites were mapped onto the metabolic pathways. The horizontal axes represent developmental stages: 1 cell, 2 cell, and sts.7-8, 16-17, 17-18, 21, 22-23, 24, 27 and 30; the vertical axes represent nmol / individual. Data are presented as mean ± standard error (*n* = 6). Different letters indicate significant differences (*p* < 0.05) between groups. Tests for significant differences were performed by one-way ANOVA followed by Tukey’s *post-hoc* test. Light blue shadings indicate the hatching stage (st. 24). -P, phosphate.

### Isotope tracing

To clarify the activity of the gluconeogenesis pathway and the metabolic substrates in lamprey prolarvae, isotope tracing analysis was performed (Jang et al., 2018). Glycerol, alanine, and glutamine were selected as candidate substrates for gluconeogenesis. As a result, enrichment of glucose with a mass increase of +3 (M+3) was significantly higher in samples supplemented with ^13^C-labeled glycerol or lactate compared to the control (Fig. 2). In addition, M+3 isotopologues of G6P and F6P—intermediate metabolites of gluconeogenesis/glycolysis—were also enriched in these groups. Furthermore, M+3 isotopologues of sedoheptulose-7-phosphate and G1P were significantly increased in samples using glycerol as a substrate. Aspartate M+3 was enriched in all ^13^C-treated samples regardless of the substrate added.

**Fig. 2.**
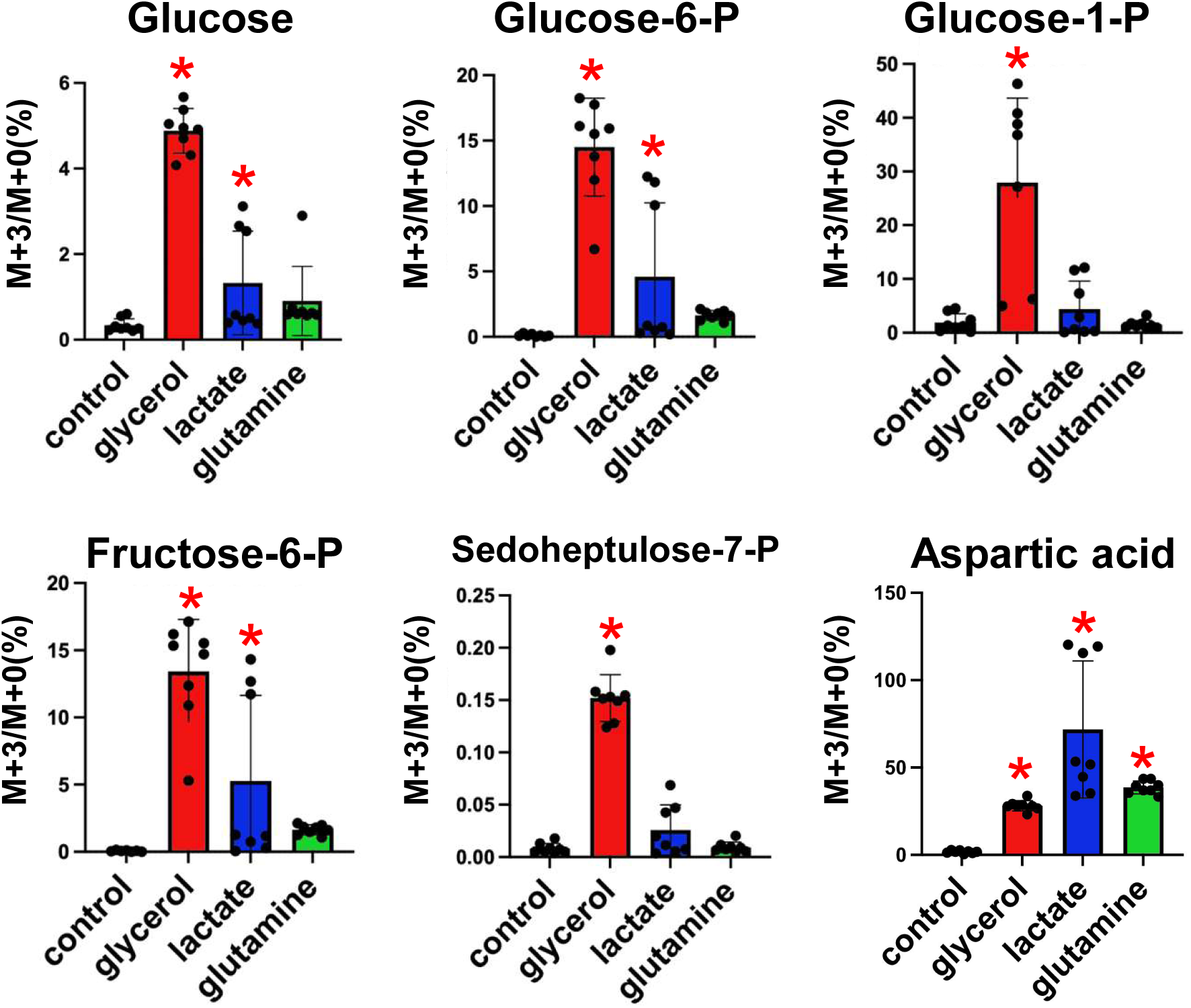
LC-MS/MS-based isotope tracing of lamprey prolarvae in st. 24-25. Abundance of M+3 isotopologues of glucose, glucose-6-phosphate (-P), fructose-6-P, glucose-1-P, and sedoheptulose-7-P, glutamine, and aspartic acid in the lamprey embryos after 3-hour incubation with ^13^C-labeled substrates. Horizontal axes show each ^13^C-labeled substrate and the control without them, and vertical axes show the levels of M+3 (as % to M+0). Data are presented as mean ± standard error (*n* = 8), and asterisks (*) indicate significant differences (*p* < 0.05) between the control and each group. Tests for significant differences were conducted with one-way ANOVA followed by Dunnett’s test.

### H&E staining

At st. 24, the head and neck regions had elongated and risen from the yolk sac, which consisted of numerous extraembryonic yolk cells (Fig. 3A) (Takeuchi et al., 2009). By st. 25, a developing liver was observed on the anterior tip of the yolk sac (Fig. 3B). At st. 27, yolk absorption had progressed further, and the trunk had become straight (Fig. 3C). From st. 28 onward, epithelialization of the yolk sac progressed from the anterior side (Fig. 3, D1 and D2), and this process was completed by st. 30 (Figs. 3, E and F).

**Fig. 3.**
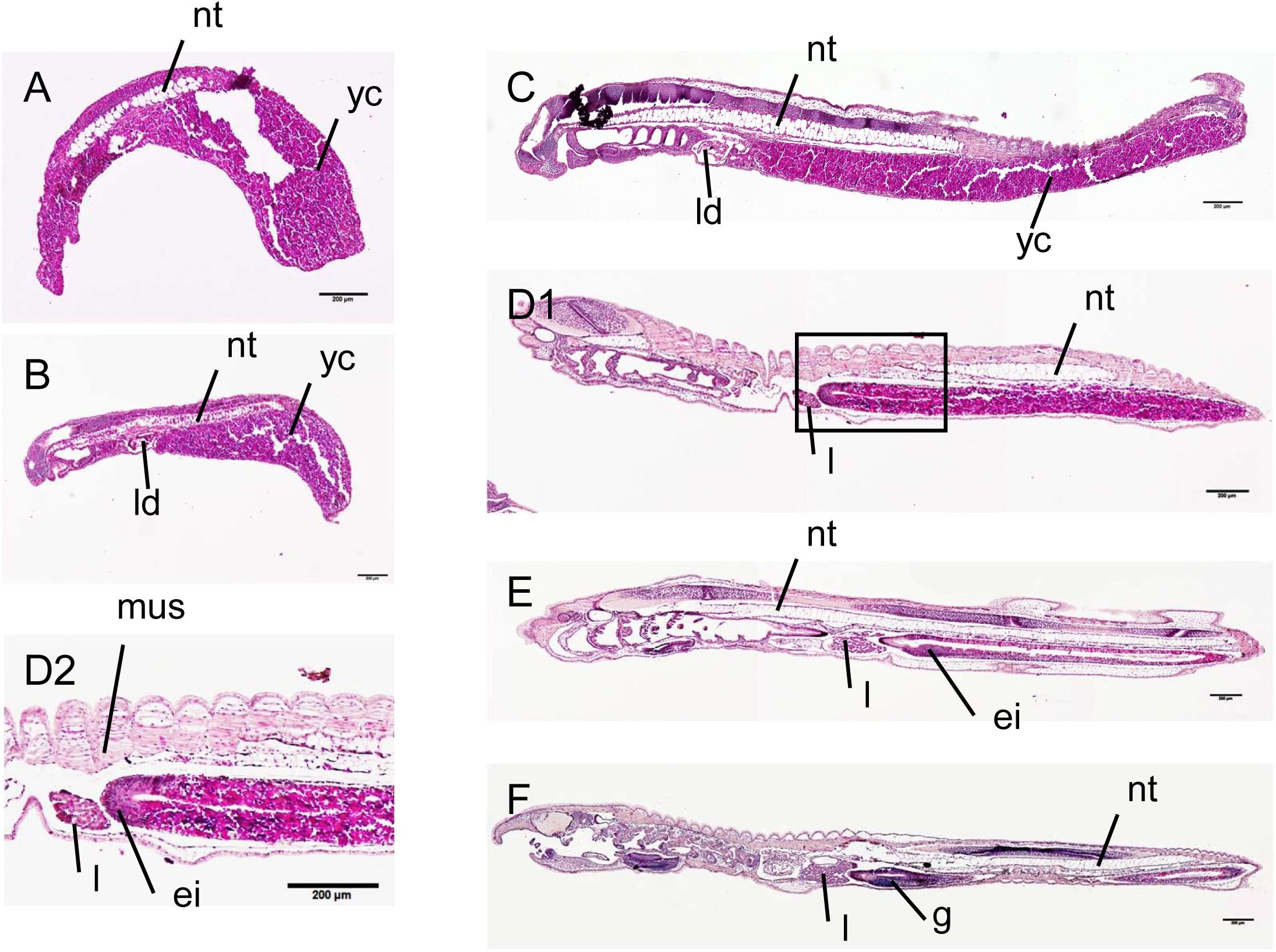
Hematoxylin-eosin (H&E) staining of Arctic lamprey embryos and prolarvae. Anterior sides to the left. (A) st. 24; (B) st. 25; (C) st. 27; (D1) st. 28; (D2) magnified view of the boxed area in D1; (E) st. 29; (F) st. 30. g, gut; ei, epithelial intestine; nt, notochord; l, liver; ld, liver diverticulum; mus, muscle; yc, yolk cell. Scale bars = 200 μm.

### In situ hybridization

Localization of gluconeogenesis- or glycogen metabolism-related transcripts in prolarvae was analyzed by *in situ* hybridization. In prolarvae at st. 24 and 27. At st. 24, *g6pc2, fbp1, pck, pc, ldha*, and *gpd1l* were expressed in the epidermis (Fig. 4). In muscle, signals for *g6pc2, fbp1/2, ldha, gpd1l*, and *gys1* were detected. *pc* was also expressed in the notochord (Fig. 4). At st. 27, in addition to these genes, *g6pc, fbp1*, *pck, ldha, gys1*, and *pygl* were also expressed in the notochord.

**Fig. 4.**
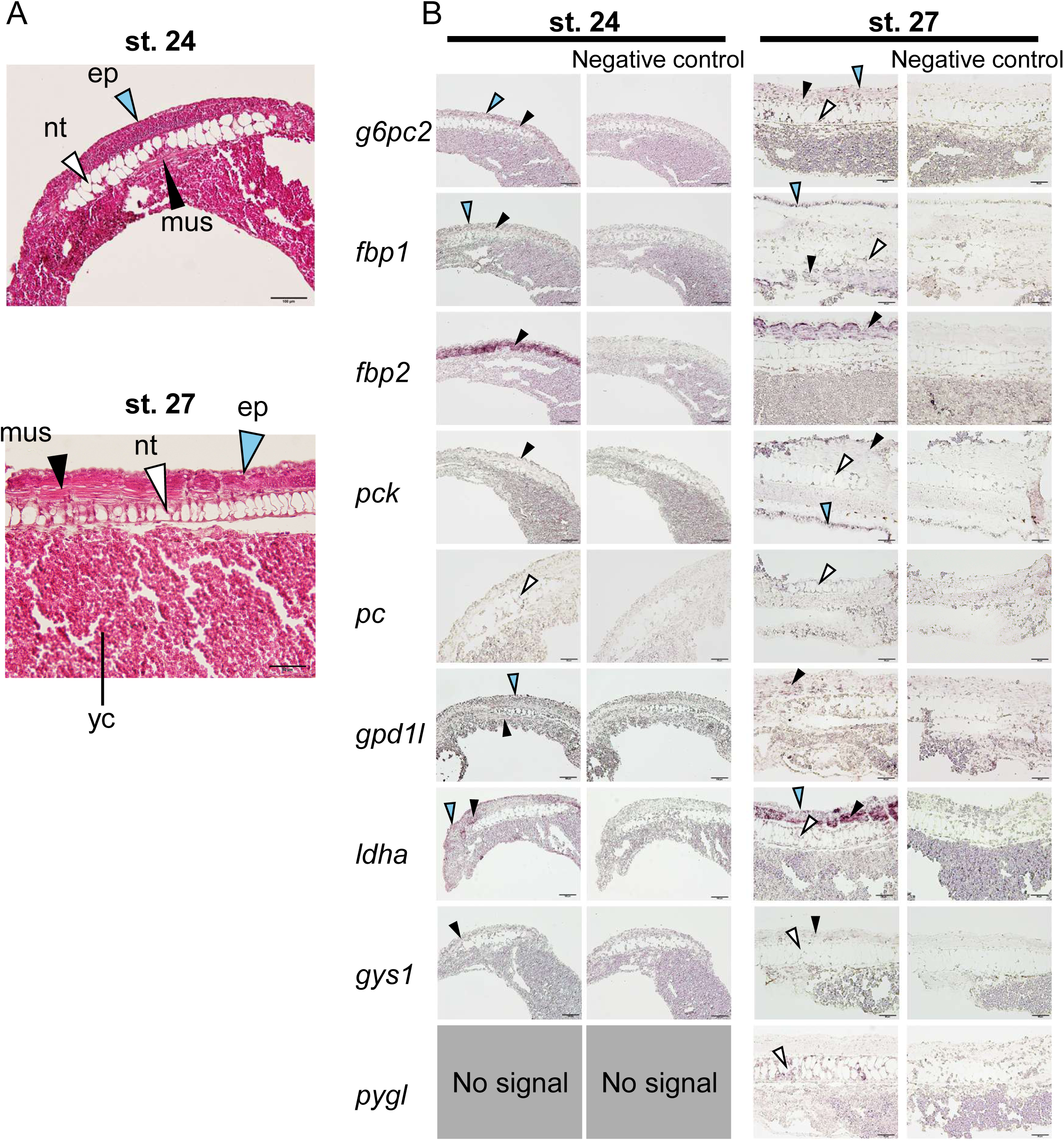
*In situ* hybridization analysis for genes related to gluconeogenesis and glycogen metabolism in the lampreys at st. 24 and 27. (A) Hematoxylin and eosin (H&E)-stained sections showing tissue morphology at st. 24 and 27. (B) *In situ* hybridization analysis. For all transcripts, sense probes (right) were used as negative controls for antisense probes (left), which showed true signals. Positive signals for transcripts are indicated by arrowheads. Black, white, and cyan arrowheads indicate muscle (mus), notochord (nt), and epidermis (ep), respectively. yc, yolk cell. (st. 24); 100 μm (st. 27), Scale bars = 50 μm.

### PAS staining

PAS staining in st. 24 samples revealed glycogen signals in the muscle, notochord, and epidermis (Fig. 5). In the epidermis, which consists of a single cell layer, PAS-positive signals were observed along the inner side of the cells. These PAS-positive signals were markedly reduced in glucoamylase-treated sections, confirming that the staining represented glycogen.

**Fig. 5.**
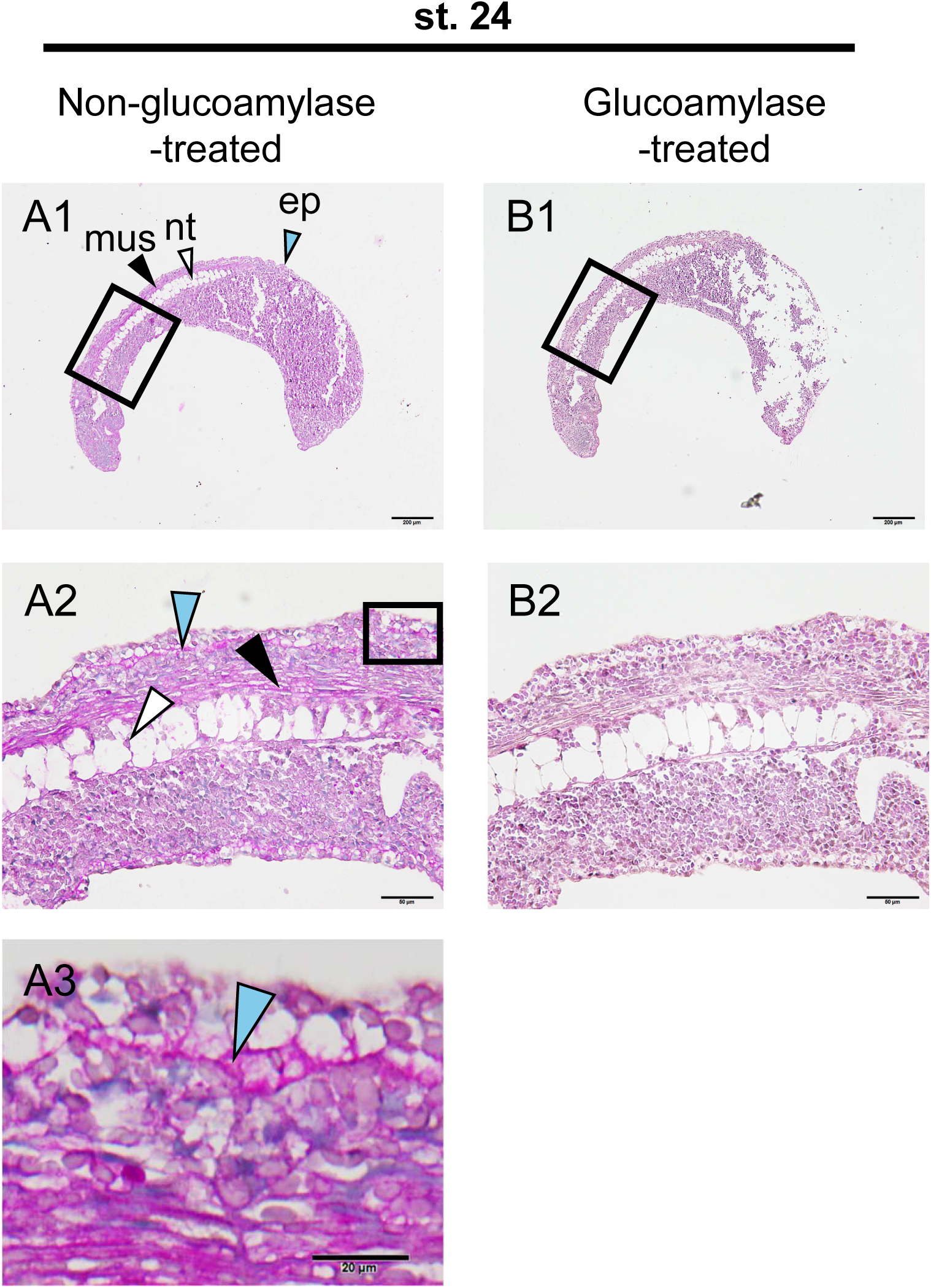
Periodic acid-Schiff (PAS) staining of the lamprey prolarvae at st.24. Adjacent sections were treated with glucoamylase prior to PAS staining, which served as negative controls (right) against the glycogen signal shown in the sections without glucoamylase treatment (left). Positive signals are indicated by arrowheads. Black, white, and cyan arrowheads indicate muscle (mus), notochord (nt), and epidermis (ep), respectively. (A1, B1) Whole embryo view. (A2, B2) Higher-magnification view of the region indicated in A1 and B1. (A3) Detail from the boxed area in A2. Scale bars = 200 μm (A1, B1), 50 μm (A2, B2), 20 μm (A3).

### Tissue distribution analysis

Real-time PCR was performed on nine tissues from adult lampreys. Most gluconeogenic genes did not show clear differences in expression levels among the tissues. In contrast, *g6pc2* exhibited higher expression in the notochord, kidney, and liver compared to most other tissues (Fig. 6). Additionally, *fbp2* showed higher expression in muscle than in other tissues.

**Fig. 6.**
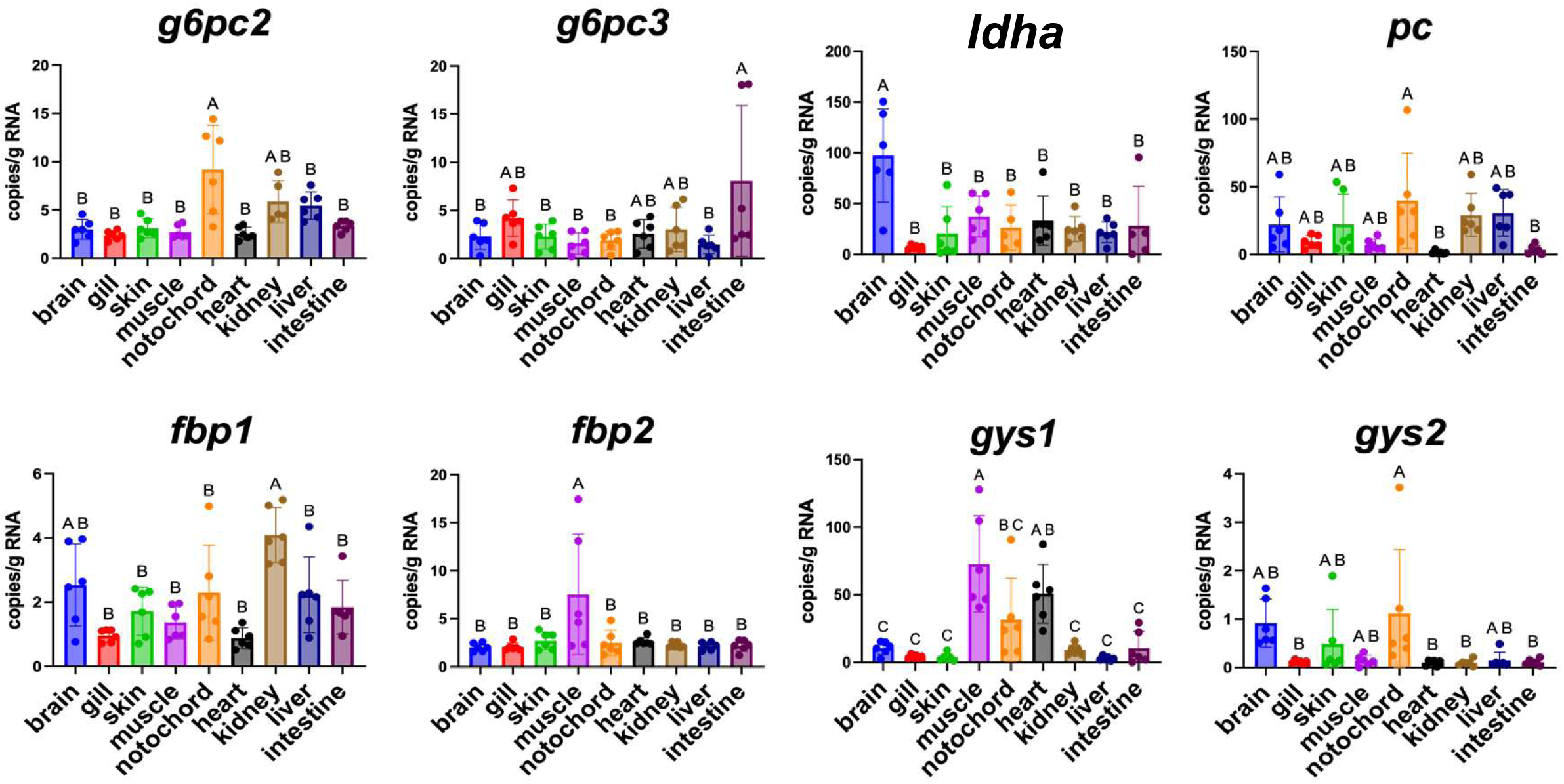
Tissue-specific expression analysis. The expression levels of gluconeogenic genes were measured. Horizontal and vertical axes indicate each tissue and mRNA levels (x 10^9^ copies / g RNA), respectively. Data are presented as mean ± standard error (*n* = 6), and different letters indicate significant differences (*p* < 0.05) between groups. Tests for significance were performed by one-way ANOVA and Tukey’s *post-hoc* test.

### Molecular phylogenetic analysis

In gnathostomes, G6PC1 and G6PC2 formed a well-supported monophyletic clade (support value = 96; Fig. 7, Fig. S1). G6PC3 formed a separate clade containing both cyclostome and gnathostome sequences, with maximal support (support value = 100). Within cyclostomes, G6PC1/2-related sequences formed a distinct clade separate from G6PC3 (support value = 100). Furthermore, two well-supported subclades were identified within the cyclostome-specific G6PC1/2-related clade (support value = 97). These results suggest that the cyclostome-specific duplication event within the G6PC1/2-related lineage occurred independently from the duplication event that generated G6PC1 and G6PC2 in gnathostomes, after the divergence of cyclostomes and gnathostomes.

**Fig. 7.**
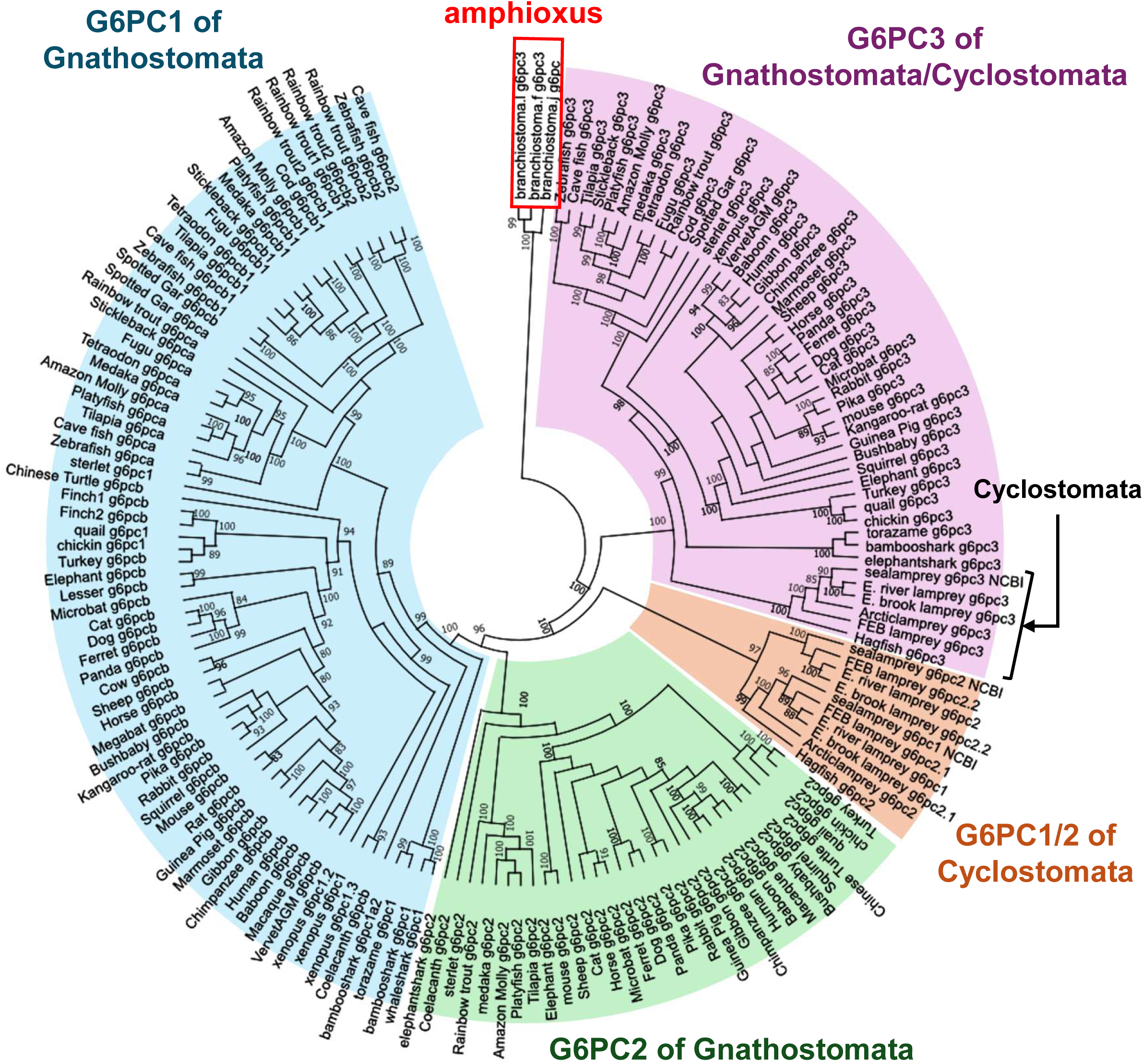
Phylogenetic analysis of Glucose-6-Phosphatase (G6PC) encoding genes. Sequence alignment and tree visualization were performed using MEGA11 (Tamura et al., 2018). The phylogenetic tree was inferred using the maximum-likelihood method implemented in IQ-TREE (Minh et al., 2019). Numbers at nodes indicate SH-aLRT support (%) and ultrafast bootstrap support (%) values, respectively. The tree was rooted using the amphioxus sequence as an outgroup. Detailed support values are provided in Supplementary Fig. S1. All accession numbers obtained from Ensembl and NCBI are listed in Supplementary Table S3.

In gnathostomes, HNF4A and HNF4G formed distinct clades (support value = 85; Fig. 8, Fig. S2), whereas HNF4B formed a separate, strongly supported clade (support value = 100). Cyclostome HNF4 sequences did not cluster within either the gnathostome HNF4B clade or the gnathostome HNF4A/G clade, but instead formed two distinct groups positioned between these lineages (support value = 96, 84). These relationships suggest a complex evolutionary history of the HNF4 gene family, although the timing of duplication events remains unresolved.

**Fig. 8.**
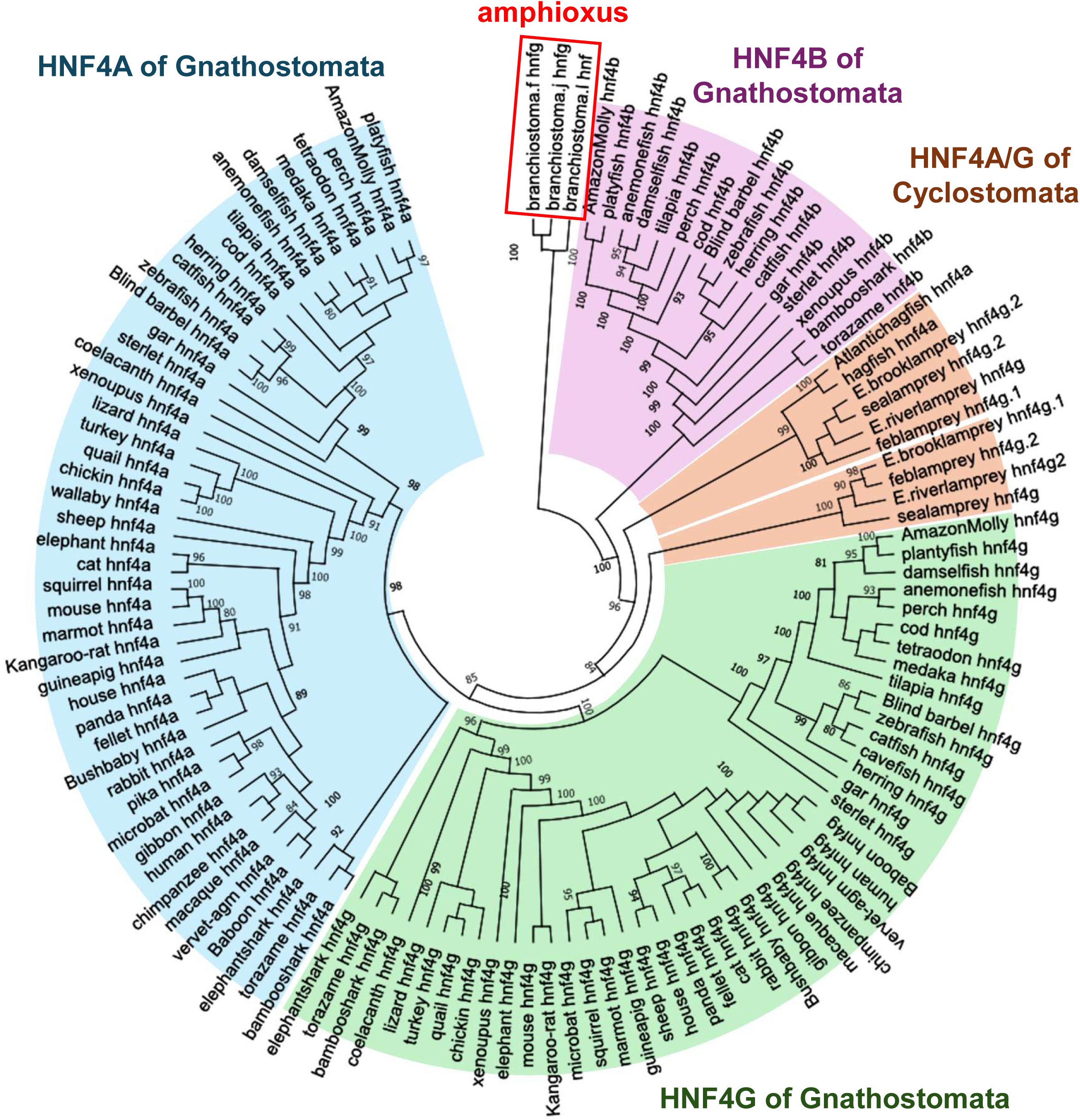
Phylogenetic analysis of Hepatocyte Nuclear Factor 4 (HNF4) encoding genes. Sequence alignment and tree visualization were performed using MEGA11 (Tamura et al., 2018). The phylogenetic tree was inferred using the maximum-likelihood method implemented in IQ-TREE (Minh et al., 2019). Numbers at nodes indicate SH-aLRT support (%) and ultrafast bootstrap support (%) values, respectively. The tree was rooted using the amphioxus sequence as an outgroup. Detailed support values are provided in Supplementary Fig. S2. All accession numbers obtained from Ensembl and NCBI are listed in Supplementary Table S4.

### Motif analysis

In the upstream regions of gnathostome *g6pc*1 genes, motif analysis identified several transcription factor-binding motifs that were conserved across species and enriched relative to background sequences, including motifs for HNF4A, HNF1B, E2A, PU.1, RORA, and PPARA (Supplementary Data 1). Comparison of motif positions among representative vertebrates in which developmental gluconeogenesis has been reported—including fugu, quail, cloudy catshark, western clawed frog, zebrafish, and sterlet—revealed that HNF4A-binding motifs were commonly located within approximately 60–160 bp upstream of the transcription start site (Fig. 9A; Supplementary Data 2, 3). Similarly, HNF4A- and HNF1B-binding motifs were also detected in the upstream regions of gnathostome *fbp1* genes (Supplementary Data 4), and their positions tended to be conserved among species (Supplementary Data 5, 6). In contrast, although similar motif searches were performed for cyclostome *g6pc1/2* and *fbp* upstream regions, no commonly conserved HNF4A-binding motifs were detected, or motif positions were not conserved among species (Fig. 9B; Supplementary Data 5, 7, 8, 9).

**Fig. 9.**
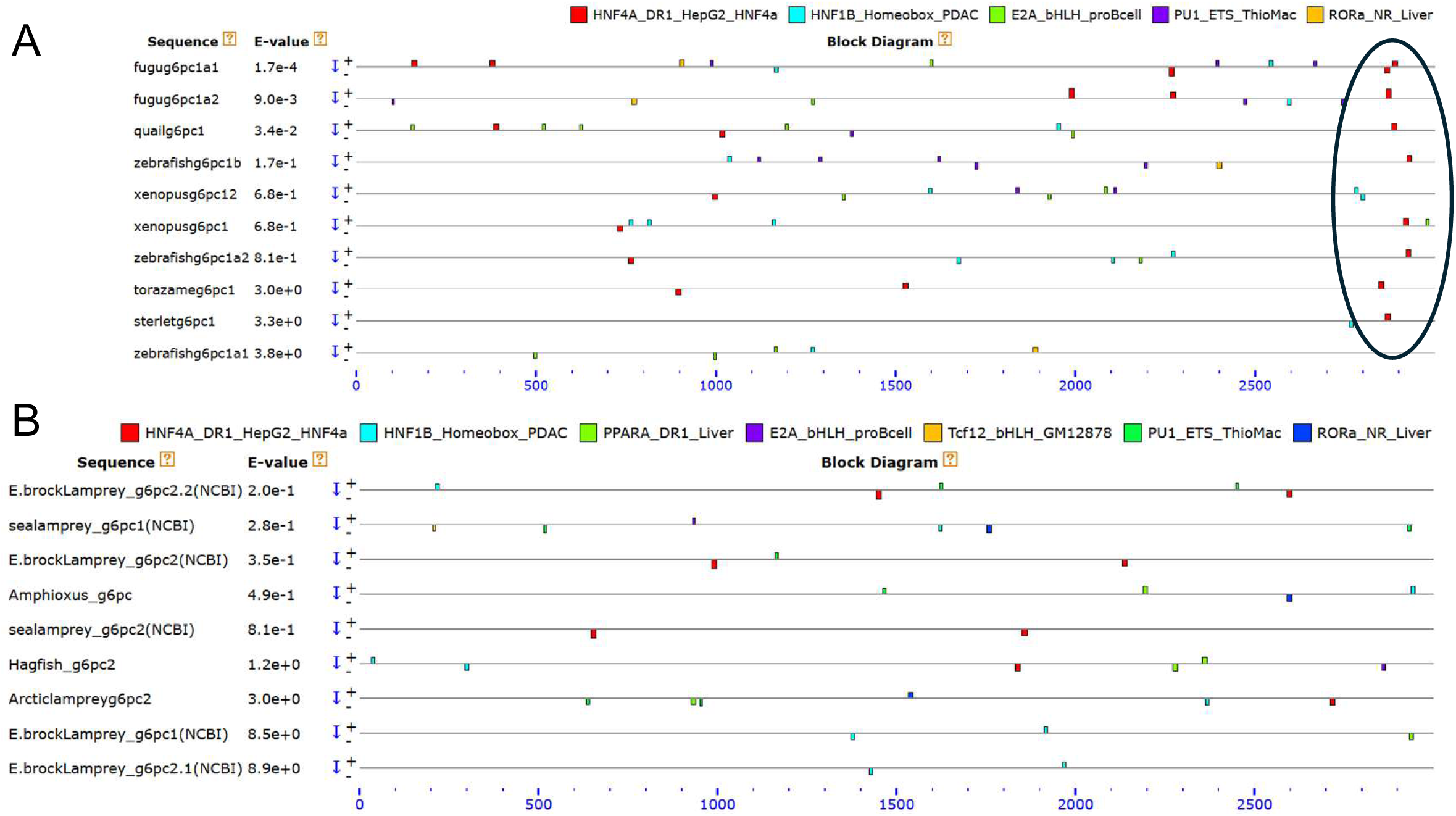
Distribution of conserved motifs in upstream regions of *g6pc* genes. (A) Gnathostomes and (B) cyclostomes. Distribution of conserved motifs in the upstream regions of *g6pc* genes identified by MAST analysis (Bailey and Gribskov, 1988). Colored boxes indicate motif matches and their relative positions within each upstream region. The circle indicates common HNF4α motifs near 5’ ends of gnathostome *g6pc* homologs. Only positions matching motifs with *p*-values < 0.0001 are shown.

A motif analysis of gnathostome *g6pc2* upstream regions identified binding motifs for pancreatic transcription factors, including NeuroD1, NeuroG2, TCF3, and TCF4 (Supplementary Data 10). In contrast, these motifs were not detected in cyclostome *g6pc1/2* upstream regions, suggesting that the pancreas-type transcriptional regulatory mechanism observed in gnathostome *g6pc2* (Dogra et al., 2006) may be absent in cyclostomes (Supplementary Data 11).

## Discussion

Recent studies have revealed that teleosts, cartilaginous fishes, chondrosteans, amphibians, and avian perform gluconeogenesis during development to supply glucose (Shibata et al., 2023; Furukawa et al., 2024; Kodama et al., 2024; Shimizu et al., 2024; 2025. Aoki et al., 2026). These findings suggest that gluconeogenesis during development may be a common and essential phenomenon among most oviparous vertebrates. In this study, we focused on the lamprey, a jawless vertebrate, and examined the similarities and differences of their developmental gluconeogenesis against the jawed vertebrates.

LC/MS-based metabolomic analysis revealed that fertilized eggs of the Arctic lamprey contain little to no glucose immediately after fertilization (Fig. 1). This finding is consistent with observations in zebrafish, cloudy catshark, sturgeon, and western clawed frog, and may represent a common feature among oviparous species (Furukawa et al., 2024; Shimizu et al., 2024; 2025; Aoki et al., 2026), potentially due to the cytotoxicity of glucose (Tchounwou et al., 2014). Glucose levels increased sharply from st. 24 onward, and glycogen showed an increasing trend at earlier stages (sts. 7–8) prior to hatching (Fig. 1), suggesting that the rise in glucose levels during development may partly result from glycogen breakdown. However, glycogen content increased again at st. 27, indicating that lampreys indeed perform gluconeogenesis using yolk-derived substrates during development. To further investigate this possibility, isotope-tracing experiments were conducted using st. 24 prolarvae to analyze actual metabolic activity and substrate utilization dynamics. When the prolarvae were cultured with ^13^C-labeled glycerol, increases in M+3 isotopologues were observed in intermediate metabolites of gluconeogenesis, glycogen metabolism, and the pentose phosphate pathway, including F6P, G6P, G1P, and S7P (Fig. 2). Furthermore, glucose M+3 also increased in samples supplemented with labeled lactate or glycerol (Fig. 2). These M+3 isotopologues most likely represent molecules containing three ^13^C atoms derived from the supplied substrates via the gluconeogenic pathway. Together, these results indicate that the labeled substrates are utilized for glucose production and carbohydrate metabolism in st. 24 lamprey prolarvae, strongly supporting the metabolomic findings. Among the tested substrates, glycerol showed the highest efficiency for glucose production. In adult migrating lampreys, glycerol has also been reported as a major precursor for gluconeogenesis (Savina and Wojtczak, 1977). Although the composition of lamprey yolk remains unclear, glycerol is likely a readily utilizable substrate during the larval stages as well.

Increased glucose levels during lamprey development may be associated with morphological and physiological changes accompanying organogenesis. St.24, at which glucose levels increased markedly in this study, corresponds to the onset of hatching and is characterized by pronounced morphological changes, including elongation of the head and trunk, elevation of the dorsal fin, and development of internal organs (Tahara, 1988; Richardson and Wright, 2003; Richardson et al., 2013). Upon hatching, embryos are released from the relatively stable intra-chorionic space to the external environment, and physiological functions such as muscle activity, fluid circulation, and respiration are likely to become rapidly activated. This shift is likely to increase the demand for glucose as a readily available energy source. The internal structures of st. 24 is characterized by the formation of a tubular heart, differentiation of the dorsal and ventral aortae, and the appearance of blood cells (Tahara, 1988). The establishment of this circulatory system likely promotes the systemic distribution and utilization of glucose. Therefore, the increase in glucose levels measured in the whole embryo may be associated with metabolic reorganization accompanying the establishment of the circulatory system. Furthermore, at st. 24, the forebrain, midbrain, and hindbrain become clearly distinguishable, and the differentiation of the central nervous system progresses markedly, as evidenced by the formation of the pineal organ and the flexure of the cerebral commissure (Tahara, 1988; Richardson and Wright, 2003). Neural tissues are known to exhibit high glucose dependency throughout development (Vannucci et al., 2000; Cacciatore et al., 2022), and glucose is essential for normal brain development (Jensen et al., 2006). These observations suggest that the rapid formation of brain structures is likely accompanied by enhanced glucose metabolism. In addition, the progression of liver bending and the differentiation of the gallbladder and bile ducts after st. 24 coincide with continued increases in glucose levels until st. 27. Taken together, the sharp increase in glucose levels at st. 24 likely reflects an increase in both energy demand and supply associated with the simultaneous progression of hatching, functional maturation of the circulatory and central nervous systems, and the formation of metabolic organs.

To elucidate the mechanisms supporting this stage-specific glucose increase, we analyzed the spatial expression patterns of gluconeogenic genes using *in situ* hybridization in parallel with periodic acid–Schiff (PAS) staining for glycogen signals and H&E staining for general morphology. Consistent with previous studies, H&E staining revealed that the anterior region of the yolk sac becomes epithelialized to form the foregut from st. 28 onward. (Fig. 3). *In situ* hybridization analysis in the lamprey prolarvae revealed the expression of gluconeogenic genes in the epidermis, muscle, and notochord, but not in the extraembryonic yolk cells or endoderm-derived tissues (Fig. 4). This result makes a clear contrast with previous reports in teleosts, cartilaginous fishes, and sturgeons, where gluconeogenesis occurs mainly in endoderm-derived tissues, but not in ectodermal or mesodermal tissues (Furukawa et al., 2024; Kodama et al., 2024; Shimizu et al., 2024; 2025). In adult mammals and teleosts, the liver and kidney are the central gluconeogenic organs (Suarez and Mommsen, 1987). In lampreys, however, although *g6pc* expression levels are higher in the liver than in the brain (Marandel et al., 2017), enzymatic activity is higher in the brain (Savina and Wojtczak, 1977). In addition, muscle has been shown to function as a major gluconeogenic tissue in adult lampreys during spawning migration (Savina and Wojtczak, 1977). Collectively, these findings suggest that, in cyclostomes, not only the liver and kidney but also the brain and muscle may contribute to gluconeogenesis. The present study further suggests that the notochord and epidermis also possess gluconeogenic capacity. Notably, during development, muscle, notochord, and epidermis differentiate earlier than the liver, kidney, and brain (meninges) (Tahara, 1988; Richardson and Wright, 2003; Richardson et al., 2010). For these reasons, it is possible that these tissues play a key role in supplying glucose during early developmental stages.

PAS staining revealed glycogen signals in the epidermis, muscle, and notochord (Fig.5). In many vertebrates, the liver and skeletal muscle functions as the major glycogen storage organ (Adeva-Andany et al., 2016; Chang et al., 2018). In contrast, lampreys reportedly possess relatively low levels of glycogen in the liver (Murat et al., 1979), and total hepatectomy does not cause major changes in carbohydrate metabolism or blood glucose levels (Larsen, 1978). Glycogen accumulation in the lamprey notochord has also been reported previously (Welsch et al., 1991). In addition, notochordal cells in human and rhesus macaque embryos, as well as in sturgeons, contain abundant glycogen in their cytoplasm (Shinohara and Tanaka, 1988; Schmitz, 1988; Wilson and Hendrickx, 1990; Shimizu, 2025). This has been interpreted as indicating that the notochord relies more on glycolysis (anaerobic metabolism) than on mitochondrial respiration (aerobic metabolism). In the present study, not only glycogen metabolism–related genes but also *g6pc* and other gluconeogenic genes were expressed in the larval lamprey notochord. This finding suggests that lamprey notochord serves as a local glucose supply system for surrounding tissues through gluconeogenesis from yolk-derived substrates. Furthermore, the prominent expression of *ldha* in the larval notochord suggests that this tissue is also capable of recycling lactate, the glycolysis end-product, into glucose via gluconeogenesis. These results imply, in this taxon, the notochord is not merely a tissue dependent on glycolytic metabolism but instead represents a metabolically active tissue with the capacity for glucose production.

To investigate whether the distinctive expression patterns of gluconeogenic genes detected by *in situ* hybridization are also conserved in adults, tissue distribution analysis was conducted. The gene *g6pc2* showed high expression levels in the notochord, whereas other gluconeogenic genes did not exhibit tissue-specific expression. In addition, unlike previous reports, high expression levels in the brain or muscle were not observed (Marandel et al., 2017; Savina and Wojtczak, 1977). In the tissue distribution analysis of adult lampreys conducted by Marandel (2017), although the species used was not explicitly stated, it is presumed from the GeneID that the sea lamprey (*Petromyzon marinus*) was examined. Possible reasons for these discrepancies include species differences, as this study used the Arctic lamprey, as well as differences in physiological condition, since the individuals analyzed had initiated upstream migration, ceased feeding, and relied on internally stored energy reserves. Further verification using younger lampreys will be necessary in the future; however, the present results are sufficient to suggest that the notochord of adult lampreys retains the ability to store glycogen and undergo gluconeogenesis.

In the developmental processes of gluconeogenesis in teleosts, cartilaginous fishes, and chondrosteans, the conversion of glucose-6-phosphate to glucose is consistently mediated by *g6pc1* across all taxa, and its expression is localized to the endoderm and endoderm-derived tissues (Furukawa et al., 2024; Kodama et al., 2024; Shimizu et al., 2024; 2025). In cyclostomes, however, *g6pc2* is predominantly expressed instead. Moreover, its expression spans multiple tissues, including the notochord, muscle, and epidermis, and at later developmental stages, it extends to the liver and kidney, covering derivatives of all three germ layers. The G6PC gene family in cyclostomes is notably complex. While G6PC3 is present in all species examined in this study, G6PC1 and G6PC2 show a mixed and inconsistent pattern of occurrence through vertebrate lineages. For example, in the European river lamprey (*Lampetra fluviatilis*), both G6PC1 (Gene ID: 144951465) and G6PC2 (Gene ID: 144951465) are annotated. In contrast, the European brook lamprey (*Lampetra planeri*) possesses two G6PC2-like genes (referred to as 2.1 [Gene ID: 144740865] and 2.2 [Gene ID: 144961665]) but lacks an annotated G6PC1. Similarly, the Far Eastern brook lamprey (*Lethenteron reissneri*) has two G6PC2-like genes (designated here as 2.1 [Gene ID: 133358373] and 2.2 [Gene ID: 133345279]). These inconsistencies suggest that the orthology relationships among cyclostome G6PC genes may not yet be fully resolved, and that current database annotations could include misannotations or lineage-specific naming differences. For the sea lamprey G6PC, Ensembl lists a single G6PC2 (ENSPMAG00000002488.1); however, NCBI database contains both G6PC1 (Gene ID: 116954788) and G6PC2 (Gene ID: 116949542). In this species, making the matters even more complex, the amino acid sequence of G6PC2 in Ensembl (ENSPMAG00000002488.1) is completely identical to that of G6PC1 in NCBI (Gene ID: 116954788), which indicates that the same gene has been annotated under different names in different databases. To clarify the evolutionary relationships among G6PC family genes, a comprehensive molecular phylogenetic analysis was performed using the maximum-likelihood method. The resulting tree strongly supported the orthology of cyclostome and gnathostome G6PC3 genes, as all G6PC3 sequences formed a single well-supported clade (support value = 100; Fig. 7, Fig. S1). In contrast, cyclostome G6PC1/2-related sequences formed a distinct clade separate from gnathostome G6PC1 and G6PC2 (support value = 100). Furthermore, two subclades were identified within the cyclostome-specific G6PC1/2-related lineage. These results suggest that the duplication event that generated G6PC1 and G6PC2 in gnathostomes occurred independently from the duplication event within the cyclostome G6PC1/2-related lineage, after the divergence of cyclostomes and gnathostomes. Thus, genes currently annotated as G6PC1/2 in cyclostomes may not represent direct orthologs of gnathostome G6PC1 and G6PC2 despite their similar nomenclature. Furthermore, for the Arctic lamprey used in this study, only a single G6PC gene is registered in each database, and no additional copies were detected through BLAST searches. To clone potential unidentified G6PC isoforms, primers were designed based on sequences conserved among G6PC2.2 of the sea lamprey, European brook lamprey, European river lamprey, and Far Eastern brook lamprey, and PCR was performed (forward 1, 5′-AATGTCCACTTAGATCGGTAAGG -3′; reverse 1, 5′-AACCAGAAGAGCAGCCAGAG -3′; forward 2, 5′-CTCTGGCTGCTCTTCTGGTT -3′; reverse 2, 5′-GGCCATCAGGACCCCTTC -3′); however, no amplification products were obtained.

Despite the challenges of complex gene database and isoform composition, it remains plausible that cyclostome G6PC1/2 (following Ensembl nomenclature, which is used hereafter) are regulated by transcription factors distinct from those controlling gnathostome G6PC1, leading to differences in expression patterns. To investigate this, HOMER was first used to identify transcription factor binding motifs commonly present in the upstream regions of gnathostome *g6pc1/2* as well as *fbp1*, another key gene involved in gluconeogenesis. In *g6pc1* and *fbp1*, binding motifs for transcription factors such as HNF4A and HNF1B—known to function in hepatocytes and to induce the expression of gluconeogenic genes (Lau et al., 2018)—were commonly identified (Supplementary Data1, 4). In contrast, *g6pc2* shared motifs for pancreas-specific transcription factors, including NeuroD1 and NeuroG2 (Anderson et al., 2009) (Supplementary Data 10). By using MEME to estimate the positions of binding motifs on the sequences, we found that in most gnathostome *g6pc1* analyzed, the HNF4A binding motifs are conserved in the close 5’ flanking regions (Fig. 9A; Supplementary Data 2, 3). A similar trend was also observed in *fbp1* (Supplementary Data 6, 7). However, these motifs were not detected in the sequences of lamprey and amphioxus, or their positions were not conserved (Fig. 9B; Supplementary Data 7, 8, 11). In Ensembl, only a single HNF4-related gene is annotated in hagfish, registered as HNF4A, with no paralogs identified. In contrast, in lampreys (both sea lamprey and Arctic lamprey), only a single gene is annotated as HNF4G in both Ensembl and NCBI, and similarly, no paralogs are present. Considering the possibility of misannotation, we constructed a molecular phylogenetic tree of HNF4A/B/G (Fig. 8, Fig. S2). Consistent with previous studies (Bertrand et al., 2004), gnathostome HNF4B formed a distinct clade separate from HNF4A and HNF4G. Within gnathostomes, HNF4A and HNF4G formed well-defined sister clades. In contrast, cyclostome HNF4 sequences did not cluster within either the gnathostome HNF4B clade or the gnathostome HNF4A/G clade, but instead formed two independent groups positioned between these lineages. These results indicate that the evolutionary relationships among cyclostome and gnathostome HNF4 genes are more complex than suggested by current gene annotations. Therefore, the assignment of cyclostome HNF4 genes as HNF4A, HNF4B, or HNF4G orthologs should be interpreted with caution. Similar to the situation observed for G6PC1/2-related genes, the current nomenclature of cyclostome HNF4 genes may not accurately reflect their evolutionary relationships with gnathostome HNF4 paralogs. In a previous study (Jiang et al., 2021), western blotting and immunostaining were performed using three different monoclonal antibodies against HNF4A of several gnathostomes and Arctic lamprey; however, among these animals, lamprey alone was negative for all antibodies. This observation is consistent with the results of the molecular phylogenetic tree for HNF4 constructed in this study. The function of HNF4 in cyclostomes has not yet been elucidated. It is generally considered that all HNF4 paralogs share the same DNA-binding motif (Fang et al., 2012), and at least in mammals, HNF4A and HNF4G have been reported to exhibit common sequence specificity (Chen et al., 2019). To predict potential similarities in binding motifs of lamprey HNF4G, the DNA-binding domains of zebrafish HNF4A/B/G and lamprey HNF4G were predicted using InterPro, and their amino acid sequences were compared using CLUSTAL 2.1 multiple sequence alignment. The DNA-binding domains were highly conserved among all examined HNF4 proteins, with most residues shared across the 75–76 aa region (Supplementary Data 12). Consistently, guide-tree branch lengths estimated by CLUSTAL were relatively low (0.026–0.067). These results suggest that the HNF4 family, including HNF4B, is likely to recognize a common DNA-binding motif across species. Therefore, the absence of HNF4-binding motifs in cyclostome *g6pc* and *fbp* suggests that these genes may not be directly regulated by HNF4. Alternative transcription factors may be responsible for regulating gluconeogenesis, potentially contributing to differences in the expression of gluconeogenic genes between gnathostomes and cyclostomes. As the genomes of more cyclostomes are sequenced in the future, common transcription factor-binding motifs will be found in cyclostome gluconeogenic genes, leading to a better understanding of their regulatory mechanisms.

This study revealed that gluconeogenic genes are expressed across all three germ layers in lamprey. These findings suggest multiple possible scenarios regarding the evolutionary origin of gluconeogenesis and the establishment of its tissue distribution in vertebrates. First, developmental gluconeogenesis may originally have been restricted to a limited set of tissues, with subsequent expansion of its expression domains in the lamprey lineage. Alternatively, the common ancestor may already have exhibited widespread gluconeogenic activity across multiple embryonic tissues, with this activity subsequently becoming more spatially restricted during gnathostome evolution. These two hypotheses are not mutually exclusive at present, and further comparative developmental analyses across a broader range of taxa, including amphioxus and hagfish, will be necessary to clarify the evolutionary history of developmental gluconeogenesis.

## Acknowledgment

We are grateful to the laboratory members at AORI for their support in maintaining lampreys, with special thanks to Mr. Kanata Suzuki and Mr. Keigo Nagasaka for their exceptional assistance.

## Funding

This work was supported partly by Grant-in-Aid for Exploratory Research (no. 21K19276) and Grant-in-Aid for scientific Research (B) (no. 22H02426) from JSPS to Fumiya Furukawa.

## Notes

### Competing Interest Statement

The authors have declared no competing interest.

## References

Aoki M, Tsuchida A, Tamura K, Baba O, Yoshitake K, Furukawa F. 2026. Gluconeogenesis and glycogen metabolism in the epidermis and endoderm of Xenopus tropicalis embryos and larvae. BioRxiv 10.64898/2026.05.08.723674

Adeva-Andany MM, González-Lucán M, Donapetry-García C, Fernández-Fernández C, Ameneiros-Rodríguez E. 2016. Glycogen metabolism in humans. BBA Clin 5; 85–100. 10.1016/j.bbacli.2016.02.001.

Anderson KR, Torres CA, Solomon K, Becker TC, Newgard CB, Wright CV, Hagman J, Sussel L. 2009. Cooperative Transcriptional Regulation of the Essential Pancreatic Islet Gene *NeuroD1* (*Beta2*) by Nkx2.2 and Neurogenin 3. J Biol Chem 284 (45): 31236–31248. 10.1074/jbc.M109.048694

Bailey TL, Grant CE, Johnson J, Noble WS. 2015. The MEME Suite. Nucleic Acids Res 43 (W1): W39–W49. 10.1093/nar/gkv416.

Bailey TL, Gribskov M. Combining evidence using p-values: application to sequence homology searches. Bioinformatics 14 (1): 48–54. 10.1093/bioinformatics/14.1.48.

Bertrand S, Brunet FG, Escriva H, Parmentier G, Laudet V, Robinson-Rechavi M. 2004. Evolutionary Genomics of Nuclear Receptors: From Twenty-Five Ancestral Genes to Derived Endocrine Systems. Mol Biol Evol 21 (10): 1923–37. 10.1093/molbev/msh200.

Cacciatore M, Grasso EA, Tripodi R, Chiarelli F. 2022. Impact of Glucose Metabolism on the Developing Brain. Front Endocrinol 13 (December): 1047545. 10.3389/fendo.2022.1047545.

Chang CH, Huang JJ, Yeh CY, Tang CH, Hwang LY, Lee TH.2018. Salinity Effects on Strategies of Glycogen Utilization in Livers of Euryhaline Milkfish (*Chanos chanos*) under Hypothermal Stress. Front Physiol 9: 81. 10.3389/fphys.2018.00081

Chea HK, Wright CV, Swalla BJ. 2005. Nodal Signaling and the Evolution of Deuterostome Gastrulation. Dev Dyn 234 (2): 269–78. 10.1002/dvdy.20549.

Chen L, Toke NH, Luo S, Vasoya RP, Fullem RL, Parthasarathy A, Perekatt AO, Verzi MP. 2019. A Reinforcing HNF4–SMAD4 Feed-Forward Module Stabilizes Enterocyte Identity. Nat Genet 51 (5): 777–85. 10.1038/s41588-019-0384-0.

Grant CE, Bailey TL, Noble WS. 2011. FIMO: scanning for occurrences of a given motif. Bioinformatics 27(7): 1017–1018. 10.1093/bioinformatics/btr064

Fang B, Mane-Padros D, Bolotin E, Jiang T, Sladek FM. 2012. Identification of a Binding Motif Specific to HNF4 by Comparative Analysis of Multiple Nuclear Receptors. Nucleic Acids Res 40 (12): 5343–56. 10.1093/nar/gks190.

Foster GD, Youson JH, Moon TW. 1993. Carbohydrate Metabolism in the Brain of the Adult Lamprey. J Exp Zool 267 (1): 27–32. 10.1002/jez.1402670105.

Furukawa F, Aoyagi A, Sano K, Sameshima K, Goto M, Tseng YC, Ikeda D, Lin CC, Uchida K, Okumura S, Yasumoto K, Jimbo M, Hwang PP. 2024. Gluconeogenesis in the Extraembryonic Yolk Syncytial Layer of the Zebrafish Embryo. PNAS Nexus 3 (4): pgae125. 10.1093/pnasnexus/pgae125.

Furukawa F, Tseng YC, Liu ST, Chou YL, Lin CC, Sung PH, Uchida K, Lin LY, Hwang PP. 2015. Induction of Phosphoenolpyruvate Carboxykinase (PEPCK) during Acute Acidosis and Its Role in Acid Secretion by V-ATPase-Expressing Ionocytes. Int J Bio. Sci 11 (6): 712–25. 10.7150/ijbs.11827.

Dogra RS, Vaidyanathan P, Prabakar KR, Marshall KE, Hutton JC, Pugliese A. 2006. Alternative splicing of G6PC2, the gene coding for the islet-specific glucose-6-phosphatase catalytic subunit-related protein (IGRP), results in differential expression in human thymus and spleen compared with pancreas. Diabetologia 49: 953–957. 10.1007/s00125-006-0185-8

Hamlett WC and Wourms JP. 1984. Ultrastructure of the Pre-Implantation Shark Yolk Sac Placenta. Tissue and Cell 16 (4): 613–25. 10.1016/0040-8166(84)90035-1.

Hamor T, Garside ET. 1977. Quantitative Composition of the Fertilized Ovum and Constituent Parts in the Atlantic Salmon *Salmo Salar* L. Can J Zool 55 (10): 1650–55. 10.1139/z77-214.

Harris JM, Esain V, Frechette GM, Harris LJ, Cox AG, Cortes M, Garnaas MK, Carroll KJ, Cutting CC, Khan T, Elks PM, Renshaw SA, Dickinson BC, Chang CJ, Murphy MP, Paw BH, Vander Heiden MG, Goessling W, North TE. 2013. Glucose Metabolism Impacts the Spatiotemporal Onset and Magnitude of HSC Induction in Vivo. Blood 121 (13): 2483–93. 10.1182/blood-2012-12-471201.

Heinz S, Benner C, Spann N, Bertolino E, Lin YC, Laslo P, Cheng JX, Murre C, Singh H, Glass CK. 2010. Simple combinations of lineage-determining transcription factors prime cis-regulatory elements required for macrophage and B cell identities. Mol Cell 38(4):576–589. https://linkinghub.elsevier.com/retrieve/pii/S1097276510003667

Hoang DT, Chernomor O, von Haeseler A, Minh BQ, Vinh LS. 2018. UFBoot2: Improving the Ultrafast Bootstrap Approximation. Mol Biol Evol 35(2): 518–522. 10.1093/molbev/msx281

Honda Y, Ogawa N, Wong MKS, Tokunaga K, Kuraku S, Hyodo S, Takagi W. 2022. Molecular Mechanism of Nutrient Uptake in Developing Embryos of Oviparous Cloudy Catshark (*Scyliorhinus Torazame)*. PLOS ONE 17 (3): e0265428. 10.1371/journal.pone.0265428.

Jensen PJ, Gitlin JD, Carayannopoulos MO. 2006. GLUT1 Deficiency Links Nutrient Availability and Apoptosis during Embryonic Development. J Biol Chem 281 (19): 13382–87. 10.1074/jbc.M601881200.

Jiang S, Tanaka T, Yagami R, Hasegawa G, Umezu H, Fujiyoshi Y, Kodama T, Naito M, Ajioka Y. 2021. Immunohistochemical Detection of Hepatocyte Nuclear Factor□4α in Vertebrates. Microsc Res Tech 84 (12): 2906–14. 10.1002/jemt.23848.

Kalyaanamoorthy S, Minh BQ, Wong TKF, von Haeseler A, Jermiin LS. 2017. ModelFinder: Fast Model Selection for Accurate Phylogenetic Estimates. Nat Methods 14(6): 587–589. 10.1038/nmeth.4285

Kimmel CB and Law RD. 1985. Cell Lineage of Zebrafish Blastomeres. Dev Biol 108 (1): 86–93. 10.1016/0012-1606(85)90011-9.

Knox D, Walton MJ, and Cowey CB. 1980. Distribution of Enzymes of Glycolysis and Gluconeogenesis in Fish Tissues. Mar Biol 56 (1): 7–10. 10.1007/BF00390588.

Knpffer C. 1890. Die Entwicklung von *Petromyzon Planeri*. Arch mikrosk Anat 35 (1): 469–558. 10.1007/BF02955888.

Kodama A, Watanabe S, Kayanuma I, Sasaki A, Kurokawa D, Baba O, Jimbo M, Furukawa F. 2024. Gluconeogenesis during Development of the Grass Puffer (*Takifugu Niphobles*). Comp Biochem Physiol A Mol Integr Physiol 295 (September): 111663. 10.1016/j.cbpa.2024.111663.

Kuraku S, Ota KG, Kuratani S. 2009. Jawless fishes (Cyclostomata). Kumar S, Hedges B. (Eds.). Time tree of Life. Oxford Univ. Press. 10.1093/oso/9780199535033.003.0040

Larsen LO. 1978. Subtotal Hepatectomy in Intact or Hypophysectomized River Lampreys (*Lampetra Fluviatilis L*.): Effects on Regeneration, Blood Glucose Regulation, and Vitellogenesis. Gen Comp Endocrinol 35 (3): 197–204. 10.1016/0016-6480(78)90063-1.

Lau HH, Ng NHJ, Loo LSW, Jasmen JB, Teo AKK. 2018. The molecular functions of hepatocyte nuclear factors – In and beyond the liver. J Hepato 68 (5): 1033–1048. 10.1016/j.jhep.2017.11.026

Leache AD. 2010. The Timetree of Life. S. Blair Hedges and Sudhir Kumar, Editors. Integr Comp Biol 50 (1): 141–42. 10.1093/icb/icp110.

Lechenault H, Wrisez F, Mellinger J. 1993. Yolk Utilization in *Scyliorhinus Canicula*, an Oviparous Dogfish. In The Reproduction and Development of Sharks, Skates, Rays and Ratfishes, edited by Leo SD and John PW, vol. 14, edited by Eugene KB. Developments in Environmental Biology of Fishes. Springer Netherlands. 10.1007/978-94-017-3450-9_22.

Li J, Han Y, Ma Q, Liu H, Pang Y, Li Q. 2019. Early Development of Lamprey *Lampetra Japonica* (Martens, 1868). Aquac Res 50 (5): 1501–14. 10.1111/are.14026.

Marandel L, Panserat S, Plagnes-Juan E, Arbenoits E, Soengas JL, Bobe J. 2017. Evolutionary History of Glucose-6-Phosphatase Encoding Genes in Vertebrate Lineages: Towards a Better Understanding of the Functions of Multiple Duplicates. BMC Genom 18 (1): 342. 10.1186/s12864-017-3727-1.

Menger MD, Steiner D, Messmer K. 1992. Microvascular Ischemia-Reperfusion Injury in Striated Muscle: Significance of ‘No Reflow.’ Am J Physiol 263 (6 Pt 2): H1892–1900. 10.1152/ajpheart.1992.263.6.H1892.

Minh BQ, Schmidt HA, Chernomor O, Schrempf D, Woodhams MD, von Haeseler A, Lanfear R. 2020. IQ-TREE 2: New Models and Efficient Methods for Phylogenetic Inference in the Genomic Era. Mol Biol Evol 37(5): 1530–1534. 10.1093/molbev/msaa015

Murakami T, Wakamatsu E, Tamahashi N, Takahashi T. 1985. The Functional Significance of Human Notochord in the Development of Vertebral Column. An Electron Microscopic Study. Tohoku J Exp Med 146 (3): 321–36. 10.1620/tjem.146.321.

Murat JC, Plisetskaya EM, Soltitskaya LP. 1979. Glucose 6-Phosphatase Activity in Kidney of the River Lamprey (*Lampetra Fluviatilis L*.). Gen Comp Endocrinol 39 (1): 115–17. 10.1016/0016-6480(79)90197-7.

Ninhaus-Silveira A, Foresti F, de Azevedo A, Agostinho CA, Veríssimo-Silveira R. 2007. Structural and Ultrastructural Characteristics of the Yolk Syncytial Layer in *Prochilodus Lineatus* (Valenciennes, 1836) (Teleostei; Prochilodontidae). Zygote 15 (3): 267–71. 10.1017/S0967199407004261.

Palmer JF, Slack C. 1970. Some bio-electric parameters of early Xenopus embryos. J Embryol Exp Morphol 24 (3): 535–553. PMID: 4992748

Richardson MK, Admiraal J, Wright GM. 2010. Developmental Anatomy of Lampreys. Biol Rev 85 (1): 1–33. 10.1111/j.1469-185X.2009.00092.x.

Richardson MK, and Wright GM. 2003. Developmental Transformations in a Normal Series of Embryos of the Sea Lamprey *Petromyzon Marinus* (Linnaeus). J Morphol 257 (3): 348–63. 10.1002/jmor.10119.

Savina MV, Wojtczak AB. 1977. Enzymes of Gluconeogenesis and the Synthesis of Glycogen from Glycerol in Various Organs of the Lamprey (*Lampetra Fluviatilis*). *Comp biochem. physiol.*, B. Comp. biochem 57 (3): 185–90. 10.1016/0305-0491(77)90141-9.

Schmitz RJ. 1998. Comparative Ultrastructure of the Cellular Components of the Unconstricted Notochord in the Sturgeon and the Lungfish. J Morphol 236 (2): 75–104. 10.1002/(SICI)1097-4687(199805)236:2%3C75::AID-JMOR1%3E3.0.CO;2-N.

Shah MA, Xie X, Rodina M, Stundl J, Braasch I, SLindelka R, Rzepkowska M, Saito T, PsLenicLka M. 2024. Sturgeon Gut Development: A Unique Yolk Utilization Strategy among Vertebrates. Front Cell Dev Biol 12 (May): 1358702. 10.3389/fcell.2024.1358702.

Shaughnessy CA., and McCormick SD. 2021. 11-Deoxycortisol Is a Stress Responsive and Gluconeogenic Hormone in a Jawless Vertebrate, the Sea Lamprey (*Petromyzon Marinus*). J Exp Biol 224 (11): jeb241943. 10.1242/jeb.241943.

Shibata M, Iwasawa A, Yayota M. 2023. Gluconeogenesis in the Yolk Sac Membrane: Enzyme Activity, Gene Expression, and Metabolites During Layer Chicken Development. J poult sci 60 (2). 10.2141/jpsa.2023020.

Shimizu M, Matsuzaki T, Kobayashi K, Aoyagi A, Kaneko M, Nagasawa T, Hiraoka K, Furukawa F. 2025. Gluconeogenesis and Glycogen Metabolism during Development in Sterlet Acipenser Ruthenus and Bester. Fish sci 91 (5): 881–90. 10.1007/s12562-025-01893-3.

Shimizu M, Takagi W, Sakai Y, Kayanuma I, Furukawa F. 2024. Gluconeogenesis in the Yolk Syncytial Layer□like Tissue of Cloudy Catshark ( *Scyliorhinus Torazame*). Physiol Rep 12 (11): e16088. 10.14814/phy2.16088.

Shinohara H, and Tanaka O. 1988. Development of the Notochord in Human Embryos: Ultrastructural, Histochemical, and Immunohistochemical Studies. Anat rec 220 (2): 171–78. 10.1002/ar.1092200208.

Soukup V, Horácek I, Cerny R. 2013. Development and Evolution of the Vertebrate Primary Mouth. J Anat 222 (1): 79–99. 10.1111/j.1469-7580.2012.01540.x.

Suarez RK, and Mommsen TP. 1987. Gluconeogenesis in Teleost Fishes. Can J Zool 65 (8): 1869–82. 10.1139/z87-287.

Tahara Y. 1988. Normal stages of development in the lamprey, *Lampetra reissneri* (Dybowski). Zoo sci 5 (1): 109–118. 10.34425/zs000449

Takeuchi M, Takahashi M, Okabe M, Aizawa S. 2009. Germ Layer Patterning in Bichir and Lamprey; an Insight into Its Evolution in Vertebrates. Dev Biol 332 (1): 90–102. 10.1016/j.ydbio.2009.05.543.

Takezaki N, Figueroa F, Zaleska-Rutczynska Z, Klein J. 2003. Molecular Phylogeny of Early Vertebrates: Monophyly of the Agnathans as Revealed by Sequences of 35 Genes. Mol Biol Evol 20 (2): 287–92. 10.1093/molbev/msg040.

Tamura K, Stecher G, Kumar S. 2021. MEGA11: Molecular Evolutionary Genetics Analysis Version 11. Mol Biol Evol 38(7): 3022–3027. 10.1093/molbev/msab120

Tchounwou CK, Yedjou CG, Farah I, Tchounwou PB. 2014. d-Glucose-Induced Cytotoxic, Genotoxic, and Apoptotic Effects on Human Breast Adenocarcinoma (MCF-7) Cells. J Cancer Sci Ther 6: 156–160. doi: 10.4172/1948-5956.1000265.

Vannucci RC and Vannucci SJ. 2000. Glucose Metabolism in the Developing Brain. Semin Perinatol 24 (2): 107–15. 10.1053/sp.2000.6361.

Weinrauch AM, Edwards SL, Goss GG, eds. 2015. Anatomy of the Pacific Hagfish (*Eptatretus Stoutii*). In Hagfish Biology, 0 ed. CRC Press. 10.1201/b18935-5.

Welsch U, Erlinger R, Potter IC. 1991. Proteoglycans in the Notochord Sheath of Lampreys. Acta Histochem 91 (1): 59–65. 10.1016/S0065-1281(11)80295-3.

Wilson DB and Hendrickx AG. 1990. Cytochemical Analysis of the Notochord in Early Rhesus Monkey Embryos. ANAT REC 228 (4): 431–36. 10.1002/ar.1092280409.

