## Supplementary Data 1 for "Evolutionary insights into glucose production in vertebrate development: new findings from Arctic lamprey (*Lethenteron camtschaticum*)": Suplementary Data1_G_g6pc1_motif.html

g6pc1.2\_motif - Homer Known Motif Enrichment Results


### Homer Known Motif Enrichment Results (g6pc1.2\_motif)

Homer *de novo* Motif Results  
Gene Ontology Enrichment Results  
Known Motif Enrichment Results (txt file)  
Total Target Sequences = 60, Total Background Sequences = 285

|  |  |  |  |  |  |  |  |  |  |  |  |
| --- | --- | --- | --- | --- | --- | --- | --- | --- | --- | --- | --- |
| Rank | Motif | Name | P-value | log P-pvalue | q-value (Benjamini) | # Target Sequences with Motif | % of Targets Sequences with Motif | # Background Sequences with Motif | % of Background Sequences with Motif | Motif File | SVG |
| 1 | G C T A G C A T A C G T A C G T A G C T G A T C G A C T G A C T A G C T A C G T A C G T A G C T | RLR1?/SacCer-Promoters/Homer | 1e-21 | -4.970e+01 | 0.0000 | 53.0 | 88.33% | 63.2 | 22.16% | motif file (matrix) | svg |
| 2 | A C G T A C G T A C G T A C G T A C G T A C G T A C G T A C G T A C G T A C G T | VRN1(ABI3VP1)/col-VRN1-DAP-Seq(GSE60143)/Homer | 1e-18 | -4.250e+01 | 0.0000 | 31.0 | 51.67% | 10.1 | 3.55% | motif file (matrix) | svg |
| 3 | G T C A T G C A T G C A G C T A C G T A G C T A G C T A G C T A | REM19(REM)/colamp-REM19-DAP-Seq(GSE60143)/Homer | 1e-15 | -3.595e+01 | 0.0000 | 56.0 | 93.33% | 109.0 | 38.25% | motif file (matrix) | svg |
| 4 | G T A C G C T A T C A G C T G A C T A G C A T G A G C T G A T C T G C A T C G A C T G A A C T G C A G T A G T C G A T C G C T A | HNF4a(NR),DR1/HepG2-HNF4a-ChIP-Seq(GSE25021)/Homer | 1e-15 | -3.496e+01 | 0.0000 | 50.0 | 83.33% | 78.1 | 27.39% | motif file (matrix) | svg |
| 5 | G T A C C T G A T A G C C G T A G C T A T C G A T G C A T G A C C T A G G T C A A G T C C G T A C T G A C T G A C G T A | At1g14580(C2H2)/colamp-At1g14580-DAP-Seq(GSE60143)/Homer | 1e-13 | -3.102e+01 | 0.0000 | 32.0 | 53.33% | 24.4 | 8.56% | motif file (matrix) | svg |
| 6 | C T A G C T A G T C G A C T A G C G T A A T C G T C G A A C T G C T G A T C G A C T G A T A C G | FRS9(ND)/col-FRS9-DAP-Seq(GSE60143)/Homer | 1e-12 | -2.942e+01 | 0.0000 | 23.0 | 38.33% | 8.0 | 2.82% | motif file (matrix) | svg |
| 7 | G A T C G A T C A G T C G T A C C G A T G T A C G T A C A G T C A G T C A G T C G C T A G A T C | ZNF148(Zf)/MDAMB231-ZNF148-ChIP-Seq(GSE147020)/Homer | 1e-11 | -2.757e+01 | 0.0000 | 30.0 | 50.00% | 24.1 | 8.46% | motif file (matrix) | svg |
| 8 | T G A C G C T A T G A C C G T A T C A G G A T C C G T A C A T G C A T G C T A G C T A G C T A G | Unknown-ESC-element(?)/mES-Nanog-ChIP-Seq(GSE11724)/Homer | 1e-11 | -2.693e+01 | 0.0000 | 40.0 | 66.67% | 55.8 | 19.56% | motif file (matrix) | svg |
| 9 | C T A G A C G T G C A T T C G A G T C A G C A T A T C G C G T A C A G T A G C T C T G A T G C A | HNF1b(Homeobox)/PDAC-HNF1B-ChIP-Seq(GSE64557)/Homer | 1e-11 | -2.573e+01 | 0.0000 | 46.0 | 76.67% | 82.4 | 28.91% | motif file (matrix) | svg |
| 10 | G A T C G A T C G T C A C G T A A C G T T A C G C G T A A C G T G C A T C T A G | ATHB18(Homeobox)/colamp-ATHB18-DAP-Seq(GSE60143)/Homer | 1e-10 | -2.484e+01 | 0.0000 | 50.0 | 83.33% | 104.6 | 36.69% | motif file (matrix) | svg |
| 11 | G C A T A G C T A C G T A C G T A C T G A C G T G A T C A C G T A C G T A G C T C G A T G C A T A G T C G A C T C A G T | IDD5(C2H2)/colamp-IDD5-DAP-Seq(GSE60143)/Homer | 1e-9 | -2.216e+01 | 0.0000 | 46.0 | 76.67% | 92.9 | 32.58% | motif file (matrix) | svg |
| 12 | T C G A T C G A C T G A C G T A A C T G A T G C A C G T A G T C | Lola-I(Zf)/Embryo-LolaI-ChIP-Seq(GSE200870)/Homer | 1e-9 | -2.131e+01 | 0.0000 | 50.0 | 83.33% | 115.6 | 40.55% | motif file (matrix) | svg |
| 13 | T C A G C A T G C A T G A C T G A C T G A G C T A C T G A C G T A C T G C A G T A T G C A G T C | KLF10(Zf)/HEK293-KLF10.GFP-ChIP-Seq(GSE58341)/Homer | 1e-9 | -2.114e+01 | 0.0000 | 33.0 | 55.00% | 44.0 | 15.45% | motif file (matrix) | svg |
| 14 | T G A C G T A C C G T A A C T G T G A C C G A T A C T G A T C G A G C T T A C G T C G A T A G C G T A C C G T A A T C G T G A C G C A T A C T G A C T G A T G C | Twist(bHLH)/HMLE-TWIST1-ChIP-Seq(Chang\_et\_al)/Homer | 1e-8 | -2.040e+01 | 0.0000 | 19.0 | 31.67% | 10.5 | 3.70% | motif file (matrix) | svg |
| 15 | G A T C G A C T G A C T A C G T A G T C A C G T A G T C A C G T A G T C A C G T A G T C A C G T G T A C C G A T G T C A | BPC6(BBRBPC)/col-BPC6-DAP-Seq(GSE60143)/Homer | 1e-8 | -2.006e+01 | 0.0000 | 11.0 | 18.33% | 0.0 | 0.00% | motif file (matrix) | svg |
| 16 | C G A T C T A G A C G T A C G T A C G T C G T A A G C T C G A T A G C T C G T A C T A G T A G C | FoxD3(forkhead)/ZebrafishEmbryo-Foxd3.biotin-ChIP-seq(GSE106676)/Homer | 1e-8 | -2.004e+01 | 0.0000 | 53.0 | 88.33% | 137.8 | 48.36% | motif file (matrix) | svg |
| 17 | C T G A T C A G A G T C C G T A A T C G A T G C C G A T A C T G A G T C G A C T A T C G A G T C | MyoD(bHLH)/Myotube-MyoD-ChIP-Seq(GSE21614)/Homer | 1e-8 | -1.982e+01 | 0.0000 | 42.0 | 70.00% | 81.0 | 28.43% | motif file (matrix) | svg |
| 18 | A T G C G A T C C G T A C G T A A C G T A T G C C G T A A C G T C G A T C T A G A C T G G T A C | HAT1(Homeobox)/col-HAT1-DAP-Seq(GSE60143)/Homer | 1e-8 | -1.959e+01 | 0.0000 | 46.0 | 76.67% | 100.5 | 35.26% | motif file (matrix) | svg |
| 19 | T G A C G T C A A G T C G C T A C T A G A G T C C T G A C A T G A C T G C T A G T A C G T C A G | Zic2(Zf)/ESC-Zic2-ChIP-Seq(SRP197560)/Homer | 1e-8 | -1.956e+01 | 0.0000 | 34.0 | 56.67% | 51.4 | 18.04% | motif file (matrix) | svg |
| 20 | C G T A T A C G T C G A A C T G A C T G C G T A C G T A T A C G A G C T T A C G | PU.1(ETS)/ThioMac-PU.1-ChIP-Seq(GSE21512)/Homer | 1e-8 | -1.920e+01 | 0.0000 | 36.0 | 60.00% | 59.4 | 20.84% | motif file (matrix) | svg |
| 21 | T C G A T C G A A G T C C G T A C T A G T A G C A C G T A C T G | MyoG(bHLH)/C2C12-MyoG-ChIP-Seq(GSE36024)/Homer | 1e-8 | -1.881e+01 | 0.0000 | 49.0 | 81.67% | 118.7 | 41.65% | motif file (matrix) | svg |
| 22 | C G T A T G A C T C G A A G T C C G T A A T C G A T G C A C G T A C T G A G T C | E2A(bHLH)/proBcell-E2A-ChIP-Seq(GSE21978)/Homer | 1e-8 | -1.860e+01 | 0.0000 | 56.0 | 93.33% | 163.6 | 57.41% | motif file (matrix) | svg |
| 23 | C T A G A G T C T A C G T A C G T G A C C G T A A C T G T A G C G C A T C A T G A T G C A G C T | Ascl1(bHLH)/NeuralTubes-Ascl1-ChIP-Seq(GSE55840)/Homer | 1e-7 | -1.824e+01 | 0.0000 | 55.0 | 91.67% | 157.6 | 55.31% | motif file (matrix) | svg |
| 24 | A G T C C G T A C G T A A C G T G C A T C G T A A C G T A C G T | ATHB20(Homeobox)/colamp-ATHB20-DAP-Seq(GSE60143)/Homer | 1e-7 | -1.805e+01 | 0.0000 | 57.0 | 95.00% | 173.1 | 60.74% | motif file (matrix) | svg |
| 25 | T C A G A C T G C A G T A G T C A G T C G T C A C G T A C G T A A C T G C A G T A G T C A G T C C T G A T G C A A G C T | dHNF4(NR)/Fly-HNF4-ChIP-Seq(GSE73675)/Homer | 1e-7 | -1.796e+01 | 0.0000 | 18.0 | 30.00% | 11.9 | 4.18% | motif file (matrix) | svg |
| 26 | C T A G A C T G C T A G T C A G T C A G T A C G C T A G A C T G | Maz(Zf)/HepG2-Maz-ChIP-Seq(GSE31477)/Homer | 1e-7 | -1.778e+01 | 0.0000 | 41.0 | 68.33% | 83.4 | 29.27% | motif file (matrix) | svg |
| 27 | A T G C G C A T T A G C C G A T T A G C G C A T T A G C G C A T A T G C G A C T | GAGA-repeat/Arabidopsis-Promoters/Homer | 1e-7 | -1.742e+01 | 0.0000 | 49.0 | 81.67% | 123.4 | 43.29% | motif file (matrix) | svg |
| 28 | C G T A C G T A C G T A A G T C A G C T C T G A A C T G A C T G A G C T A G T C C G T A C T A G C T A G C T A G T G C A | RORa(NR)/Liver-Rora-ChIP-Seq(GSE101115)/Homer | 1e-7 | -1.741e+01 | 0.0000 | 30.0 | 50.00% | 43.9 | 15.39% | motif file (matrix) | svg |
| 29 | T C G A A G T C C G T A A T C G A T G C C G A T A C T G A G T C A G C T A C T G | Tcf12(bHLH)/GM12878-Tcf12-ChIP-Seq(GSE32465)/Homer | 1e-7 | -1.726e+01 | 0.0000 | 45.0 | 75.00% | 103.4 | 36.26% | motif file (matrix) | svg |
| 30 | T G A C C G A T C T G A C T A G C T A G A C G T A T G C T G C A T C G A C T G A C T A G C A T G A C G T A G T C C G T A | PPARa(NR),DR1/Liver-Ppara-ChIP-Seq(GSE47954)/Homer | 1e-7 | -1.722e+01 | 0.0000 | 54.0 | 90.00% | 154.3 | 54.13% | motif file (matrix) | svg |
| 31 | C T A G C T A G T C G A C G T A A T G C C G T A A T C G T C G A T A C G G C A T A C T G C A G T T A G C G A T C G A C T | MRE(NR)/Neuro2A-NR3C2-ChIPnexus(GSE115417)/Homer | 1e-7 | -1.660e+01 | 0.0000 | 60.0 | 100.00% | 209.2 | 73.39% | motif file (matrix) | svg |
| 32 | T A G C G T A C A G T C G T A C C G A T A G T C A G T C A G T C A G T C A G T C C G T A G A T C | Zfp281(Zf)/ES-Zfp281-ChIP-Seq(GSE81042)/Homer | 1e-7 | -1.627e+01 | 0.0000 | 9.0 | 15.00% | 0.7 | 0.25% | motif file (matrix) | svg |
| 33 | A C G T G A C T T A G C C G T A C T G A C A T G C T A G G A C T G A T C C G T A | Nr5a2(NR)/Pancreas-LRH1-ChIP-Seq(GSE34295)/Homer | 1e-6 | -1.591e+01 | 0.0000 | 47.0 | 78.33% | 118.7 | 41.66% | motif file (matrix) | svg |
| 34 | G A T C C G T A G A C T C T A G G A T C C T G A G A C T C T G A G A C T C T A G G A T C C T G A G A C T C T G A G A C T | OCT:OCT(POU,Homeobox)/NPC-OCT6-ChIP-Seq(GSE43916)/Homer | 1e-6 | -1.548e+01 | 0.0000 | 20.0 | 33.33% | 19.7 | 6.90% | motif file (matrix) | svg |
| 35 | C T A G T C A G C T G A C T G A C T A G C T G A C A T G C A T G C T G A C T A G C T A G C G T A C T A G C G T A G T C A | TF3A(C2H2)/col-TF3A-DAP-Seq(GSE60143)/Homer | 1e-6 | -1.496e+01 | 0.0000 | 48.0 | 80.00% | 127.8 | 44.85% | motif file (matrix) | svg |
| 36 | C G A T T A C G T G C A G T A C G A T C G A C T A G C T A C G T A T C G G T A C G A T C G T A C G A T C G T C A | PPARE(NR),DR1/3T3L1-Pparg-ChIP-Seq(GSE13511)/Homer | 1e-6 | -1.422e+01 | 0.0000 | 48.0 | 80.00% | 130.7 | 45.85% | motif file (matrix) | svg |
| 37 | A C G T T G A C A G T C A G C T A G T C A G C T A C T G G A C T A G C T G A C T | REF6(Zf)/Arabidopsis-REF6-ChIP-Seq(GSE106942)/Homer | 1e-5 | -1.368e+01 | 0.0000 | 37.0 | 61.67% | 80.3 | 28.16% | motif file (matrix) | svg |
| 38 | C T A G T C G A T G A C A G T C C G T A A C T G G T A C A C G T A C T G A C T G | BHLHA15(bHLH)/NIH3T3-BHLHB8.HA-ChIP-Seq(GSE119782)/Homer | 1e-5 | -1.358e+01 | 0.0000 | 55.0 | 91.67% | 176.5 | 61.94% | motif file (matrix) | svg |
| 39 | A C G T T C G A T C G A A G T C G T C A T A C G A T G C A C G T A C T G A G C T | Myf5(bHLH)/GM-Myf5-ChIP-Seq(GSE24852)/Homer | 1e-5 | -1.355e+01 | 0.0000 | 39.0 | 65.00% | 89.1 | 31.25% | motif file (matrix) | svg |
| 40 | A G T C G T A C A G C T C T A G A G T C C G A T A C T G C G T A A C T G G T C A | Zic(Zf)/Cerebellum-ZIC1.2-ChIP-Seq(GSE60731)/Homer | 1e-5 | -1.342e+01 | 0.0000 | 47.0 | 78.33% | 128.8 | 45.19% | motif file (matrix) | svg |
| 41 | A C G T G A C T A T G C G C T A C T G A C T A G A C T G G A C T A G T C C G T A | Nr5a2(NR)/mES-Nr5a2-ChIP-Seq(GSE19019)/Homer | 1e-5 | -1.340e+01 | 0.0000 | 40.0 | 66.67% | 94.4 | 33.14% | motif file (matrix) | svg |
| 42 | C A T G A C G T A G T C G A T C G A T C G A T C G C T A C T A G C T A G C T A G T C A G T C G A | EBF1(EBF)/Near-E2A-ChIP-Seq(GSE21512)/Homer | 1e-5 | -1.328e+01 | 0.0000 | 41.0 | 68.33% | 99.7 | 35.00% | motif file (matrix) | svg |
| 43 | T G A C A G T C C G T A A C T G G T A C A C G T A C T G A C G T G A C T G A T C | Twist2(bHLH)/Myoblast-Twist2.Ty1-ChIP-Seq(GSE127998)/Homer | 1e-5 | -1.325e+01 | 0.0000 | 59.0 | 98.33% | 211.3 | 74.14% | motif file (matrix) | svg |
| 44 | C G A T C T A G C T G A A T G C C T G A T C G A C G T A C T G A T C G A T A G C A G T C C G T A A C T G T C G A A T G C | Hand2(bHLH)/Mesoderm-Hand2-ChIP-Seq(GSE61475)/Homer | 1e-5 | -1.317e+01 | 0.0000 | 32.0 | 53.33% | 62.5 | 21.93% | motif file (matrix) | svg |
| 45 | C G A T C G T A G T C A C G T A A G T C A G T C A G T C A C G T | TRP2(MYBrelated)/colamp-TRP2-DAP-Seq(GSE60143)/Homer | 1e-5 | -1.317e+01 | 0.0000 | 32.0 | 53.33% | 62.6 | 21.96% | motif file (matrix) | svg |
| 46 | G T C A T G C A G C T A A G T C C G T A A C T G T G A C G C A T T C A G C A G T | Ap4(bHLH)/AML-Tfap4-ChIP-Seq(GSE45738)/Homer | 1e-5 | -1.304e+01 | 0.0000 | 48.0 | 80.00% | 135.5 | 47.53% | motif file (matrix) | svg |
| 47 | T A C G T G C A A G T C C G T A A C G T T G A C A C G T A C T G A C T G G C A T | TCF4(bHLH)/SHSY5Y-TCF4-ChIP-Seq(GSE96915)/Homer | 1e-5 | -1.296e+01 | 0.0001 | 57.0 | 95.00% | 194.9 | 68.39% | motif file (matrix) | svg |
| 48 | G A C T C T A G C T A G C T A G A C T G T C G A C T G A C T A G C T A G C T A G G T A C G T C A | ZNF467(Zf)/HEK293-ZNF467.GFP-ChIP-Seq(GSE58341)/Homer | 1e-5 | -1.287e+01 | 0.0001 | 37.0 | 61.67% | 83.4 | 29.27% | motif file (matrix) | svg |
| 49 | T G A C A T G C C G T A A T C G A T G C C A G T C A T G A C T G A G T C G T A C | HEB(bHLH)/mES-Heb-ChIP-Seq(GSE53233)/Homer | 1e-5 | -1.255e+01 | 0.0001 | 59.0 | 98.33% | 214.2 | 75.15% | motif file (matrix) | svg |
| 50 | C T G A A T G C G C T A C G A T A T G C C G T A C G T A C G T A C T A G T A C G | Tcf3(HMG)/mES-Tcf3-ChIP-Seq(GSE11724)/Homer | 1e-5 | -1.254e+01 | 0.0001 | 33.0 | 55.00% | 68.4 | 24.00% | motif file (matrix) | svg |
| 51 | T C A G T G A C G T A C C G T A A C G T T G A C A C G T T C A G A G C T G A C T | NeuroD1(bHLH)/Islet-NeuroD1-ChIP-Seq(GSE30298)/Homer | 1e-5 | -1.253e+01 | 0.0001 | 41.0 | 68.33% | 102.1 | 35.84% | motif file (matrix) | svg |
| 52 | G A C T A G C T A G C T C T A G A C G T G A T C A C G T A C G T G A C T C G A T G C A T A G T C | IDD4(C2H2)/col-IDD4-DAP-Seq(GSE60143)/Homer | 1e-5 | -1.244e+01 | 0.0001 | 50.0 | 83.33% | 149.9 | 52.58% | motif file (matrix) | svg |
| 53 | T A C G C T G A C A T G G A T C G T A C G C A T T C A G T A C G A G C T G T C A G A T C G C A T T A C G C G T A C T A G G A T C G A T C C G A T A C T G T C A G | ZNF322(Zf)/HEK293-ZNF322.GFP-ChIP-Seq(GSE58341)/Homer | 1e-5 | -1.237e+01 | 0.0001 | 16.0 | 26.67% | 15.2 | 5.34% | motif file (matrix) | svg |
| 54 | C T G A C T A G T C G A C G T A A T G C C G T A A T C G C G A T T A G C G C A T A T C G G C A T A G C T G A T C G A C T A G C T | ARE(NR)/LNCAP-AR-ChIP-Seq(GSE27824)/Homer | 1e-5 | -1.163e+01 | 0.0002 | 24.0 | 40.00% | 39.2 | 13.76% | motif file (matrix) | svg |
| 55 | C A T G A G C T T A C G G T C A G T A C T A G C A G C T G A C T A T C G T C G A | Esrrb(NR)/mES-Esrrb-ChIP-Seq(GSE11431)/Homer | 1e-5 | -1.161e+01 | 0.0002 | 47.0 | 78.33% | 136.8 | 47.99% | motif file (matrix) | svg |
| 56 | A C T G A G C T A G T C G T C A A G C T T C A G A T G C G A T C G C A T A T C G T C G A T A G C C G A T C A T G T A G C | Pax8(Paired,Homeobox)/Thyroid-Pax8-ChIP-Seq(GSE26938)/Homer | 1e-5 | -1.154e+01 | 0.0002 | 22.0 | 36.67% | 33.1 | 11.60% | motif file (matrix) | svg |
| 57 | T C A G C T G A C T A G C A T G A C G T A T G C C T G A C T G A C T G A C T A G C A T G A C G T A T G C C T G A | TR4(NR),DR1/Hela-TR4-ChIP-Seq(GSE24685)/Homer | 1e-4 | -1.147e+01 | 0.0002 | 14.0 | 23.33% | 12.0 | 4.22% | motif file (matrix) | svg |
| 58 | T C A G A C G T T C G A T A G C A G T C C G T A A C T G G T A C A C G T A C T G A T C G A G T C | Atoh1(bHLH)/Cerebellum-Atoh1-ChIP-Seq(GSE22111)/Homer | 1e-4 | -1.128e+01 | 0.0002 | 48.0 | 80.00% | 143.9 | 50.48% | motif file (matrix) | svg |
| 59 | T G C A C G T A G T C A A G C T A G T C G C T A T A G C C G A T C T A G G A T C | Gfi1b(Zf)/HPC7-Gfi1b-ChIP-Seq(GSE22178)/Homer | 1e-4 | -1.126e+01 | 0.0002 | 44.0 | 73.33% | 122.7 | 43.06% | motif file (matrix) | svg |
| 60 | A T G C T C A G T C G A G C A T A C T G C G T A A G T C T C A G G A C T T G A C C G T A A G C T | Atf2(bZIP)/3T3L1-Atf2-ChIP-Seq(GSE56872)/Homer | 1e-4 | -1.094e+01 | 0.0003 | 32.0 | 53.33% | 70.1 | 24.60% | motif file (matrix) | svg |
| 61 | C G T A C T G A A C T G A G C T A G T C G A T C G A T C G C A T C T G A C T A G C T A G T A C G T C G A T G C A G C A T | EBF2(EBF)/BrownAdipose-EBF2-ChIP-Seq(GSE97114)/Homer | 1e-4 | -1.086e+01 | 0.0003 | 42.0 | 70.00% | 114.2 | 40.08% | motif file (matrix) | svg |
| 62 | A T G C T C G A A G T C A G C T A C G T G T A C A G T C G C T A C T A G C A T G G T C A C T G A T C A G A G T C | Stat3+il21(Stat)/CD4-Stat3-ChIP-Seq(GSE19198)/Homer | 1e-4 | -1.083e+01 | 0.0003 | 44.0 | 73.33% | 124.1 | 43.56% | motif file (matrix) | svg |
| 63 | C A T G G T C A A G T C C G T A C T A G G A T C C G A T A C T G A C G T G T A C C G T A C G T A | bZIP69(bZIP)/col-bZIP69-DAP-Seq(GSE60143)/Homer | 1e-4 | -1.075e+01 | 0.0004 | 13.0 | 21.67% | 11.8 | 4.15% | motif file (matrix) | svg |
| 64 | C T G A C G A T A C G T G C A T C G T A C G T A A C G T A C T G | At1g76110(ARID)/colamp-At1g76110-DAP-Seq(GSE60143)/Homer | 1e-4 | -1.074e+01 | 0.0004 | 55.0 | 91.67% | 190.0 | 66.66% | motif file (matrix) | svg |
| 65 | T C A G G A C T G T C A C G T A A C G T A T C G C G T A A C G T A C G T C T G A | ATHB15(HB)/col-ATHB15-DAP-Seq(GSE60143)/Homer | 1e-4 | -1.072e+01 | 0.0004 | 51.0 | 85.00% | 163.4 | 57.34% | motif file (matrix) | svg |
| 66 | C A T G A C T G A G C T A T G C C G T A A T G C G T A C G A C T T A C G C T G A A C T G A C T G G C A T A T G C C T G A | THRb(NR)/HepG2-THRb.Flag-ChIP-Seq(Encode)/Homer | 1e-4 | -1.064e+01 | 0.0004 | 37.0 | 61.67% | 92.1 | 32.31% | motif file (matrix) | svg |
| 67 | T C G A T C A G T C G A A C T G C A T G A C G T A G T C C T G A | COUP-TFII(NR)/Artia-Nr2f2-ChIP-Seq(GSE46497)/Homer | 1e-4 | -1.062e+01 | 0.0004 | 60.0 | 100.00% | 234.2 | 82.16% | motif file (matrix) | svg |
| 68 | C G A T T G C A T G C A G A T C C G T A A C T G T G A C G A C T C A T G A C T G | Tcf21(bHLH)/ArterySmoothMuscle-Tcf21-ChIP-Seq(GSE61369)/Homer | 1e-4 | -1.048e+01 | 0.0004 | 41.0 | 68.33% | 111.1 | 38.99% | motif file (matrix) | svg |
| 69 | T C G A C A T G C A T G A C G T A T G C T C G A C T G A A G C T T A C G T G C A G T A C G A T C A G C T A G T C | FXR(NR),IR1/Liver-FXR-ChIP-Seq(Chong\_et\_al.)/Homer | 1e-4 | -1.043e+01 | 0.0004 | 32.0 | 53.33% | 72.5 | 25.42% | motif file (matrix) | svg |
| 70 | C A G T T C A G T C G A A G T C C G T A A C T G T G A C C G A T A C T G A C T G A C G T A T C G | Atoh7(bHLH)/Retina-Atoh7-CutnRun(GSE156756)/Homer | 1e-4 | -1.029e+01 | 0.0005 | 38.0 | 63.33% | 98.4 | 34.53% | motif file (matrix) | svg |
| 71 | C A T G T A C G T A G C G A T C G A T C A T G C G T A C G A C T T C A G A T G C C G A T A T C G C A G T A C T G G T A C | Zic3(Zf)/mES-Zic3-ChIP-Seq(GSE37889)/Homer | 1e-4 | -1.027e+01 | 0.0005 | 29.0 | 48.33% | 61.9 | 21.72% | motif file (matrix) | svg |
| 72 | C A T G G C T A C T A G T A C G C G T A T C A G C G T A A C T G C G T A C A T G C T G A C G T A | BPC1(BBRBPC)/colamp-BPC1-DAP-Seq(GSE60143)/Homer | 1e-4 | -1.018e+01 | 0.0006 | 32.0 | 53.33% | 73.5 | 25.77% | motif file (matrix) | svg |
| 73 | A T G C T G C A A G T C C G T A A G T C A G T C A C G T A C T G A T G C G T C A | E2A(bHLH),near\_PU.1/Bcell-PU.1-ChIP-Seq(GSE21512)/Homer | 1e-4 | -1.013e+01 | 0.0006 | 55.0 | 91.67% | 192.9 | 67.69% | motif file (matrix) | svg |
| 74 | A T G C T C G A T A C G A C G T A T G C A G T C A C G T A G T C A G T C G A T C | Znf263(Zf)/K562-Znf263-ChIP-Seq(GSE31477)/Homer | 1e-4 | -1.012e+01 | 0.0006 | 51.0 | 85.00% | 166.8 | 58.54% | motif file (matrix) | svg |
| 75 | T C G A T G A C G T A C C G T A C A G T T G A C A C G T A C T G A G C T A G C T | NeuroG2(bHLH)/Fibroblast-NeuroG2-ChIP-Seq(GSE75910)/Homer | 1e-4 | -9.778e+00 | 0.0008 | 56.0 | 93.33% | 201.2 | 70.60% | motif file (matrix) | svg |
| 76 | C T A G A C T G T G C A A G T C C G T A A C T G A C T G A C G T C T A G C G A T T A C G A G T C | ZEB2(Zf)/SNU398-ZEB2-ChIP-Seq(GSE103048)/Homer | 1e-4 | -9.775e+00 | 0.0008 | 49.0 | 81.67% | 156.6 | 54.95% | motif file (matrix) | svg |
| 77 | C T A G C A T G A C G T A G T C G C T A A G C T A G T C A G C T T C A G C T G A A C T G C A T G G C A T A T G C C G T A | THRa(NR)/C17.2-THRa-ChIP-Seq(GSE38347)/Homer | 1e-4 | -9.725e+00 | 0.0008 | 33.0 | 55.00% | 79.7 | 27.97% | motif file (matrix) | svg |
| 78 | T G C A C T G A A T G C G T C A A C G T A T G C A C G T A C T G A C T G T G C A | ZBTB18(Zf)/HEK293-ZBTB18.GFP-ChIP-Seq(GSE58341)/Homer | 1e-4 | -9.717e+00 | 0.0008 | 30.0 | 50.00% | 68.0 | 23.85% | motif file (matrix) | svg |
| 79 | C A T G A C T G C T A G T C G A T C G A T C G A T C G A T C A G T C A G T C A G T G A C T G A C C G T A A C T G T G C A C G A T A C T G | RBPJ:Ebox(?,bHLH)/Panc1-Rbpj1-ChIP-Seq(GSE47459)/Homer | 1e-4 | -9.704e+00 | 0.0008 | 18.0 | 30.00% | 26.7 | 9.35% | motif file (matrix) | svg |
| 80 | T C A G T G A C G T A C T G C A G T A C C T A G G T A C A T G C A G T C G T C A A G T C G A C T | Klf9(Zf)/GBM-Klf9-ChIP-Seq(GSE62211)/Homer | 1e-4 | -9.593e+00 | 0.0009 | 15.0 | 25.00% | 18.8 | 6.60% | motif file (matrix) | svg |
| 81 | A G C T T C G A G T A C T C G A A T G C A T G C G C A T A T C G A G T C A G C T | Snail1(Zf)/LS174T-SNAIL1.HA-ChIP-Seq(GSE127183)/Homer | 1e-4 | -9.481e+00 | 0.0010 | 46.0 | 76.67% | 141.9 | 49.79% | motif file (matrix) | svg |
| 82 | G A C T C G A T C T G A G T C A G A C T C G A T T C G A C G T A G C T A G C T A T G A C G T A C C G T A A C T G T G C A C G A T A C T G A C G T | Pitx1:Ebox(Homeobox,bHLH)/Hindlimb-Pitx1-ChIP-Seq(GSE41591)/Homer | 1e-4 | -9.305e+00 | 0.0012 | 23.0 | 38.33% | 43.8 | 15.38% | motif file (matrix) | svg |
| 83 | C A T G G A C T T A C G G T C A G T A C G A T C G A C T A G C T A T C G T C G A T A C G T A G C | ERRg(NR)/Kidney-ESRRG-ChIP-Seq(GSE104905)/Homer | 1e-4 | -9.266e+00 | 0.0012 | 48.0 | 80.00% | 153.4 | 53.81% | motif file (matrix) | svg |
| 84 | T A G C G C A T A G T C G A T C A T G C G A C T C T A G A C T G A C T G C T G A A C T G C T A G A G T C T G A C C G A T | GLIS3(Zf)/Thyroid-Glis3.GFP-ChIP-Seq(GSE103297)/Homer | 1e-4 | -9.238e+00 | 0.0012 | 45.0 | 75.00% | 137.9 | 48.40% | motif file (matrix) | svg |
| 85 | A G C T C T G A C T A G C T A G A C T G T A G C T G C A T C G A C T G A C T A G C A T G A C G T A T G C T C G A | RXR(NR),DR1/3T3L1-RXR-ChIP-Seq(GSE13511)/Homer | 1e-4 | -9.238e+00 | 0.0012 | 45.0 | 75.00% | 137.3 | 48.17% | motif file (matrix) | svg |
| 86 | G T A C G A T C C A G T A G T C A G T C A G T C T G C A G A T C C T G A A T G C G T C A A C G T | WT1(Zf)/Kidney-WT1-ChIP-Seq(GSE90016)/Homer | 1e-4 | -9.234e+00 | 0.0012 | 30.0 | 50.00% | 69.7 | 24.46% | motif file (matrix) | svg |
| 87 | G A C T T C A G C T A G A G T C A G T C G T A C A G T C C T G A A G T C A G T C A G T C G A C T A G T C A C T G A T G C | KLF3(Zf)/MEF-Klf3-ChIP-Seq(GSE44748)/Homer | 1e-3 | -9.108e+00 | 0.0013 | 21.0 | 35.00% | 37.3 | 13.09% | motif file (matrix) | svg |
| 88 | C T G A T C A G C T G A C T A G C A T G A C G T A T G C C G T A A T G C G C A T T C A G C T G A A C T G A C G T C A G T A G T C C G T A C A G T C T A G C A T G | VDR(NR),DR3/GM10855-VDR+vitD-ChIP-Seq(GSE22484)/Homer | 1e-3 | -8.993e+00 | 0.0015 | 19.0 | 31.67% | 31.2 | 10.96% | motif file (matrix) | svg |
| 89 | C T G A A C G T A C G T A C G T A G T C G A C T C G A T C T G A A C T G C G T A C G T A T C G A | STAT5(Stat)/mCD4+-Stat5-ChIP-Seq(GSE12346)/Homer | 1e-3 | -8.781e+00 | 0.0018 | 29.0 | 48.33% | 67.8 | 23.79% | motif file (matrix) | svg |
| 90 | G C T A C G A T C T A G G T A C G C T A C G A T C G T A G C T A C G A T C A G T G A T C G C T A | Pit1+1bp(Homeobox)/GCrat-Pit1-ChIP-Seq(GSE58009)/Homer | 1e-3 | -8.746e+00 | 0.0019 | 39.0 | 65.00% | 110.7 | 38.85% | motif file (matrix) | svg |
| 91 | T G C A G C A T A G C T G C A T A G T C A G T C A G T C C T G A A C T G T C G A T C G A C A G T A T C G A G T C G A T C | ZNF143\|STAF(Zf)/CUTLL-ZNF143-ChIP-Seq(GSE29600)/Homer | 1e-3 | -8.632e+00 | 0.0021 | 18.0 | 30.00% | 29.4 | 10.30% | motif file (matrix) | svg |
| 92 | C A T G A C T G C A T G A C T G T A C G A T G C A G T C G T A C G T A C T G A C G A T C G A C T | TCP1(TCP)/col-TCP1-DAP-Seq(GSE60143)/Homer | 1e-3 | -8.608e+00 | 0.0021 | 24.0 | 40.00% | 49.2 | 17.25% | motif file (matrix) | svg |
| 93 | C T G A A T G C G C T A G C A T A T G C C G T A T C G A C T G A C T A G T C A G T A C G G T C A | Tcf4(HMG)/Hct116-Tcf4-ChIP-Seq(SRA012054)/Homer | 1e-3 | -8.513e+00 | 0.0023 | 37.0 | 61.67% | 102.3 | 35.90% | motif file (matrix) | svg |
| 94 | C T G A C T A G C T A G C A G T A G T C C T G A T A C G C T A G T A C G G A C T T A C G G T C A G A T C G T A C A G C T | ERb(NR),IR3/Ovary-ERb-ChIP-Seq(GSE203391)/Homer | 1e-3 | -8.507e+00 | 0.0023 | 25.0 | 41.67% | 53.9 | 18.92% | motif file (matrix) | svg |
| 95 | T A C G G A C T A C T G A C T G C T A G A T G C A G T C A G T C A G T C C T G A | ZNF692(Zf)/HEK293-ZNF692.GFP-ChIP-Seq(GSE58341)/Homer | 1e-3 | -8.444e+00 | 0.0024 | 14.0 | 23.33% | 18.6 | 6.54% | motif file (matrix) | svg |
| 96 | C T G A T C G A C G T A A T G C C G T A C G T A C G A T C T A G T C A G G A T C | Sox15(HMG)/CPA-Sox15-ChIP-Seq(GSE62909)/Homer | 1e-3 | -8.404e+00 | 0.0024 | 54.0 | 90.00% | 195.0 | 68.41% | motif file (matrix) | svg |
| 97 | G A C T A G T C C G A T A C T G C T G A T G A C G T A C C G T A A T C G G C A T C T G A C T A G | Bcl11a(Zf)/HSPC-BCL11A-ChIP-Seq(GSE104676)/Homer | 1e-3 | -8.360e+00 | 0.0025 | 39.0 | 65.00% | 112.2 | 39.36% | motif file (matrix) | svg |
| 98 | A G T C C T G A A T C G A G C T A G C T G A C T A G T C G C T A A C G T C G A T G C A T C G A T A T C G C G T A T A G C G C A T A T G C C G T A | bZIP:IRF(bZIP,IRF)/Th17-BatF-ChIP-Seq(GSE39756)/Homer | 1e-3 | -8.323e+00 | 0.0026 | 30.0 | 50.00% | 73.7 | 25.85% | motif file (matrix) | svg |
| 99 | C A T G G T A C G A C T G C T A C G T A C G T A C G T A G C T A G A C T C T G A T C A G G T A C | Mef2c(MADS)/GM12878-Mef2c-ChIP-Seq(GSE32465)/Homer | 1e-3 | -8.310e+00 | 0.0026 | 40.0 | 66.67% | 117.9 | 41.36% | motif file (matrix) | svg |
| 100 | A G T C A G T C C G A T A C G T A C G T A C T G A C G T A G C T A G T C A G T C | Sox4(HMG)/proB-Sox4-ChIP-Seq(GSE50066)/Homer | 1e-3 | -8.305e+00 | 0.0026 | 45.0 | 75.00% | 142.3 | 49.91% | motif file (matrix) | svg |
| 101 | T C G A G C A T A C T G C T G A A G T C T C A G G A C T G T A C C G T A A G C T A G T C G A T C | c-Jun-CRE(bZIP)/K562-cJun-ChIP-Seq(GSE31477)/Homer | 1e-3 | -8.257e+00 | 0.0027 | 25.0 | 41.67% | 54.9 | 19.26% | motif file (matrix) | svg |
| 102 | G C A T C T A G G T A C A G T C C G A T A C T G C T A G C T A G G T A C G C T A | ZNF416(Zf)/HEK293-ZNF416.GFP-ChIP-Seq(GSE58341)/Homer | 1e-3 | -8.177e+00 | 0.0029 | 46.0 | 76.67% | 148.9 | 52.23% | motif file (matrix) | svg |
| 103 | A C G T C T A G G T C A G C T A C G A T G C T A G C T A G C A T C A G T G A C T T G C A C A G T | POU4F3(POU,Homeobox)/MEF-Pou4f3-ChIP-Seq(GSE150279)/Homer | 1e-3 | -8.143e+00 | 0.0030 | 27.0 | 45.00% | 62.6 | 21.95% | motif file (matrix) | svg |
| 104 | A T G C C T G A G A C T A C G T A C G T G T A C G A T C C G A T C T A G C A T G C G T A C G T A C T G A G A C T | STAT1(Stat)/HelaS3-STAT1-ChIP-Seq(GSE12782)/Homer | 1e-3 | -8.115e+00 | 0.0030 | 28.0 | 46.67% | 67.0 | 23.50% | motif file (matrix) | svg |
| 105 | A C T G G A T C G A C T A C T G A C G T C A T G A C T G A C G T A G C T C G A T | RUNX-AML(Runt)/CD4+-PolII-ChIP-Seq(Barski\_et\_al.)/Homer | 1e-3 | -8.092e+00 | 0.0031 | 41.0 | 68.33% | 123.7 | 43.42% | motif file (matrix) | svg |
| 106 | A T G C A G T C C T G A A G T C C G A T A C G T A G T C A G T C A C G T A T C G G A C T A C G T | Etv2(ETS)/ES-ER71-ChIP-Seq(GSE59402)/Homer | 1e-3 | -8.075e+00 | 0.0031 | 42.0 | 70.00% | 128.8 | 45.18% | motif file (matrix) | svg |
| 107 | G C T A G C A T T A C G A G C T C A T G A C T G C A T G C A T G G A T C A G T C A G T C C T G A A G T C G T A C G C T A | At1g69690(TCP)/colamp-At1g69690-DAP-Seq(GSE60143)/Homer | 1e-3 | -7.947e+00 | 0.0035 | 21.0 | 35.00% | 41.5 | 14.57% | motif file (matrix) | svg |
| 108 | C G A T C G A T G C A T G A C T A C G T C G T A C G T A A C T G T A G C C G T A C G T A C G T A | AT5G60130(ABI3VP1)/col-AT5G60130-DAP-Seq(GSE60143)/Homer | 1e-3 | -7.944e+00 | 0.0035 | 60.0 | 100.00% | 246.8 | 86.60% | motif file (matrix) | svg |
| 109 | G C T A T C G A C G T A C T A G A G C T G T C A G T C A C G T A A G T C C G T A | FOXA1(Forkhead)/MCF7-FOXA1-ChIP-Seq(GSE26831)/Homer | 1e-3 | -7.941e+00 | 0.0035 | 53.0 | 88.33% | 190.1 | 66.71% | motif file (matrix) | svg |
| 110 | T C G A A C G T A C T G C T G A A G T C T C A G A G C T G T A C C G T A A G C T G A T C T C G A | JunD(bZIP)/K562-JunD-ChIP-Seq/Homer | 1e-3 | -7.915e+00 | 0.0035 | 12.0 | 20.00% | 14.1 | 4.96% | motif file (matrix) | svg |
| 111 | C T G A C G A T C A G T C A T G A C T G A T C G G T C A A G T C A G T C G T C A A G T C G C T A | PTF1(TCP)/colamp-PTF1-DAP-Seq(GSE60143)/Homer | 1e-3 | -7.915e+00 | 0.0035 | 12.0 | 20.00% | 14.9 | 5.22% | motif file (matrix) | svg |
| 112 | C G T A T G A C T A G C T G C A A C T G A C T G C G T A C G T A T C A G G A C T | ELF3(ETS)/PDAC-ELF3-ChIP-Seq(GSE64557)/Homer | 1e-3 | -7.910e+00 | 0.0035 | 41.0 | 68.33% | 124.7 | 43.76% | motif file (matrix) | svg |
| 113 | T A C G C T A G T A C G A G T C C G T A A G T C A G T C A C G T A C T G A G T C G A T C T A G C | Slug(Zf)/Mesoderm-Snai2-ChIP-Seq(GSE61475)/Homer | 1e-3 | -7.887e+00 | 0.0035 | 29.0 | 48.33% | 72.0 | 25.25% | motif file (matrix) | svg |
| 114 | A C T G A C G T A C T G A C T G C A G T A G T C A G T C A G C T C G T A T A C G G A C T A G C T A G C T T A G C C G T A | TCP16(TCP)/colamp-TCP16-DAP-Seq(GSE60143)/Homer | 1e-3 | -7.672e+00 | 0.0043 | 34.0 | 56.67% | 93.8 | 32.91% | motif file (matrix) | svg |
| 115 | T C G A T A G C G T C A A C T G A C T G C G T A C G T A C T A G A G C T T C A G | ERG(ETS)/VCaP-ERG-ChIP-Seq(GSE14097)/Homer | 1e-3 | -7.586e+00 | 0.0046 | 53.0 | 88.33% | 192.0 | 67.37% | motif file (matrix) | svg |
| 116 | T C A G T C A G T A G C A G T C C T G A A G T C C T A G A C G T A C T G A T C G | c-Myc(bHLH)/mES-cMyc-ChIP-Seq(GSE11431)/Homer | 1e-3 | -7.482e+00 | 0.0051 | 34.0 | 56.67% | 95.0 | 33.33% | motif file (matrix) | svg |
| 117 | G C A T C G T A G C A T C G T A T C G A C G T A C T G A A C T G C G T A C G T A C G T A A C G T A C T G G T C A G C A T | AT2G31460(REMB3)/col-AT2G31460-DAP-Seq(GSE60143)/Homer | 1e-3 | -7.374e+00 | 0.0056 | 35.0 | 58.33% | 99.8 | 35.02% | motif file (matrix) | svg |
| 118 | A C T G A T C G A G T C A C G T C G T A A G T C A G T C A C G T A C T G C G T A | Zelda(Zf)/Embryo-zld-ChIP-Seq(GSE65441)/Homer | 1e-3 | -7.371e+00 | 0.0056 | 42.0 | 70.00% | 132.2 | 46.39% | motif file (matrix) | svg |
| 119 | C G T A C T A G G A C T G T C A G T C A C G T A A G T C C G T A T C G A T C G A T C G A C G T A C T G A C T A G G C T A C G T A T A G C C G T A C G A T C G T A | FOXA1:AR(Forkhead,NR)/LNCAP-AR-ChIP-Seq(GSE27824)/Homer | 1e-3 | -7.349e+00 | 0.0057 | 9.0 | 15.00% | 8.5 | 2.98% | motif file (matrix) | svg |
| 120 | A T C G A G C T C T G A C T A G A C T G A C G T G T A C G C T A A T G C A C G T C T A G C A T G T A C G C G A T A T G C C G T A | Reverb(NR),DR2/RAW-Reverba.biotin-ChIP-Seq(GSE45914)/Homer | 1e-3 | -7.278e+00 | 0.0060 | 8.0 | 13.33% | 6.2 | 2.17% | motif file (matrix) | svg |
| 121 | C T A G C T A G A T G C G T A C T C A G A T G C A G T C G C A T G A T C G A T C | ZNF91(Zf)/HEK-ZNF91.HA-ChIP-Seq(GSE162571)/Homer | 1e-3 | -7.269e+00 | 0.0060 | 39.0 | 65.00% | 118.3 | 41.51% | motif file (matrix) | svg |
| 122 | T C A G T C A G A C G T G T A C G C T A T C A G C T G A A C T G A C T G A G C T A G T C C G T A | EAR2(NR)/K562-NR2F6-ChIP-Seq(Encode)/Homer | 1e-3 | -6.973e+00 | 0.0081 | 54.0 | 90.00% | 202.8 | 71.14% | motif file (matrix) | svg |
| 123 | C T A G T C A G C T G A T C A G T G C A A C T G T C G A T C A G | Trl(Zf)/S2-GAGAfactor-ChIP-Seq(GSE40646)/Homer | 1e-3 | -6.963e+00 | 0.0081 | 59.0 | 98.33% | 240.5 | 84.38% | motif file (matrix) | svg |
| 124 | G C A T A C G T A C G T A T C G C G T A C G T A C G T A C G T A | At2g41835(C2H2)/col-At2g41835-DAP-Seq(GSE60143)/Homer | 1e-2 | -6.894e+00 | 0.0086 | 40.0 | 66.67% | 125.5 | 44.03% | motif file (matrix) | svg |
| 125 | A G T C G A C T C A G T A C T G C T A G T G A C G C T A A T G C G C A T A T C G C G A T A C T G G A T C G T A C G T C A C T G A | NF1(CTF)/LNCAP-NF1-ChIP-Seq(Unpublished)/Homer | 1e-2 | -6.829e+00 | 0.0091 | 22.0 | 36.67% | 49.7 | 17.43% | motif file (matrix) | svg |
| 126 | A C G T A T G C A C G T A C G T A G C T A G T C A G C T A G C T A G C T A G C T A G C T | hTCT(CPE) | 1e-2 | -6.816e+00 | 0.0091 | 57.0 | 95.00% | 224.6 | 78.82% | motif file (matrix) | svg |
| 127 | C T A G T C A G C A G T T C A G A C T G A C T G G A T C C T A G A C T G C T A G T C A G A T G C | KLF14(Zf)/HEK293-KLF14.GFP-ChIP-Seq(GSE58341)/Homer | 1e-2 | -6.765e+00 | 0.0095 | 45.0 | 75.00% | 151.0 | 52.98% | motif file (matrix) | svg |
| 128 | T G A C G C T A T C G A T G C A A G T C A G T C C G T A A G T C C G T A C T G A G C T A G T A C | RUNX2(Runt)/PCa-RUNX2-ChIP-Seq(GSE33889)/Homer | 1e-2 | -6.719e+00 | 0.0099 | 47.0 | 78.33% | 162.4 | 56.99% | motif file (matrix) | svg |
| 129 | T A C G T A C G C T A G T C A G A G T C C G T A A T C G A T G C A C G T A C T G A G T C G A C T | Ascl2(bHLH)/ESC-Ascl2-ChIP-Seq(GSE97712)/Homer | 1e-2 | -6.710e+00 | 0.0099 | 41.0 | 68.33% | 131.5 | 46.15% | motif file (matrix) | svg |
| 130 | T C G A A G T C C G T A A T C G T A G C A C G T A C T G A G C T A C G T A G T C | Ptf1a(bHLH)/Panc1-Ptf1a-ChIP-Seq(GSE47459)/Homer | 1e-2 | -6.651e+00 | 0.0104 | 60.0 | 100.00% | 252.6 | 88.64% | motif file (matrix) | svg |
| 131 | G T A C C G T A C G T A A C G T G C T A C G T A A C G T C A G T | ATHB13(Homeobox)/col-ATHB13-DAP-Seq(GSE60143)/Homer | 1e-2 | -6.617e+00 | 0.0107 | 58.0 | 96.67% | 233.7 | 81.98% | motif file (matrix) | svg |
| 132 | T G C A C G T A A C T G T C A G C A G T C A T G T C A G G A T C T A C G A G T C T G C A A C T G A C T G T G A C G T C A | ZNF165(Zf)/WHIM12-ZNF165-ChIP-Seq(GSE65937)/Homer | 1e-2 | -6.610e+00 | 0.0107 | 7.0 | 11.67% | 5.1 | 1.79% | motif file (matrix) | svg |
| 133 | A T G C A G T C A G C T A G C T A C G T A T C G C G T A C G A T T A G C G A C T | LEF1(HMG)/H1-LEF1-ChIP-Seq(GSE64758)/Homer | 1e-2 | -6.495e+00 | 0.0119 | 46.0 | 76.67% | 159.0 | 55.78% | motif file (matrix) | svg |
| 134 | T A G C A G T C T G A C A G T C C T A G A T C G A G T C C A T G T G A C A G T C G T A C A G T C A G T C G C A T C T A G A T C G G C A T A C T G A T C G G A T C | BORIS(Zf)/K562-CTCFL-ChIP-Seq(GSE32465)/Homer | 1e-2 | -6.387e+00 | 0.0132 | 11.0 | 18.33% | 15.6 | 5.46% | motif file (matrix) | svg |
| 135 | T A C G T A G C G C T A C G A T C T A G A C G T C A G T C A G T G C T A A G T C G T C A G C A T | FOXK2(Forkhead)/U2OS-FOXK2-ChIP-Seq(E-MTAB-2204)/Homer | 1e-2 | -6.264e+00 | 0.0147 | 49.0 | 81.67% | 176.2 | 61.84% | motif file (matrix) | svg |
| 136 | T C A G A G C T G T C A C G T A A C G T A T G C C G T A A C G T A C G T C T G A | PHV(HB)/col-PHV-DAP-Seq(GSE60143)/Homer | 1e-2 | -6.264e+00 | 0.0147 | 49.0 | 81.67% | 176.2 | 61.81% | motif file (matrix) | svg |
| 137 | A G T C A C G T A C T G A G C T A C G T A C G T G T C A A G T C | Foxo1(Forkhead)/RAW-Foxo1-ChIP-Seq(Fan\_et\_al.)/Homer | 1e-2 | -6.226e+00 | 0.0151 | 60.0 | 100.00% | 254.3 | 89.22% | motif file (matrix) | svg |
| 138 | C T G A T A C G G C A T C T A G A T G C G A T C C G A T A C T G C T A G G A T C C T G A A T G C | MYRF(MYRF)/CFPAC1-MYRF-ChIP-Seq(GSE145627)/Homer | 1e-2 | -6.123e+00 | 0.0167 | 23.0 | 38.33% | 56.1 | 19.69% | motif file (matrix) | svg |
| 139 | A G T C C G A T A C T G A T C G T G A C G C T A C A T G A T C G T G A C C G A T A C T G T A G C G T A C G T C A | Tlx?(NR)/NPC-H3K4me1-ChIP-Seq(GSE16256)/Homer | 1e-2 | -6.120e+00 | 0.0167 | 16.0 | 26.67% | 31.4 | 11.02% | motif file (matrix) | svg |
| 140 | T A C G A T C G T A G C G A T C A C T G A C G T A G T C A C G T C T A G A T C G | Smad4(MAD)/ESC-SMAD4-ChIP-Seq(GSE29422)/Homer | 1e-2 | -6.098e+00 | 0.0168 | 57.0 | 95.00% | 228.0 | 80.00% | motif file (matrix) | svg |
| 141 | C A G T C G T A C G T A G C A T G A C T G C A T G T A C A G C T A C T G G A C T A C G T C A T G | RAV1(RAV)/colamp-RAV1-DAP-Seq(GSE60143)/Homer | 1e-2 | -6.092e+00 | 0.0168 | 47.0 | 78.33% | 166.8 | 58.52% | motif file (matrix) | svg |
| 142 | A C G T C T A G C G T A A G T C G T A C A C G T A C G T A C G T G T C A G T A C T G A C G A C T | Nur77(NR)/K562-NR4A1-ChIP-Seq(GSE31363)/Homer | 1e-2 | -6.082e+00 | 0.0169 | 27.0 | 45.00% | 72.5 | 25.42% | motif file (matrix) | svg |
| 143 | G A C T C T A G G A T C C A G T A C T G C T G A A T G C G C A T A T G C C T G A | MafA(bZIP)/Islet-MafA-ChIP-Seq(GSE30298)/Homer | 1e-2 | -6.074e+00 | 0.0169 | 42.0 | 70.00% | 140.7 | 49.36% | motif file (matrix) | svg |
| 144 | G C T A C G T A A C G T C A T G C G T A A C G T A C G T C T A G | ATHB5(HB)/colamp-ATHB5-DAP-Seq(GSE60143)/Homer | 1e-2 | -6.059e+00 | 0.0170 | 58.0 | 96.67% | 236.4 | 82.96% | motif file (matrix) | svg |
| 145 | T A C G C T G A C T A G T C G A C G T A A G T C C T G A A T C G G C A T T A G C G A C T A C T G A C G T A G C T A G T C G A C T | GRE(NR),IR3/A549-GR-ChIP-Seq(GSE32465)/Homer | 1e-2 | -5.966e+00 | 0.0185 | 13.0 | 21.67% | 22.2 | 7.78% | motif file (matrix) | svg |
| 146 | A G T C G T C A G C T A C G A T T G C A C G T A C G A T C A G T A T C G C T A G | ATHB6(Homeobox)/Arabidopsis-HB6-ChIP-Seq(GSE80564)/Homer | 1e-2 | -5.922e+00 | 0.0193 | 57.0 | 95.00% | 229.8 | 80.62% | motif file (matrix) | svg |
| 147 | C G T A G C T A C G A T C T A G A C G T G T C A C G T A C G T A A G T C C G T A T G C A T A C G | FoxL2(Forkhead)/Ovary-FoxL2-ChIP-Seq(GSE60858)/Homer | 1e-2 | -5.890e+00 | 0.0197 | 51.0 | 85.00% | 190.8 | 66.94% | motif file (matrix) | svg |
| 148 | G A T C G C T A A G T C A C G T A C T G C G T A A G T C C G T A G C T A C G A T C A G T G C A T G C T A C G T A G C A T | GRF9(GRF)/colamp-GRF9-DAP-Seq(GSE60143)/Homer | 1e-2 | -5.761e+00 | 0.0223 | 50.0 | 83.33% | 185.4 | 65.05% | motif file (matrix) | svg |
| 149 | C T G A C G A T C T A G T C A G G A T C C T G A T C A G G A T C C T G A A C T G A G T C G C T A A C G T A G T C G C A T | PRDM9(Zf)/Testis-DMC1-ChIP-Seq(GSE35498)/Homer | 1e-2 | -5.724e+00 | 0.0230 | 27.0 | 45.00% | 75.0 | 26.30% | motif file (matrix) | svg |
| 150 | T G C A A C T G T A C G A T G C A G T C G A C T T C G A A T C G | ZNF711(Zf)/SHSY5Y-ZNF711-ChIP-Seq(GSE20673)/Homer | 1e-2 | -5.651e+00 | 0.0246 | 54.0 | 90.00% | 210.9 | 74.00% | motif file (matrix) | svg |
| 151 | C T A G A G C T C A T G C A T G C T A G A T G C G A T C G T A C G T A C C T G A A G T C G C T A G C T A G C A T A T C G | TCP7(TCP)/col-TCP7-DAP-Seq(GSE60143)/Homer | 1e-2 | -5.614e+00 | 0.0253 | 29.0 | 48.33% | 83.1 | 29.15% | motif file (matrix) | svg |
| 152 | A C T G A C T G C A T G G A T C A G T C A G T C C T G A A G T C | TCP20(TCP)/col-TCP20-DAP-Seq(GSE60143)/Homer | 1e-2 | -5.599e+00 | 0.0255 | 21.0 | 35.00% | 51.6 | 18.11% | motif file (matrix) | svg |
| 153 | G A C T T C G A C G A T T A C G C T A G T A C G A C T G A T G C G T A C G T A C | Zac1(Zf)/Neuro2A-Plagl1-ChIP-Seq(GSE75942)/Homer | 1e-2 | -5.515e+00 | 0.0276 | 58.0 | 96.67% | 239.5 | 84.04% | motif file (matrix) | svg |
| 154 | T G C A C T G A C A T G C T A G C A G T A G T C C G T A A T G C A T G C T A C G G C A T T C A G G T C A G A T C G T A C | ERE(NR),IR3/MCF7-ERa-ChIP-Seq(Unpublished)/Homer | 1e-2 | -5.401e+00 | 0.0307 | 21.0 | 35.00% | 52.1 | 18.26% | motif file (matrix) | svg |
| 155 | G C T A T A G C A G C T A T C G G T C A C G T A G C T A A T G C G A T C C T G A | IRF4(IRF)/GM12878-IRF4-ChIP-Seq(GSE32465)/Homer | 1e-2 | -5.347e+00 | 0.0322 | 40.0 | 66.67% | 135.7 | 47.62% | motif file (matrix) | svg |
| 156 | C T A G C T A G G C A T C G A T C T G A G T C A G C T A A T G C C G T A C A G T G A C T C G T A T G C A | Hnf1(Homeobox)/Liver-Foxa2-Chip-Seq(GSE25694)/Homer | 1e-2 | -5.343e+00 | 0.0322 | 24.0 | 40.00% | 64.4 | 22.61% | motif file (matrix) | svg |
| 157 | T C G A A C T G C A T G A G C T A G T C C G T A C T G A C T A G A C T G C G A T A T G C C T G A | RAR:RXR(NR),DR0/ES-RAR-ChIP-Seq(GSE56893)/Homer | 1e-2 | -5.341e+00 | 0.0322 | 12.0 | 20.00% | 22.0 | 7.71% | motif file (matrix) | svg |
| 158 | G T A C T C G A T A G C C G T A C G T A C T G A T G C A T G A C A C T G C G T A A G T C C T G A C T G A T C G A C G T A | NUC(C2H2)/col-NUC-DAP-Seq(GSE60143)/Homer | 1e-2 | -5.341e+00 | 0.0322 | 12.0 | 20.00% | 21.9 | 7.70% | motif file (matrix) | svg |
| 159 | G C A T G C A T C T G A A C G T C T G A A C G T C G T A C G T A C G T A A G T C G T C A G T C A | Foxf1(Forkhead)/Lung-Foxf1-ChIP-Seq(GSE77951)/Homer | 1e-2 | -5.338e+00 | 0.0322 | 54.0 | 90.00% | 212.3 | 74.48% | motif file (matrix) | svg |
| 160 | C T G A C G A T C A T G A T C G G C A T C A T G G C T A A G T C | ASHR1(ND)/col-ASHR1-DAP-Seq(GSE60143)/Homer | 1e-2 | -5.338e+00 | 0.0322 | 54.0 | 90.00% | 212.2 | 74.44% | motif file (matrix) | svg |
| 161 | T C A G A G C T A C T G A C T G C A G T A G T C A G T C A G T C G T A C C T G A | TCP17(TCP)/col-TCP17-DAP-Seq(GSE60143)/Homer | 1e-2 | -5.296e+00 | 0.0327 | 5.0 | 8.33% | 3.2 | 1.12% | motif file (matrix) | svg |
| 162 | C G A T A C G T G T A C A G C T C T G A A C T G C G T A C T G A A C T G T G A C C G A T A C G T A G T C A G C T C T G A | AT1G10720(BSD)/col-AT1G10720-DAP-Seq(GSE60143)/Homer | 1e-2 | -5.290e+00 | 0.0327 | 3.0 | 5.00% | 0.0 | 0.00% | motif file (matrix) | svg |
| 163 | G A T C G T A C C G A T A C T G A C T G C G T A C G T A A C G T A C T G G A T C | TEAD(TEA)/Fibroblast-PU.1-ChIP-Seq(Unpublished)/Homer | 1e-2 | -5.223e+00 | 0.0347 | 39.0 | 65.00% | 131.7 | 46.21% | motif file (matrix) | svg |
| 164 | T G A C C T G A C T A G T C G A C T G A A T G C C G T A A C T G G C A T G T A C G C A T A T C G G C A T A G C T G A T C | PR(NR)/T47D-PR-ChIP-Seq(GSE31130)/Homer | 1e-2 | -5.177e+00 | 0.0361 | 60.0 | 100.00% | 259.6 | 91.10% | motif file (matrix) | svg |
| 165 | T G A C C T A G T C A G G T C A C G T A T C A G C G A T T C A G T C G A T G C A C T G A T A G C | PU.1-IRF(ETS:IRF)/Bcell-PU.1-ChIP-Seq(GSE21512)/Homer | 1e-2 | -5.158e+00 | 0.0366 | 46.0 | 76.67% | 167.1 | 58.64% | motif file (matrix) | svg |
| 166 | T C G A A C T G A C T G C G T A C G T A T C G A A G T C C T G A A T C G G T A C G C A T C A T G | ETS:E-box(ETS,bHLH)/HPC7-Scl-ChIP-Seq(GSE22178)/Homer | 1e-2 | -5.071e+00 | 0.0397 | 9.0 | 15.00% | 13.7 | 4.81% | motif file (matrix) | svg |
| 167 | A G T C T G C A T C G A C T G A A C T G C A T G A C G T A T G C G T C A T A C G | Erra(NR)/HepG2-Erra-ChIP-Seq(GSE31477)/Homer | 1e-2 | -5.068e+00 | 0.0397 | 57.0 | 95.00% | 235.0 | 82.45% | motif file (matrix) | svg |
| 168 | T G C A G T A C C G T A A T C G A C T G A C G T C T A G C G A T T C G A A G T C | ZEB1(Zf)/PDAC-ZEB1-ChIP-Seq(GSE64557)/Homer | 1e-2 | -5.068e+00 | 0.0397 | 57.0 | 95.00% | 235.0 | 82.44% | motif file (matrix) | svg |
| 169 | T G A C C G T A C T G A A C T G A C T G G A C T G A T C T G C A G T A C T A C G | SF1(NR)/H295R-Nr5a1-ChIP-Seq(GSE44220)/Homer | 1e-2 | -5.007e+00 | 0.0415 | 25.0 | 41.67% | 70.7 | 24.80% | motif file (matrix) | svg |
| 170 | G C T A C G A T C A T G A T G C A G T C A G T C G A C T T A C G T C G A C T A G A C T G T A G C | AP-2alpha(AP2)/Hela-AP2alpha-ChIP-Seq(GSE31477)/Homer | 1e-2 | -5.001e+00 | 0.0415 | 32.0 | 53.33% | 100.5 | 35.27% | motif file (matrix) | svg |
| 171 | T C G A G A C T T C A G T G C A G T A C G T A C A G C T G T A C C A T G T C G A C A T G C A T G A C G T A G T C C T G A | FXR(NR),ER2/Liver-FXR-ChIP-Seq(GSE133700)/Homer | 1e-2 | -4.936e+00 | 0.0441 | 31.0 | 51.67% | 97.0 | 34.03% | motif file (matrix) | svg |
| 172 | T C A G A G C T A C G T A C G T G T A C G A T C C G T A C T A G C A T G G T C A C G T A T C G A | STAT4(Stat)/CD4-Stat4-ChIP-Seq(GSE22104)/Homer | 1e-2 | -4.883e+00 | 0.0462 | 46.0 | 76.67% | 169.6 | 59.51% | motif file (matrix) | svg |
| 173 | A C G T C T A G A G C T A C G T A C G T C T G A A G T C G A C T A G C T C G T A | FOXM1(Forkhead)/MCF7-FOXM1-ChIP-Seq(GSE72977)/Homer | 1e-2 | -4.882e+00 | 0.0462 | 54.0 | 90.00% | 215.7 | 75.68% | motif file (matrix) | svg |
| 174 | C T G A C T A G C A T G C T A G A C T G A C T G G A T C A C T G A C T G C T A G T C A G A G T C | Klf15(Zf)/Liver-Klf15-ChIP-Seq(GSE166083)/Homer | 1e-2 | -4.831e+00 | 0.0481 | 38.0 | 63.33% | 129.1 | 45.31% | motif file (matrix) | svg |
| 175 | C T G A A C T G A C G T A C T G A C G T A G T C C G T A A C T G | CELF2(RRM)/JSL1-CELF2-CLIP-Seq(GSE71264)/Homer | 1e-2 | -4.829e+00 | 0.0481 | 24.0 | 40.00% | 67.8 | 23.78% | motif file (matrix) | svg |
| 176 | G A T C G A T C G C T A G T C A G A C T A T G C T C G A C G A T C G A T C T A G | HAT2(Homeobox)/colamp-HAT2-DAP-Seq(GSE60143)/Homer | 1e-2 | -4.812e+00 | 0.0485 | 58.0 | 96.67% | 243.0 | 85.26% | motif file (matrix) | svg |
| 177 | A T C G A G T C A G T C G A C T A T G C C T G A C T A G A C T G T A C G G T A C C T G A C G A T | AP-2gamma(AP2)/MCF7-TFAP2C-ChIP-Seq(GSE21234)/Homer | 1e-2 | -4.764e+00 | 0.0506 | 36.0 | 60.00% | 120.6 | 42.32% | motif file (matrix) | svg |
| 178 | G T A C C T A G T C A G A G C T T A G C C G T A A T G C T A C G A G T C G T A C G T C A A G T C | Srebp2(bHLH)/HepG2-Srebp2-ChIP-Seq(GSE31477)/Homer | 1e-2 | -4.733e+00 | 0.0519 | 11.0 | 18.33% | 20.6 | 7.21% | motif file (matrix) | svg |
| 179 | A C T G A C T G C T A G G A T C A G T C A G T C C T G A A G T C | At1g72010(TCP)/colamp-At1g72010-DAP-Seq(GSE60143)/Homer | 1e-2 | -4.709e+00 | 0.0528 | 26.0 | 43.33% | 76.3 | 26.75% | motif file (matrix) | svg |
| 180 | A T G C C T G A A T C G T A C G A G T C C G A T T C A G C G A T C T A G A G C T G T C A G T C A C G T A A G T C C G T A T A C G C T G A | Fox:Ebox(Forkhead,bHLH)/Panc1-Foxa2-ChIP-Seq(GSE47459)/Homer | 1e-2 | -4.664e+00 | 0.0549 | 47.0 | 78.33% | 176.2 | 61.82% | motif file (matrix) | svg |
| 181 | C G A T A G T C G T C A T G C A A G C T C G A T C T G A A C G T C G A T T C A G | HAT5(Homeobox)/colamp-HAT5-DAP-Seq(GSE60143)/Homer | 1e-2 | -4.648e+00 | 0.0555 | 31.0 | 51.67% | 98.6 | 34.59% | motif file (matrix) | svg |
