## Supplementary Data 3 for "Evolutionary insights into glucose production in vertebrate development: new findings from Arctic lamprey (*Lethenteron camtschaticum*)": Suplementary Data3_G_g6pc1_mast.html

ACGTN

The name of the motif.

The alternate name of the motif.

The width of the motif. No gaps are allowed in motifs supplied to MAST
as it only works for motifs of a fixed width.

The sequence that would achieve the best possible match score and its
reverse complement for nucleotide motifs.

MAST computes the pairwise correlations between each pair of motifs.
The correlation between two motifs is the maximum sum of Pearson's
correlation coefficients for aligned columns divided by the width of
the shorter motif. The maximum is found by trying all alignments of the
two motifs.

Motifs with correlations below **0.60** have little effect on
the accuracy of the *E*-values computed by MAST. Motifs with higher
correlations with other motifs should be removed from the query. You can
also request MAST to remove redundant motifs from its analysis
under **Advanced options** from the MAST web page,
or by specifying `--remcorr`
when running MAST on your own computer.

The name of the (FASTA) sequence database file.

The number of sequences in the database.

The number of residues in the sequence database.

The date of the last modification to the sequence database.

The name of the file containing the (MEME-formatted) motifs used in the search.

The date of the last modification to the motif database.

This diagram shows the normal spacing of the motifs specified to MAST.

MAST will calculate larger *p*-values for sites that diverge from the order and spacing in the diagram.

The name of the sequence. This maybe be linked to search a sequence database for the sequence name.

The *E*-value of the sequence.

If strands were scored separately then there will be two
*E*-values for the sequence separated by a slash (/). The score for the
provided sequence will be first and the score for the reverse-complement
will be second.

The block diagram shows the best non-overlapping tiling of motif matches on the sequence.
These motif matches are the ones used by MAST to compute the *E*-value for the sequence.
**Hovering the mouse cursor over a motif match causes the display of the motif name,
position *p*-value of the match and other details in the hovering text.**

- The length of the line shows the length of a sequence relative to all the other sequences.
- A block is shown where the position
  *p*-value
  of a motif is less (more significant) than the significance threshold,
  which is 0.0001 by default.
- If a significant motif match (as specified above) overlaps other
  significant motif matches, then it is only displayed as a block if its
  position *p*-value is less (more significant) then the
  product of the position *p*-values of the significant matches that it
  overlaps.
- The position of a block shows where a motif has matched the sequence.
- **Complementable alphabets (like DNA) only:** Blocks displayed above the line are a match on the given sequence, whereas blocks
  displayed below the line are matches to the reverse-complement of the given sequence.
- **Complementable alphabets (like DNA) only:** When strands are scored separately, then blocks may overlap on opposing strands.
- The width of a block shows the width of the motif relative to the length of the sequence.
- The colour and border of a block identifies the matching motif as in the legend.
  **Note: You can change the color of a motif by clicking on the motif in the legend.**
- The height of a block gives an indication of the significance of the match as
  taller blocks are more significant. The height is calculated to be proportional
  to the negative logarithm of the position *p*-value,
  truncated at the height for a *p*-value of 1e-10.

The description appearing after the identifier in the FASTA file used to specify the sequence.

The combined *p*-value of the sequence.

If strands were scored separately with a complementable alphabet then
there will be two *p*-values for the sequence separated by a slash (/).
The score for the given sequence will be first and the score for the
reverse-complement will be second.

This indicates the offset used for translation of the DNA.

The annotated sequence shows a portion of the sequence with the
matching motif sequences displayed above.

The displayed portion of the sequence can be modified by sliding the
two buttons below the sequence block diagram so that the portion you want
to see is between the two needles attached to the buttons. By default the
two buttons move together but you can drag one individually by holding
shift before you start the drag.

If the strands were scored separately then overlaps in motif sites may
occur so you can choose to display only one strand at a time. This is done
by selecting "Matches on given strand" or "Matches on opposite strand"
from the drop-down list.

The name of the alphabet symbol.

The frequency of the alphabet symbol as defined by the background model.

The score for the match of a position in a sequence to a motif is
computed by by summing the appropriate entry from each column of the
position-dependent scoring matrix that represents the motif. Sequences
shorter than one or more of the motifs are skipped.

The position *p*-value of a match is the probability of a single random
subsequence of the length of the motif scoring at least as
well as the observed match.

The sequence *p*-value of a score is defined as the probability of a
random sequence of the same length containing some match with as good or
better a score.

The combined *p*-value of a sequence measures the strength of the match
of the sequence to all the motifs and is calculated by

1. finding the score
   of the single best match of each motif to the sequence (best matches
   may overlap),
2. calculating the sequence *p*-value
   of each score,
3. forming the product of the *p*-values,
4. taking the *p*-value of the product.

The *E*-value of a sequence is the expected number of sequences in a
random database of the same size that would match the motifs as well as
the sequence does and is equal to the combined *p*-value of
the sequence times the number of sequences in the database.

|  |  |
| --- | --- |
| Motif | 1 |
| *p*-value | 8.23e-7 |
| Start | 23 |
| End | 33 |

Change the portion of annotated sequence by **dragging the buttons**; hold shift to drag them individually.

###### Sequence Comment:

###### Combined *p*-value:

 /

###### Annotated Sequence

Matches on either strand
Matches on given strand
Matches on opposite strand

### MAST

#### Motif Alignment & Search Tool

For further information on how to interpret these results or to get a
copy of the MEME software please access
http://meme-suite.org.

If you use MAST in your research, please cite the following paper:  

Timothy L. Bailey and Michael Gribskov,
"Combining evidence using p-values: application to sequence homology searches",
*Bioinformatics*, **14**(1):48-54, 1998.
[pdf]

Motifs
  |  
Search Results
  |  
Inputs & Settings
  |  
Program information

### Javascript is required to view these results!

### Your browser does not support canvas!

#### Motifs

Next Top

Motifs with a pale red background are very similar to other earlier
specified motifs and may be biasing the results.  
It is recommended that you re-run MAST and request it to remove redundant motifs.

Motifs which are grayed-out were very similar to other earlier
specified motifs and were removed from the scan as you requested.

#### Search Results

Prev Next Top

###### Top Scoring Sequences

#### Inputs & Settings

Prev Next Top

###### Alphabet

###### Sequence Alphabet

###### Sequences

The following sequence databases were supplied to MAST.

| Database | Sequence Count | Residue Count | Last Modified |
| --- | --- | --- | --- |

###### Motifs

The following motif databases were supplied to MAST.

| Database | Last Modified |
| --- | --- |

###### Nominal Order and Spacing

The expected order and spacing of the motifs (as specified by you).

###### Other Settings

|  |
| --- |
| Strand Handling |
| Max Correlation |
| Remove Correlated |
| Max Sequence *E*-value |
| Adjust Hit *p*-value |
| Displayed Hits |
| Displayed Weak Hits |

Prev Top

###### MAST version

(Release date: )

###### Reference

Timothy L. Bailey and Michael Gribskov,
"Combining evidence using p-values: application to sequence homology searches",
*Bioinformatics*, **14**(1):48-54, 1998.

###### Command line summary

Ran in  on  on
