## Supplementary Data 4 for "Evolutionary insights into glucose production in vertebrate development: new findings from Arctic lamprey (*Lethenteron camtschaticum*)": Suplementary Data4_G_fbp1_motif.html

fbp1\_motif - Homer Known Motif Enrichment Results


### Homer Known Motif Enrichment Results (fbp1\_motif)

Homer *de novo* Motif Results  
Gene Ontology Enrichment Results  
Known Motif Enrichment Results (txt file)  
Total Target Sequences = 51, Total Background Sequences = 235

|  |  |  |  |  |  |  |  |  |  |  |  |
| --- | --- | --- | --- | --- | --- | --- | --- | --- | --- | --- | --- |
| Rank | Motif | Name | P-value | log P-pvalue | q-value (Benjamini) | # Target Sequences with Motif | % of Targets Sequences with Motif | # Background Sequences with Motif | % of Background Sequences with Motif | Motif File | SVG |
| 1 | G C T A G C A T A C G T A C G T A G C T G A T C G A C T G A C T A G C T A C G T A C G T A G C T | RLR1?/SacCer-Promoters/Homer | 1e-11 | -2.587e+01 | 0.0000 | 37.0 | 72.55% | 50.8 | 21.62% | motif file (matrix) | svg |
| 2 | G C A T A G C T A C G T A C G T A C T G A C G T G A T C A C G T A C G T A G C T C G A T G C A T A G T C G A C T C A G T | IDD5(C2H2)/colamp-IDD5-DAP-Seq(GSE60143)/Homer | 1e-10 | -2.505e+01 | 0.0000 | 44.0 | 86.27% | 83.6 | 35.55% | motif file (matrix) | svg |
| 3 | G A T C G A T C A G T C G T A C C G A T G T A C G T A C A G T C A G T C A G T C G C T A G A T C | ZNF148(Zf)/MDAMB231-ZNF148-ChIP-Seq(GSE147020)/Homer | 1e-10 | -2.381e+01 | 0.0000 | 30.0 | 58.82% | 31.8 | 13.52% | motif file (matrix) | svg |
| 4 | A C G T A C G T A C G T A C G T A C G T A C G T A C G T A C G T A C G T A C G T | VRN1(ABI3VP1)/col-VRN1-DAP-Seq(GSE60143)/Homer | 1e-9 | -2.230e+01 | 0.0000 | 17.0 | 33.33% | 5.8 | 2.46% | motif file (matrix) | svg |
| 5 | G T C A T G C A T G C A G C T A C G T A G C T A G C T A G C T A | REM19(REM)/colamp-REM19-DAP-Seq(GSE60143)/Homer | 1e-9 | -2.143e+01 | 0.0000 | 40.0 | 78.43% | 73.3 | 31.19% | motif file (matrix) | svg |
| 6 | T A C G T A C G C T A G T C A G A G T C C G T A A T C G A T G C A C G T A C T G A G T C G A C T | Ascl2(bHLH)/ESC-Ascl2-ChIP-Seq(GSE97712)/Homer | 1e-8 | -2.041e+01 | 0.0000 | 46.0 | 90.20% | 108.2 | 46.03% | motif file (matrix) | svg |
| 7 | G T C A T G C A G C T A A G T C C G T A A C T G T G A C G C A T T C A G C A G T | Ap4(bHLH)/AML-Tfap4-ChIP-Seq(GSE45738)/Homer | 1e-8 | -1.980e+01 | 0.0000 | 44.0 | 86.27% | 98.3 | 41.84% | motif file (matrix) | svg |
| 8 | C A T G G C T A C T A G T A C G C G T A T C A G C G T A A C T G C G T A C A T G C T G A C G T A | BPC1(BBRBPC)/colamp-BPC1-DAP-Seq(GSE60143)/Homer | 1e-7 | -1.767e+01 | 0.0000 | 36.0 | 70.59% | 67.0 | 28.50% | motif file (matrix) | svg |
| 9 | C G T A T G A C T C G A A G T C C G T A A T C G A T G C A C G T A C T G A G T C | E2A(bHLH)/proBcell-E2A-ChIP-Seq(GSE21978)/Homer | 1e-7 | -1.682e+01 | 0.0000 | 46.0 | 90.20% | 120.6 | 51.29% | motif file (matrix) | svg |
| 10 | A G C T T C G A G T A C T C G A A T G C A T G C G C A T A T C G A G T C A G C T | Snail1(Zf)/LS174T-SNAIL1.HA-ChIP-Seq(GSE127183)/Homer | 1e-7 | -1.638e+01 | 0.0000 | 40.0 | 78.43% | 88.6 | 37.68% | motif file (matrix) | svg |
| 11 | G T A C G C T A T C A G C T G A C T A G C A T G A G C T G A T C T G C A T C G A C T G A A C T G C A G T A G T C G A T C G C T A | HNF4a(NR),DR1/HepG2-HNF4a-ChIP-Seq(GSE25021)/Homer | 1e-7 | -1.625e+01 | 0.0000 | 30.0 | 58.82% | 47.8 | 20.32% | motif file (matrix) | svg |
| 12 | T A C G C T A G T A C G A G T C C G T A A G T C A G T C A C G T A C T G A G T C G A T C T A G C | Slug(Zf)/Mesoderm-Snai2-ChIP-Seq(GSE61475)/Homer | 1e-6 | -1.590e+01 | 0.0000 | 33.0 | 64.71% | 59.7 | 25.41% | motif file (matrix) | svg |
| 13 | C T A G A G T C T A C G T A C G T G A C C G T A A C T G T A G C G C A T C A T G A T G C A G C T | Ascl1(bHLH)/NeuralTubes-Ascl1-ChIP-Seq(GSE55840)/Homer | 1e-6 | -1.484e+01 | 0.0000 | 47.0 | 92.16% | 134.5 | 57.24% | motif file (matrix) | svg |
| 14 | T C G A A G T C C G T A A T C G A T G C C G A T A C T G A G T C A G C T A C T G | Tcf12(bHLH)/GM12878-Tcf12-ChIP-Seq(GSE32465)/Homer | 1e-6 | -1.472e+01 | 0.0000 | 38.0 | 74.51% | 84.4 | 35.92% | motif file (matrix) | svg |
| 15 | A T G C T G C A A G T C C G T A A G T C A G T C A C G T A C T G A T G C G T C A | E2A(bHLH),near\_PU.1/Bcell-PU.1-ChIP-Seq(GSE21512)/Homer | 1e-6 | -1.466e+01 | 0.0000 | 48.0 | 94.12% | 143.0 | 60.83% | motif file (matrix) | svg |
| 16 | C T A G C T A G T C G A C T A G C G T A A T C G T C G A A C T G C T G A T C G A C T G A T A C G | FRS9(ND)/col-FRS9-DAP-Seq(GSE60143)/Homer | 1e-6 | -1.465e+01 | 0.0000 | 19.0 | 37.25% | 18.6 | 7.93% | motif file (matrix) | svg |
| 17 | C T G A T C A G A G T C C G T A A T C G A T G C C G A T A C T G A G T C G A C T A T C G A G T C | MyoD(bHLH)/Myotube-MyoD-ChIP-Seq(GSE21614)/Homer | 1e-6 | -1.457e+01 | 0.0000 | 34.0 | 66.67% | 67.0 | 28.52% | motif file (matrix) | svg |
| 18 | T C A G A C T G C A G T A G T C A G T C G T C A C G T A C G T A A C T G C A G T A G T C A G T C C T G A T G C A A G C T | dHNF4(NR)/Fly-HNF4-ChIP-Seq(GSE73675)/Homer | 1e-6 | -1.413e+01 | 0.0000 | 10.0 | 19.61% | 2.1 | 0.89% | motif file (matrix) | svg |
| 19 | T G A C G C T A T G A C C G T A T C A G G A T C C G T A C A T G C A T G C T A G C T A G C T A G | Unknown-ESC-element(?)/mES-Nanog-ChIP-Seq(GSE11724)/Homer | 1e-5 | -1.335e+01 | 0.0001 | 26.0 | 50.98% | 41.9 | 17.85% | motif file (matrix) | svg |
| 20 | T G A C A T G C C G T A A T C G A T G C C A G T C A T G A C T G A G T C G T A C | HEB(bHLH)/mES-Heb-ChIP-Seq(GSE53233)/Homer | 1e-5 | -1.313e+01 | 0.0001 | 49.0 | 96.08% | 156.2 | 66.47% | motif file (matrix) | svg |
| 21 | G C T A C G A T C T A G G T A C G C T A C G A T C G T A G C T A C G A T C A G T G A T C G C T A | Pit1+1bp(Homeobox)/GCrat-Pit1-ChIP-Seq(GSE58009)/Homer | 1e-5 | -1.277e+01 | 0.0001 | 38.0 | 74.51% | 91.7 | 39.01% | motif file (matrix) | svg |
| 22 | C A G T C G T A C G T A G C A T G A C T G C A T G T A C A G C T A C T G G A C T A C G T C A T G | RAV1(RAV)/colamp-RAV1-DAP-Seq(GSE60143)/Homer | 1e-5 | -1.253e+01 | 0.0002 | 43.0 | 84.31% | 118.6 | 50.45% | motif file (matrix) | svg |
| 23 | C T G A C T A G C T A G C A G T A G T C C T G A T A C G C T A G T A C G G A C T T A C G G T C A G A T C G T A C A G C T | ERb(NR),IR3/Ovary-ERb-ChIP-Seq(GSE203391)/Homer | 1e-5 | -1.192e+01 | 0.0003 | 22.0 | 43.14% | 32.3 | 13.75% | motif file (matrix) | svg |
| 24 | C T A G A C T G C T A G T C A G T C A G T A C G C T A G A C T G | Maz(Zf)/HepG2-Maz-ChIP-Seq(GSE31477)/Homer | 1e-5 | -1.180e+01 | 0.0003 | 37.0 | 72.55% | 90.4 | 38.46% | motif file (matrix) | svg |
| 25 | T C A G A C G T T C G A T A G C A G T C C G T A A C T G G T A C A C G T A C T G A T C G A G T C | Atoh1(bHLH)/Cerebellum-Atoh1-ChIP-Seq(GSE22111)/Homer | 1e-5 | -1.179e+01 | 0.0003 | 41.0 | 80.39% | 110.7 | 47.10% | motif file (matrix) | svg |
| 26 | T G A C C T A G T C A G G T C A C G T A T C A G C G A T T C A G T C G A T G C A C T G A T A G C | PU.1-IRF(ETS:IRF)/Bcell-PU.1-ChIP-Seq(GSE21512)/Homer | 1e-5 | -1.179e+01 | 0.0003 | 45.0 | 88.24% | 133.5 | 56.78% | motif file (matrix) | svg |
| 27 | T C G A C G T A A G T C C G T A C T A G A G T C C G A T A C T G G A C T A G C T A C T G G A C T | HLH-1(bHLH)/cElegans-Embryo-HLH1-ChIP-Seq(modEncode)/Homer | 1e-5 | -1.155e+01 | 0.0004 | 41.0 | 80.39% | 111.7 | 47.53% | motif file (matrix) | svg |
| 28 | G A C T C A G T A G C T C G A T A G T C G A T C A G T C C G T A A T G C T C A G | Rbpj1(?)/Panc1-Rbpj1-ChIP-Seq(GSE47459)/Homer | 1e-4 | -1.144e+01 | 0.0004 | 48.0 | 94.12% | 155.3 | 66.09% | motif file (matrix) | svg |
| 29 | T C G A T C G A A G T C C G T A C T A G T A G C A C G T A C T G | MyoG(bHLH)/C2C12-MyoG-ChIP-Seq(GSE36024)/Homer | 1e-4 | -1.129e+01 | 0.0005 | 37.0 | 72.55% | 92.9 | 39.54% | motif file (matrix) | svg |
| 30 | C A G T T C A G T C G A A G T C C G T A A C T G T G A C C G A T A C T G A C T G A C G T A T C G | Atoh7(bHLH)/Retina-Atoh7-CutnRun(GSE156756)/Homer | 1e-4 | -1.125e+01 | 0.0005 | 30.0 | 58.82% | 62.5 | 26.57% | motif file (matrix) | svg |
| 31 | G A C T A G C T A G C T C T A G A C G T G A T C A C G T A C G T G A C T C G A T G C A T A G T C | IDD4(C2H2)/col-IDD4-DAP-Seq(GSE60143)/Homer | 1e-4 | -1.106e+01 | 0.0005 | 44.0 | 86.27% | 130.3 | 55.42% | motif file (matrix) | svg |
| 32 | T G C A A G C T C T G A A T C G G A C T C T A G G T A C G A T C G T C A A G T C G T A C G A C T C T A G A T C G G C A T C A T G C A T G G A T C G T A C C T G A | CTCF(Zf)/CD4+-CTCF-ChIP-Seq(Barski\_et\_al.)/Homer | 1e-4 | -1.075e+01 | 0.0007 | 10.0 | 19.61% | 5.1 | 2.17% | motif file (matrix) | svg |
| 33 | A T G C T C G A T A C G A C G T A T G C A G T C A C G T A G T C A G T C G A T C | Znf263(Zf)/K562-Znf263-ChIP-Seq(GSE31477)/Homer | 1e-4 | -1.052e+01 | 0.0009 | 48.0 | 94.12% | 159.8 | 68.00% | motif file (matrix) | svg |
| 34 | C T G A C G A T C T A G T C A G G A T C C T G A T C A G G A T C C T G A A C T G A G T C G C T A A C G T A G T C G C A T | PRDM9(Zf)/Testis-DMC1-ChIP-Seq(GSE35498)/Homer | 1e-4 | -1.049e+01 | 0.0009 | 25.0 | 49.02% | 46.6 | 19.81% | motif file (matrix) | svg |
| 35 | A C G T T G A C A G T C A G C T A G T C A G C T A C T G G A C T A G C T G A C T | REF6(Zf)/Arabidopsis-REF6-ChIP-Seq(GSE106942)/Homer | 1e-4 | -1.038e+01 | 0.0009 | 34.0 | 66.67% | 82.5 | 35.10% | motif file (matrix) | svg |
| 36 | A G C T G A C T A C G T A C T G A C G T A G T C A C T G A C G T G C A T C G A T | AtIDD11(C2H2)/colamp-AtIDD11-DAP-Seq(GSE60143)/Homer | 1e-4 | -1.025e+01 | 0.0010 | 35.0 | 68.63% | 87.3 | 37.13% | motif file (matrix) | svg |
| 37 | T G C A G T A C C G T A A T C G A C T G A C G T C T A G C G A T T C G A A G T C | ZEB1(Zf)/PDAC-ZEB1-ChIP-Seq(GSE64557)/Homer | 1e-4 | -1.017e+01 | 0.0011 | 51.0 | 100.00% | 188.3 | 80.09% | motif file (matrix) | svg |
| 38 | T A G C A G T C T G A C A G T C C T A G A T C G A G T C C A T G T G A C A G T C G T A C A G T C A G T C G C A T C T A G A T C G G C A T A C T G A T C G G A T C | BORIS(Zf)/K562-CTCFL-ChIP-Seq(GSE32465)/Homer | 1e-4 | -9.912e+00 | 0.0014 | 10.0 | 19.61% | 6.4 | 2.74% | motif file (matrix) | svg |
| 39 | G C A T C T A G G T A C A G T C C G A T A C T G C T A G C T A G G T A C G C T A | ZNF416(Zf)/HEK293-ZNF416.GFP-ChIP-Seq(GSE58341)/Homer | 1e-4 | -9.836e+00 | 0.0014 | 39.0 | 76.47% | 108.5 | 46.17% | motif file (matrix) | svg |
| 40 | C T G A T C A G C A G T C T A G A C T G C T A G G A T C A T C G A C T G C T G A T C A G G A T C | Sp5(Zf)/mES-Sp5.Flag-ChIP-Seq(GSE72989)/Homer | 1e-4 | -9.779e+00 | 0.0015 | 35.0 | 68.63% | 89.7 | 38.15% | motif file (matrix) | svg |
| 41 | C T A G T C A G C T G A C T G A C T A G C T G A C A T G C A T G C T G A C T A G C T A G C G T A C T A G C G T A G T C A | TF3A(C2H2)/col-TF3A-DAP-Seq(GSE60143)/Homer | 1e-4 | -9.717e+00 | 0.0015 | 41.0 | 80.39% | 119.2 | 50.71% | motif file (matrix) | svg |
| 42 | C T G A T C A G C T G A C T A G C A T G A C G T A T G C C G T A A T G C G C A T T C A G C T G A A C T G A C G T C A G T A G T C C G T A C A G T C T A G C A T G | VDR(NR),DR3/GM10855-VDR+vitD-ChIP-Seq(GSE22484)/Homer | 1e-4 | -9.635e+00 | 0.0016 | 20.0 | 39.22% | 32.3 | 13.76% | motif file (matrix) | svg |
| 43 | G A C T A C G T A C G T A C T G A C G T A G T C G C A T A G C T G C A T G C A T G A C T A G C T | SGR5(C2H2)/colamp-SGR5-DAP-Seq(GSE60143)/Homer | 1e-4 | -9.501e+00 | 0.0018 | 41.0 | 80.39% | 120.3 | 51.19% | motif file (matrix) | svg |
| 44 | T C A G C T G A C T A G C A T G A C G T A T G C C T G A C T G A C T G A C T A G C A T G A C G T A T G C C T G A | TR4(NR),DR1/Hela-TR4-ChIP-Seq(GSE24685)/Homer | 1e-4 | -9.313e+00 | 0.0022 | 12.0 | 23.53% | 11.8 | 5.02% | motif file (matrix) | svg |
| 45 | A C G T T C G A T C G A A G T C G T C A T A C G A T G C A C G T A C T G A G C T | Myf5(bHLH)/GM-Myf5-ChIP-Seq(GSE24852)/Homer | 1e-3 | -9.167e+00 | 0.0024 | 27.0 | 52.94% | 58.7 | 24.98% | motif file (matrix) | svg |
| 46 | T G C A C T G A A T G C G T C A A C G T A T G C A C G T A C T G A C T G T G C A | ZBTB18(Zf)/HEK293-ZBTB18.GFP-ChIP-Seq(GSE58341)/Homer | 1e-3 | -9.153e+00 | 0.0024 | 22.0 | 43.14% | 40.2 | 17.12% | motif file (matrix) | svg |
| 47 | C G A T T G C A T G C A G A T C C G T A A C T G T G A C G A C T C A T G A C T G | Tcf21(bHLH)/ArterySmoothMuscle-Tcf21-ChIP-Seq(GSE61369)/Homer | 1e-3 | -9.094e+00 | 0.0025 | 33.0 | 64.71% | 83.1 | 35.35% | motif file (matrix) | svg |
| 48 | C T A G A C T G T G C A A G T C C G T A A C T G A C T G A C G T C T A G C G A T T A C G A G T C | ZEB2(Zf)/SNU398-ZEB2-ChIP-Seq(GSE103048)/Homer | 1e-3 | -8.868e+00 | 0.0031 | 41.0 | 80.39% | 123.3 | 52.46% | motif file (matrix) | svg |
| 49 | G A T C G A C T G A C T A C G T A G T C A C G T A G T C A C G T A G T C A C G T A G T C A C G T G T A C C G A T G T C A | BPC6(BBRBPC)/col-BPC6-DAP-Seq(GSE60143)/Homer | 1e-3 | -8.800e+00 | 0.0032 | 6.0 | 11.76% | 1.6 | 0.69% | motif file (matrix) | svg |
| 50 | G C A T G C A T C T G A A C G T C T G A A C G T C G T A C G T A C G T A A G T C G T C A G T C A | Foxf1(Forkhead)/Lung-Foxf1-ChIP-Seq(GSE77951)/Homer | 1e-3 | -8.761e+00 | 0.0033 | 48.0 | 94.12% | 167.9 | 71.41% | motif file (matrix) | svg |
| 51 | G A C T T C A G C T A G A G T C A G T C G T A C A G T C C T G A A G T C A G T C A G T C G A C T A G T C A C T G A T G C | KLF3(Zf)/MEF-Klf3-ChIP-Seq(GSE44748)/Homer | 1e-3 | -8.701e+00 | 0.0034 | 23.0 | 45.10% | 46.0 | 19.55% | motif file (matrix) | svg |
| 52 | G T C A T C G A T C G A C G T A G C T A C G T A T C G A T G A C A C T G C G T A A G T C C G T A C G T A T C G A G C T A | IDD2(C2H2)/colamp-IDD2-DAP-Seq(GSE60143)/Homer | 1e-3 | -8.529e+00 | 0.0040 | 18.0 | 35.29% | 29.1 | 12.38% | motif file (matrix) | svg |
| 53 | A T G C A G C T T C A G T G A C T C A G A T G C T G C A A C G T A T C G G A T C A C T G A G T C | NRF1(NRF)/MCF7-NRF1-ChIP-Seq(Unpublished)/Homer | 1e-3 | -8.505e+00 | 0.0040 | 9.0 | 17.65% | 6.8 | 2.91% | motif file (matrix) | svg |
| 54 | T G C A G C A T C G A T C G T A C A G T A C T G G T A C C G T A C T G A A G C T G T C A A C T G C T A G G T C A C G A T A C T G G T A C T G C A C G T A A G C T | CEBP:CEBP(bZIP)/MEF-Chop-ChIP-Seq(GSE35681)/Homer | 1e-3 | -7.945e+00 | 0.0069 | 19.0 | 37.25% | 35.0 | 14.88% | motif file (matrix) | svg |
| 55 | T A G C T A G C G A C T C T A G A G C T A G T C G T C A T G C A A C G T A T G C G C T A T G C A | Pbx3(Homeobox)/GM12878-PBX3-ChIP-Seq(GSE32465)/Homer | 1e-3 | -7.867e+00 | 0.0073 | 21.0 | 41.18% | 42.0 | 17.87% | motif file (matrix) | svg |
| 56 | C A T G T A C G T A G C G A T C G A T C A T G C G T A C G A C T T C A G A T G C C G A T A T C G C A G T A C T G G T A C | Zic3(Zf)/mES-Zic3-ChIP-Seq(GSE37889)/Homer | 1e-3 | -7.635e+00 | 0.0091 | 23.0 | 45.10% | 49.4 | 21.04% | motif file (matrix) | svg |
| 57 | C G A T C T A G C T G A A T G C C T G A T C G A C G T A C T G A T C G A T A G C A G T C C G T A A C T G T C G A A T G C | Hand2(bHLH)/Mesoderm-Hand2-ChIP-Seq(GSE61475)/Homer | 1e-3 | -7.514e+00 | 0.0100 | 25.0 | 49.02% | 57.2 | 24.35% | motif file (matrix) | svg |
| 58 | A T G C G C A T T A G C C G A T T A G C G C A T T A G C G C A T A T G C G A C T | GAGA-repeat/Arabidopsis-Promoters/Homer | 1e-3 | -7.417e+00 | 0.0109 | 40.0 | 78.43% | 125.4 | 53.37% | motif file (matrix) | svg |
| 59 | G C T A T C G A C G T A C T A G A G C T G T C A G T C A C G T A A G T C C G T A | FOXA1(Forkhead)/MCF7-FOXA1-ChIP-Seq(GSE26831)/Homer | 1e-3 | -7.379e+00 | 0.0111 | 46.0 | 90.20% | 160.2 | 68.14% | motif file (matrix) | svg |
| 60 | G A C T A G T C C G A T A C T G C T G A T G A C G T A C C G T A A T C G G C A T C T G A C T A G | Bcl11a(Zf)/HSPC-BCL11A-ChIP-Seq(GSE104676)/Homer | 1e-3 | -7.352e+00 | 0.0112 | 30.0 | 58.82% | 78.4 | 33.37% | motif file (matrix) | svg |
| 61 | A G C T A G C T A G C T A C T G A C G T A G T C A C T G A C G T G C A T G C A T G C A T A C G T | At5g66730(C2H2)/colamp-At5g66730-DAP-Seq(GSE60143)/Homer | 1e-3 | -7.259e+00 | 0.0121 | 27.0 | 52.94% | 66.2 | 28.17% | motif file (matrix) | svg |
| 62 | C T G A T A C G G C A T C T A G A T G C G A T C C G A T A C T G C T A G G A T C C T G A A T G C | MYRF(MYRF)/CFPAC1-MYRF-ChIP-Seq(GSE145627)/Homer | 1e-3 | -7.145e+00 | 0.0134 | 23.0 | 45.10% | 51.5 | 21.90% | motif file (matrix) | svg |
| 63 | C T A G T C G A T G A C A G T C C G T A A C T G G T A C A C G T A C T G A C T G | BHLHA15(bHLH)/NIH3T3-BHLHB8.HA-ChIP-Seq(GSE119782)/Homer | 1e-3 | -7.142e+00 | 0.0134 | 43.0 | 84.31% | 143.2 | 60.94% | motif file (matrix) | svg |
| 64 | T G A C A G T C C G T A A C T G G T A C A C G T A C T G A C G T G A C T G A T C | Twist2(bHLH)/Myoblast-Twist2.Ty1-ChIP-Seq(GSE127998)/Homer | 1e-3 | -7.122e+00 | 0.0134 | 48.0 | 94.12% | 175.0 | 74.46% | motif file (matrix) | svg |
| 65 | A G T C G T A C A G C T C T A G A G T C C G A T A C T G C G T A A C T G G T C A | Zic(Zf)/Cerebellum-ZIC1.2-ChIP-Seq(GSE60731)/Homer | 1e-3 | -7.046e+00 | 0.0141 | 40.0 | 78.43% | 127.0 | 54.03% | motif file (matrix) | svg |
| 66 | A G T C C T G A A T C G A G C T A G C T G A C T A G T C G C T A A C G T C G A T G C A T C G A T A T C G C G T A T A G C G C A T A T G C C G T A | bZIP:IRF(bZIP,IRF)/Th17-BatF-ChIP-Seq(GSE39756)/Homer | 1e-3 | -7.041e+00 | 0.0141 | 26.0 | 50.98% | 63.7 | 27.10% | motif file (matrix) | svg |
| 67 | A G T C G A C T C A G T A C T G C T A G T G A C G C T A A T G C G C A T A T C G C G A T A C T G G A T C G T A C G T C A C T G A | NF1(CTF)/LNCAP-NF1-ChIP-Seq(Unpublished)/Homer | 1e-2 | -6.902e+00 | 0.0157 | 17.0 | 33.33% | 31.3 | 13.32% | motif file (matrix) | svg |
| 68 | T A C G C T A G A T G C G A T C G T A C A G T C C T A G A G T C A G T C A G T C G T A C A G T C | Sp1(Zf)/Promoter/Homer | 1e-2 | -6.839e+00 | 0.0165 | 13.0 | 25.49% | 19.7 | 8.40% | motif file (matrix) | svg |
| 69 | C A T G A C G T A G T C G A T C G A T C G A T C G C T A C T A G C T A G C T A G T C A G T C G A | EBF1(EBF)/Near-E2A-ChIP-Seq(GSE21512)/Homer | 1e-2 | -6.818e+00 | 0.0166 | 31.0 | 60.78% | 85.9 | 36.54% | motif file (matrix) | svg |
| 70 | T A C G T G C A A G T C C G T A A C G T T G A C A C G T A C T G A C T G G C A T | TCF4(bHLH)/SHSY5Y-TCF4-ChIP-Seq(GSE96915)/Homer | 1e-2 | -6.810e+00 | 0.0166 | 46.0 | 90.20% | 163.1 | 69.37% | motif file (matrix) | svg |
| 71 | C T G A C T A G C A T G C T A G A C T G A C T G G A T C A C T G A C T G C T A G T C A G A G T C | Klf15(Zf)/Liver-Klf15-ChIP-Seq(GSE166083)/Homer | 1e-2 | -6.806e+00 | 0.0166 | 37.0 | 72.55% | 113.8 | 48.43% | motif file (matrix) | svg |
| 72 | T C G A T G A C G C A T A G C T C A G T G A T C G C T A G A T C G A C T A C G T G C A T A G T C | PRDM1(Zf)/Hela-PRDM1-ChIP-Seq(GSE31477)/Homer | 1e-2 | -6.711e+00 | 0.0177 | 32.0 | 62.75% | 90.5 | 38.50% | motif file (matrix) | svg |
| 73 | T C G A A G T C C G T A A T C G T A G C A C G T A C T G A G C T A C G T A G T C | Ptf1a(bHLH)/Panc1-Ptf1a-ChIP-Seq(GSE47459)/Homer | 1e-2 | -6.702e+00 | 0.0177 | 50.0 | 98.04% | 193.2 | 82.18% | motif file (matrix) | svg |
| 74 | T G A C G T A C C G T A A C T G T G A C C G A T A C T G A T C G A G C T T A C G T C G A T A G C G T A C C G T A A T C G T G A C G C A T A C T G A C T G A T G C | Twist(bHLH)/HMLE-TWIST1-ChIP-Seq(Chang\_et\_al)/Homer | 1e-2 | -6.676e+00 | 0.0179 | 9.0 | 17.65% | 9.2 | 3.90% | motif file (matrix) | svg |
| 75 | T A C G C T G A C A T G G A T C G T A C G C A T T C A G T A C G A G C T G T C A G A T C G C A T T A C G C G T A C T A G G A T C G A T C C G A T A C T G T C A G | ZNF322(Zf)/HEK293-ZNF322.GFP-ChIP-Seq(GSE58341)/Homer | 1e-2 | -6.656e+00 | 0.0180 | 11.0 | 21.57% | 14.0 | 5.96% | motif file (matrix) | svg |
| 76 | T C G A T G A C G T A C C G T A C A G T T G A C A C G T A C T G A G C T A G C T | NeuroG2(bHLH)/Fibroblast-NeuroG2-ChIP-Seq(GSE75910)/Homer | 1e-2 | -6.625e+00 | 0.0183 | 46.0 | 90.20% | 164.3 | 69.91% | motif file (matrix) | svg |
| 77 | C A G T A C T G T C A G T G C A G C T A A T G C T C G A A T C G G T C A T G C A | ZNF189(Zf)/HEK293-ZNF189.GFP-ChIP-Seq(GSE58341)/Homer | 1e-2 | -6.501e+00 | 0.0205 | 35.0 | 68.63% | 105.6 | 44.95% | motif file (matrix) | svg |
| 78 | T G C A C T G A C A T G C T A G C A G T A G T C C G T A A T G C A T G C T A C G G C A T T C A G G T C A G A T C G T A C | ERE(NR),IR3/MCF7-ERa-ChIP-Seq(Unpublished)/Homer | 1e-2 | -6.473e+00 | 0.0208 | 14.0 | 27.45% | 23.1 | 9.82% | motif file (matrix) | svg |
| 79 | A G C T T G A C C G A T C G A T C T A G A C G T C A G T C A G T G C T A A G T C | FOXK1(Forkhead)/HEK293-FOXK1-ChIP-Seq(GSE51673)/Homer | 1e-2 | -6.436e+00 | 0.0213 | 49.0 | 96.08% | 187.0 | 79.54% | motif file (matrix) | svg |
| 80 | C G T A G C T A C G A T C T A G A C G T G T C A C G T A C G T A A G T C C G T A T G C A T A C G | FoxL2(Forkhead)/Ovary-FoxL2-ChIP-Seq(GSE60858)/Homer | 1e-2 | -6.393e+00 | 0.0219 | 45.0 | 88.24% | 159.0 | 67.67% | motif file (matrix) | svg |
| 81 | A G C T A G C T A G C T A C T G A C G T A G T C A C T G A C G T G A C T C G A T G C A T A T C G | IDD7(C2H2)/col-IDD7-DAP-Seq(GSE60143)/Homer | 1e-2 | -6.367e+00 | 0.0222 | 30.0 | 58.82% | 83.3 | 35.44% | motif file (matrix) | svg |
| 82 | G A T C G C T A A G T C A C G T A C T G C G T A A G T C C G T A G C T A C G A T C A G T G C A T G C T A C G T A G C A T | GRF9(GRF)/colamp-GRF9-DAP-Seq(GSE60143)/Homer | 1e-2 | -6.215e+00 | 0.0256 | 45.0 | 88.24% | 160.6 | 68.31% | motif file (matrix) | svg |
| 83 | C G T A A C G T A G C T C G A T C T A G G T A C C G T A A G C T C G T A G C T A | Oct4(POU,Homeobox)/mES-Oct4-ChIP-Seq(GSE11431)/Homer | 1e-2 | -6.196e+00 | 0.0258 | 34.0 | 66.67% | 102.9 | 43.76% | motif file (matrix) | svg |
| 84 | C A T G A C T G A G C T A T G C C G T A A T G C G T A C G A C T T A C G C T G A A C T G A C T G G C A T A T G C C T G A | THRb(NR)/HepG2-THRb.Flag-ChIP-Seq(Encode)/Homer | 1e-2 | -6.009e+00 | 0.0307 | 24.0 | 47.06% | 60.2 | 25.62% | motif file (matrix) | svg |
| 85 | C T G A A T G C G C T A G C A T A T G C C G T A T C G A C T G A C T A G T C A G T A C G G T C A | Tcf4(HMG)/Hct116-Tcf4-ChIP-Seq(SRA012054)/Homer | 1e-2 | -6.000e+00 | 0.0307 | 30.0 | 58.82% | 85.0 | 36.17% | motif file (matrix) | svg |
| 86 | C G A T T A C G T G C A G T A C G A T C G A C T A G C T A C G T A T C G G T A C G A T C G T A C G A T C G T C A | PPARE(NR),DR1/3T3L1-Pparg-ChIP-Seq(GSE13511)/Homer | 1e-2 | -5.977e+00 | 0.0309 | 35.0 | 68.63% | 108.3 | 46.08% | motif file (matrix) | svg |
| 87 | A G C T A C G T A C T G A T G C A G T C C G T A C T G A T A C G | NF1-halfsite(CTF)/LNCaP-NF1-ChIP-Seq(Unpublished)/Homer | 1e-2 | -5.971e+00 | 0.0309 | 48.0 | 94.12% | 181.8 | 77.34% | motif file (matrix) | svg |
| 88 | C G T A G C T A C G A T A C G T A G C T A G C T C T G A G T C A C G T A G C T A | Unknown6/Drosophila-Promoters/Homer | 1e-2 | -5.933e+00 | 0.0316 | 37.0 | 72.55% | 118.5 | 50.41% | motif file (matrix) | svg |
| 89 | A G T C C G A T A C T G A T C G T G A C G C T A C A T G A T C G T G A C C G A T A C T G T A G C G T A C G T C A | Tlx?(NR)/NPC-H3K4me1-ChIP-Seq(GSE16256)/Homer | 1e-2 | -5.913e+00 | 0.0319 | 15.0 | 29.41% | 28.0 | 11.93% | motif file (matrix) | svg |
| 90 | G T A C G A T C C A G T A G T C A G T C A G T C T G C A G A T C C T G A A T G C G T C A A C G T | WT1(Zf)/Kidney-WT1-ChIP-Seq(GSE90016)/Homer | 1e-2 | -5.823e+00 | 0.0345 | 26.0 | 50.98% | 69.5 | 29.55% | motif file (matrix) | svg |
| 91 | G A C T C T A G C T A G C T A G A C T G T C G A C T G A C T A G C T A G C T A G G T A C G T C A | ZNF467(Zf)/HEK293-ZNF467.GFP-ChIP-Seq(GSE58341)/Homer | 1e-2 | -5.822e+00 | 0.0345 | 30.0 | 58.82% | 86.8 | 36.94% | motif file (matrix) | svg |
| 92 | A T G C G T A C A G T C A G T C A C G T A C G T C G A T A C G T | AT5G02460(C2C2dof)/col-AT5G02460-DAP-Seq(GSE60143)/Homer | 1e-2 | -5.808e+00 | 0.0345 | 51.0 | 100.00% | 208.0 | 88.49% | motif file (matrix) | svg |
| 93 | G C A T A G C T A G C T A G C T A C T G A C G T A G T C A C T G A C G T G A C T C G A T G C A T | JKD(C2H2)/col-JKD-DAP-Seq(GSE60143)/Homer | 1e-2 | -5.748e+00 | 0.0360 | 20.0 | 39.22% | 46.5 | 19.80% | motif file (matrix) | svg |
| 94 | A G T C G A T C A G T C C G T A A T C G C A G T A G T C G T A C C T G A A C T G T C A G A G C T A G C T A G C T A G C T | PRDM15(Zf)/ESC-Prdm15-ChIP-Seq(GSE73694)/Homer | 1e-2 | -5.719e+00 | 0.0366 | 41.0 | 80.39% | 140.6 | 59.82% | motif file (matrix) | svg |
| 95 | C T G A T C A G G T A C T G C A A G T C C G T A A G T C A C T G A C G T A C T G | MNT(bHLH)/HepG2-MNT-ChIP-Seq(Encode)/Homer | 1e-2 | -5.694e+00 | 0.0372 | 45.0 | 88.24% | 163.3 | 69.46% | motif file (matrix) | svg |
| 96 | C G T A C G T A C T G A A C T G C G T A C G T A A C G T C T A G C G A T C G A T | AT2G38300(G2like)/col-AT2G38300-DAP-Seq(GSE60143)/Homer | 1e-2 | -5.661e+00 | 0.0380 | 49.0 | 96.08% | 190.5 | 81.03% | motif file (matrix) | svg |
| 97 | G T C A G C A T G C T A C A G T C T A G G A T C C G T A C T G A C G T A C G A T | Oct2(POU,Homeobox)/Bcell-Oct2-ChIP-Seq(GSE21512)/Homer | 1e-2 | -5.648e+00 | 0.0381 | 22.0 | 43.14% | 54.5 | 23.20% | motif file (matrix) | svg |
| 98 | A C T G A C T G C A T G G A T C A G T C A G T C C T G A A G T C | TCP20(TCP)/col-TCP20-DAP-Seq(GSE60143)/Homer | 1e-2 | -5.603e+00 | 0.0395 | 19.0 | 37.25% | 43.1 | 18.36% | motif file (matrix) | svg |
| 99 | C T A G T C A G C A G T T C A G A C T G A C T G G A T C C T A G A C T G C T A G T C A G A T G C | KLF14(Zf)/HEK293-KLF14.GFP-ChIP-Seq(GSE58341)/Homer | 1e-2 | -5.556e+00 | 0.0409 | 41.0 | 80.39% | 141.4 | 60.16% | motif file (matrix) | svg |
| 100 | C T G A T C G A C G T A A T G C C G T A C G T A C G A T C T A G T C A G G A T C | Sox15(HMG)/CPA-Sox15-ChIP-Seq(GSE62909)/Homer | 1e-2 | -5.525e+00 | 0.0418 | 45.0 | 88.24% | 164.1 | 69.81% | motif file (matrix) | svg |
| 101 | C G A T C A G T C T A G G C T A A G T C C G T A T C A G A G T C A C G T A C T G A C G T G T A C G C T A G C T A G C T A | bZIP52(bZIP)/colamp-bZIP52-DAP-Seq(GSE60143)/Homer | 1e-2 | -5.460e+00 | 0.0442 | 47.0 | 92.16% | 177.5 | 75.53% | motif file (matrix) | svg |
| 102 | T C A G T G A C G T A C C G T A A C G T T G A C A C G T T C A G A G C T G A C T | NeuroD1(bHLH)/Islet-NeuroD1-ChIP-Seq(GSE30298)/Homer | 1e-2 | -5.411e+00 | 0.0460 | 29.0 | 56.86% | 84.5 | 35.96% | motif file (matrix) | svg |
| 103 | C G A T C A G T C A G T C A T G G T C A G A T C C G T A T C A G A G T C A C G T C T A G A C G T G T A C G T C A G C T A | VIP1(bZIP)/col-VIP1-DAP-Seq(GSE60143)/Homer | 1e-2 | -5.379e+00 | 0.0470 | 15.0 | 29.41% | 30.3 | 12.90% | motif file (matrix) | svg |
| 104 | T C A G C T A G C T A G A C T G A C T G G A T C A C T G A C T G C T A G C T A G A G T C G A T C | KLF1(Zf)/HUDEP2-KLF1-CutnRun(GSE136251)/Homer | 1e-2 | -5.355e+00 | 0.0476 | 28.0 | 54.90% | 80.2 | 34.13% | motif file (matrix) | svg |
| 105 | C A T G G A C T T A C G G T C A G T A C G A T C G A C T A G C T A T C G T C G A T A C G T A G C | ERRg(NR)/Kidney-ESRRG-ChIP-Seq(GSE104905)/Homer | 1e-2 | -5.295e+00 | 0.0501 | 36.0 | 70.59% | 117.3 | 49.89% | motif file (matrix) | svg |
| 106 | A G T C T G C A T C G A C T G A A C T G C A T G A C G T A T G C G T C A T A C G | Erra(NR)/HepG2-Erra-ChIP-Seq(GSE31477)/Homer | 1e-2 | -5.284e+00 | 0.0501 | 49.0 | 96.08% | 192.9 | 82.07% | motif file (matrix) | svg |
| 107 | C T A G C T G A T C A G A T C G A G C T C A T G G A C T A G T C C T G A T G C A | Tbx6(T-box)/ESC-Tbx6-ChIP-Seq(GSE93524)/Homer | 1e-2 | -5.284e+00 | 0.0501 | 49.0 | 96.08% | 192.5 | 81.89% | motif file (matrix) | svg |
| 108 | A G T C G A C T A G C T C G A T A T C G G C T A C G A T A T C G C G A T A C T G T A C G A C G T | Tcf7(HMG)/GM12878-TCF7-ChIP-Seq(Encode)/Homer | 1e-2 | -5.276e+00 | 0.0501 | 26.0 | 50.98% | 72.1 | 30.67% | motif file (matrix) | svg |
| 109 | C T A G A G T C G A T C G T A C A G T C C T A G A G T C G T A C G A T C G T A C G A T C G C A T | KLF17(Zf)/2cell-Klf17-CutnTag(GSE211845)/Homer | 1e-2 | -5.253e+00 | 0.0504 | 25.0 | 49.02% | 68.0 | 28.93% | motif file (matrix) | svg |
| 110 | G C T A T C G A C G T A C T A G A G C T G T C A G T C A C G T A A G T C C G T A | FOXA1(Forkhead)/LNCAP-FOXA1-ChIP-Seq(GSE27824)/Homer | 1e-2 | -5.243e+00 | 0.0504 | 48.0 | 94.12% | 185.1 | 78.76% | motif file (matrix) | svg |
| 111 | C G A T C T A G A C G T A C G T A C G T C G T A A G C T C G A T A G C T C G T A C T A G T A G C | FoxD3(forkhead)/ZebrafishEmbryo-Foxd3.biotin-ChIP-seq(GSE106676)/Homer | 1e-2 | -5.133e+00 | 0.0558 | 38.0 | 74.51% | 128.5 | 54.66% | motif file (matrix) | svg |
| 112 | T A G C T C A G C A T G G C A T A G C T C G A T A T G C C G T A C G T A G T C A | CHR(?)/Hela-CellCycle-Expression/Homer | 1e-2 | -5.129e+00 | 0.0558 | 37.0 | 72.55% | 123.3 | 52.47% | motif file (matrix) | svg |
| 113 | A G T C C G T A A C G T A G T C A C G T A C T G | Tal1 | 1e-2 | -5.086e+00 | 0.0574 | 43.0 | 84.31% | 155.3 | 66.06% | motif file (matrix) | svg |
| 114 | G A C T C G A T T C A G G A T C G A C T A G C T A G C T A G T C G A T C C G T A C T A G C T A G T C G A T C G A C T G A | Bcl6(Zf)/Liver-Bcl6-ChIP-Seq(GSE31578)/Homer | 1e-2 | -5.081e+00 | 0.0574 | 41.0 | 80.39% | 144.6 | 61.53% | motif file (matrix) | svg |
| 115 | A T G C A G T C A G C T A G C T A C G T A T C G C G T A C G A T T A G C G A C T | LEF1(HMG)/H1-LEF1-ChIP-Seq(GSE64758)/Homer | 1e-2 | -4.986e+00 | 0.0623 | 36.0 | 70.59% | 119.3 | 50.74% | motif file (matrix) | svg |
| 116 | C T A G C T A G A T G C G T A C T C A G A T G C A G T C G C A T G A T C G A T C | ZNF91(Zf)/HEK-ZNF91.HA-ChIP-Seq(GSE162571)/Homer | 1e-2 | -4.916e+00 | 0.0663 | 29.0 | 56.86% | 87.8 | 37.34% | motif file (matrix) | svg |
| 117 | A C G T A T G C A C G T A C G T A G C T A G T C A G C T A G C T A G C T A G C T A G C T | hTCT(CPE) | 1e-2 | -4.914e+00 | 0.0663 | 49.0 | 96.08% | 194.0 | 82.55% | motif file (matrix) | svg |
| 118 | A C G T C T A G A G C T A C G T A C G T C T G A A G T C G A C T A G C T C G T A | FOXM1(Forkhead)/MCF7-FOXM1-ChIP-Seq(GSE72977)/Homer | 1e-2 | -4.869e+00 | 0.0683 | 45.0 | 88.24% | 168.4 | 71.63% | motif file (matrix) | svg |
| 119 | C T G A C G T A C G T A G C T A C T G A A C G T G T C A A G T C A G T C C T G A G T C A G C T A | Unknown4/Drosophila-Promoters/Homer | 1e-2 | -4.825e+00 | 0.0708 | 18.0 | 35.29% | 43.4 | 18.47% | motif file (matrix) | svg |
| 120 | C G A T C G A T G C A T G A C T A C G T C G T A C G T A A C T G T A G C C G T A C G T A C G T A | AT5G60130(ABI3VP1)/col-AT5G60130-DAP-Seq(GSE60143)/Homer | 1e-2 | -4.733e+00 | 0.0770 | 49.0 | 96.08% | 195.5 | 83.18% | motif file (matrix) | svg |
| 121 | C T A G C T A G T C G A C G T A A T G C C G T A A T C G T C G A T A C G G C A T A C T G C A G T T A G C G A T C G A C T | MRE(NR)/Neuro2A-NR3C2-ChIPnexus(GSE115417)/Homer | 1e-2 | -4.723e+00 | 0.0771 | 46.0 | 90.20% | 175.7 | 74.76% | motif file (matrix) | svg |
| 122 | G T A C G T A C G T C A G C T A C G T A C G T A C G T A C T A G C T A G C T A G | SEP3(MADS)/Arabidoposis-Flower-Sep3-ChIP-Seq/Homer | 1e-2 | -4.711e+00 | 0.0773 | 45.0 | 88.24% | 169.0 | 71.91% | motif file (matrix) | svg |
| 123 | A T G C A G T C C T G A A G T C C G A T A C G T A G T C A G T C A C G T A T C G G A C T A C G T | Etv2(ETS)/ES-ER71-ChIP-Seq(GSE59402)/Homer | 1e-2 | -4.686e+00 | 0.0786 | 32.0 | 62.75% | 102.8 | 43.72% | motif file (matrix) | svg |
