## Supplementary Data 10 for "Evolutionary insights into glucose production in vertebrate development: new findings from Arctic lamprey (*Lethenteron camtschaticum*)": Suplementary Data10_G_g6pc2_motif.html

g6pc2.2\_motif - Homer Known Motif Enrichment Results


### Homer Known Motif Enrichment Results (g6pc2.2\_motif)

Homer *de novo* Motif Results  
Gene Ontology Enrichment Results  
Known Motif Enrichment Results (txt file)  
Total Target Sequences = 34, Total Background Sequences = 163

|  |  |  |  |  |  |  |  |  |  |  |  |
| --- | --- | --- | --- | --- | --- | --- | --- | --- | --- | --- | --- |
| Rank | Motif | Name | P-value | log P-pvalue | q-value (Benjamini) | # Target Sequences with Motif | % of Targets Sequences with Motif | # Background Sequences with Motif | % of Background Sequences with Motif | Motif File | SVG |
| 1 | T C A G T G A C G T A C C G T A A C G T T G A C A C G T T C A G A G C T G A C T | NeuroD1(bHLH)/Islet-NeuroD1-ChIP-Seq(GSE30298)/Homer | 1e-9 | -2.098e+01 | 0.0000 | 29.0 | 85.29% | 46.2 | 28.42% | motif file (matrix) | svg |
| 2 | C T G A A T G C G C T A G C A T A T G C C G T A T C G A C T G A C T A G T C A G T A C G G T C A | Tcf4(HMG)/Hct116-Tcf4-ChIP-Seq(SRA012054)/Homer | 1e-9 | -2.098e+01 | 0.0000 | 29.0 | 85.29% | 46.7 | 28.72% | motif file (matrix) | svg |
| 3 | G C A T C G T A G C A T C G T A T C G A C G T A C T G A A C T G C G T A C G T A C G T A A C G T A C T G G T C A G C A T | AT2G31460(REMB3)/col-AT2G31460-DAP-Seq(GSE60143)/Homer | 1e-8 | -1.856e+01 | 0.0000 | 31.0 | 91.18% | 63.5 | 39.07% | motif file (matrix) | svg |
| 4 | A C G T T G A C A G T C A G C T A G T C A G C T A C T G G A C T A G C T G A C T | REF6(Zf)/Arabidopsis-REF6-ChIP-Seq(GSE106942)/Homer | 1e-7 | -1.767e+01 | 0.0000 | 28.0 | 82.35% | 49.3 | 30.33% | motif file (matrix) | svg |
| 5 | A C G T A C G T A C G T A C G T A C G T A C G T A C G T A C G T A C G T A C G T | VRN1(ABI3VP1)/col-VRN1-DAP-Seq(GSE60143)/Homer | 1e-7 | -1.712e+01 | 0.0000 | 15.0 | 44.12% | 8.7 | 5.36% | motif file (matrix) | svg |
| 6 | C T G A C T A G T C G A C G T A A T G C C G T A A T C G C G A T T A G C G C A T A T C G G C A T A G C T G A T C G A C T A G C T | ARE(NR)/LNCAP-AR-ChIP-Seq(GSE27824)/Homer | 1e-7 | -1.692e+01 | 0.0000 | 18.0 | 52.94% | 15.6 | 9.63% | motif file (matrix) | svg |
| 7 | G C T A G C A T A C G T A C G T A G C T G A T C G A C T G A C T A G C T A C G T A C G T A G C T | RLR1?/SacCer-Promoters/Homer | 1e-7 | -1.659e+01 | 0.0000 | 27.0 | 79.41% | 47.5 | 29.26% | motif file (matrix) | svg |
| 8 | C T G A A T G C G C T A C G A T A T G C C G T A C G T A C G T A C T A G T A C G | Tcf3(HMG)/mES-Tcf3-ChIP-Seq(GSE11724)/Homer | 1e-6 | -1.591e+01 | 0.0000 | 24.0 | 70.59% | 36.6 | 22.50% | motif file (matrix) | svg |
| 9 | C T A G T C A G C T G A C T G A C T A G C T G A C A T G C A T G C T G A C T A G C T A G C G T A C T A G C G T A G T C A | TF3A(C2H2)/col-TF3A-DAP-Seq(GSE60143)/Homer | 1e-6 | -1.512e+01 | 0.0000 | 31.0 | 91.18% | 73.4 | 45.15% | motif file (matrix) | svg |
| 10 | T C A G A C G T A G T C T C G A A G T C T C A G G C A T C T A G C T A G A G C T | Usf2(bHLH)/C2C12-Usf2-ChIP-Seq(GSE36030)/Homer | 1e-6 | -1.388e+01 | 0.0001 | 23.0 | 67.65% | 37.5 | 23.08% | motif file (matrix) | svg |
| 11 | T C G A T G A C G T A C C G T A C A G T T G A C A C G T A C T G A G C T A G C T | NeuroG2(bHLH)/Fibroblast-NeuroG2-ChIP-Seq(GSE75910)/Homer | 1e-5 | -1.305e+01 | 0.0002 | 34.0 | 100.00% | 105.7 | 65.06% | motif file (matrix) | svg |
| 12 | A G T C G T A C A G C T C T A G A G T C C G A T A C T G C G T A A C T G G T C A | Zic(Zf)/Cerebellum-ZIC1.2-ChIP-Seq(GSE60731)/Homer | 1e-5 | -1.274e+01 | 0.0003 | 26.0 | 76.47% | 53.7 | 33.06% | motif file (matrix) | svg |
| 13 | T A C G C T G A C A T G G A T C G T A C G C A T T C A G T A C G A G C T G T C A G A T C G C A T T A C G C G T A C T A G G A T C G A T C C G A T A C T G T C A G | ZNF322(Zf)/HEK293-ZNF322.GFP-ChIP-Seq(GSE58341)/Homer | 1e-5 | -1.270e+01 | 0.0003 | 8.0 | 23.53% | 1.1 | 0.65% | motif file (matrix) | svg |
| 14 | T G A C C T A G T C A G G T C A C G T A T C A G C G A T T C A G T C G A T G C A C T G A T A G C | PU.1-IRF(ETS:IRF)/Bcell-PU.1-ChIP-Seq(GSE21512)/Homer | 1e-5 | -1.195e+01 | 0.0005 | 34.0 | 100.00% | 109.3 | 67.24% | motif file (matrix) | svg |
| 15 | T C G A A C T G C A T G A G C T A G T C C G T A C T G A C T A G A C T G C G A T A T G C C T G A | RAR:RXR(NR),DR0/ES-RAR-ChIP-Seq(GSE56893)/Homer | 1e-4 | -1.136e+01 | 0.0008 | 12.0 | 35.29% | 9.5 | 5.82% | motif file (matrix) | svg |
| 16 | T A G C A G T C T G A C A G T C C T A G A T C G A G T C C A T G T G A C A G T C G T A C A G T C A G T C G C A T C T A G A T C G G C A T A C T G A T C G G A T C | BORIS(Zf)/K562-CTCFL-ChIP-Seq(GSE32465)/Homer | 1e-4 | -1.122e+01 | 0.0009 | 8.0 | 23.53% | 2.4 | 1.46% | motif file (matrix) | svg |
| 17 | G A C T C T A G C T A G C T A G A C T G T C G A C T G A C T A G C T A G C T A G G T A C G T C A | ZNF467(Zf)/HEK293-ZNF467.GFP-ChIP-Seq(GSE58341)/Homer | 1e-4 | -1.111e+01 | 0.0009 | 18.0 | 52.94% | 26.8 | 16.49% | motif file (matrix) | svg |
| 18 | C T A G T C A G C A G T T C A G A C T G A C T G G A T C C T A G A C T G C T A G T C A G A T G C | KLF14(Zf)/HEK293-KLF14.GFP-ChIP-Seq(GSE58341)/Homer | 1e-4 | -1.087e+01 | 0.0011 | 26.0 | 76.47% | 59.7 | 36.76% | motif file (matrix) | svg |
| 19 | G A T C G A C T G A C T A C G T A G T C A C G T A G T C A C G T A G T C A C G T A G T C A C G T G T A C C G A T G T C A | BPC6(BBRBPC)/col-BPC6-DAP-Seq(GSE60143)/Homer | 1e-4 | -1.087e+01 | 0.0011 | 7.0 | 20.59% | 1.4 | 0.89% | motif file (matrix) | svg |
| 20 | T G C A C T G A C A T G C T A G C A G T A G T C C G T A A T G C A T G C T A C G G C A T T C A G G T C A G A T C G T A C | ERE(NR),IR3/MCF7-ERa-ChIP-Seq(Unpublished)/Homer | 1e-4 | -1.059e+01 | 0.0013 | 17.0 | 50.00% | 24.2 | 14.87% | motif file (matrix) | svg |
| 21 | T C G A A G C T A C G T A C G T A G T C A G T C A C G T A T C G G A C T A T C G | EWS:ERG-fusion(ETS)/CADO\_ES1-EWS:ERG-ChIP-Seq(SRA014231)/Homer | 1e-4 | -1.059e+01 | 0.0013 | 29.0 | 85.29% | 76.3 | 46.93% | motif file (matrix) | svg |
| 22 | T G C A C T G A A T G C G T C A A C G T A T G C A C G T A C T G A C T G T G C A | ZBTB18(Zf)/HEK293-ZBTB18.GFP-ChIP-Seq(GSE58341)/Homer | 1e-4 | -1.049e+01 | 0.0013 | 19.0 | 55.88% | 31.1 | 19.16% | motif file (matrix) | svg |
| 23 | G A C T A G T C C G A T A C T G C T G A T G A C G T A C C G T A A T C G G C A T C T G A C T A G | Bcl11a(Zf)/HSPC-BCL11A-ChIP-Seq(GSE104676)/Homer | 1e-4 | -1.047e+01 | 0.0013 | 24.0 | 70.59% | 51.6 | 31.75% | motif file (matrix) | svg |
| 24 | T C G A T G C A C A G T T C G A G A T C A G T C C G T A C G T A A C T G A G T C C G T A C G T A T C A G C G A T A G T C | AT5G25475(ABI3VP1)/col-AT5G25475-DAP-Seq(GSE60143)/Homer | 1e-4 | -1.034e+01 | 0.0014 | 33.0 | 97.06% | 105.1 | 64.68% | motif file (matrix) | svg |
| 25 | G C A T C T A G G T A C A G T C C G A T A C T G C T A G C T A G G T A C G C T A | ZNF416(Zf)/HEK293-ZNF416.GFP-ChIP-Seq(GSE58341)/Homer | 1e-4 | -1.032e+01 | 0.0014 | 27.0 | 79.41% | 66.5 | 40.95% | motif file (matrix) | svg |
| 26 | C A T G G C T A C T A G T A C G C G T A T C A G C G T A A C T G C G T A C A T G C T G A C G T A | BPC1(BBRBPC)/colamp-BPC1-DAP-Seq(GSE60143)/Homer | 1e-4 | -1.017e+01 | 0.0015 | 24.0 | 70.59% | 52.3 | 32.20% | motif file (matrix) | svg |
| 27 | C G T A C G T A C T G A C A G T T A G C C G T A G A T C C T A G G C A T C A T G G T A C G A C T | BIM2(bHLH)/col-BIM2-DAP-Seq(GSE60143)/Homer | 1e-4 | -1.004e+01 | 0.0017 | 32.0 | 94.12% | 98.5 | 60.62% | motif file (matrix) | svg |
| 28 | C G T A A T G C C G A T A C G T A G T C C G T A C G T A C G T A C T A G A T C G | TCFL2(HMG)/K562-TCF7L2-ChIP-Seq(GSE29196)/Homer | 1e-4 | -9.821e+00 | 0.0020 | 13.0 | 38.24% | 14.4 | 8.85% | motif file (matrix) | svg |
| 29 | A T G C G C A T T A G C C G A T T A G C G C A T T A G C G C A T A T G C G A C T | GAGA-repeat/Arabidopsis-Promoters/Homer | 1e-4 | -9.776e+00 | 0.0021 | 27.0 | 79.41% | 68.5 | 42.16% | motif file (matrix) | svg |
| 30 | A G T C G A C T C A G T G T A C A G T C A T C G T C A G A C T G G T C A C G T A | Stat3(Stat)/mES-Stat3-ChIP-Seq(GSE11431)/Homer | 1e-4 | -9.339e+00 | 0.0031 | 21.0 | 61.76% | 42.0 | 25.86% | motif file (matrix) | svg |
| 31 | A G T C C G A T A C T G A T C G T G A C G C T A C A T G A T C G T G A C C G A T A C T G T A G C G T A C G T C A | Tlx?(NR)/NPC-H3K4me1-ChIP-Seq(GSE16256)/Homer | 1e-4 | -9.260e+00 | 0.0031 | 11.0 | 32.35% | 10.3 | 6.34% | motif file (matrix) | svg |
| 32 | C T A G C T A G T C G A C T A G C G T A A T C G T C G A A C T G C T G A T C G A C T G A T A C G | FRS9(ND)/col-FRS9-DAP-Seq(GSE60143)/Homer | 1e-4 | -9.260e+00 | 0.0031 | 11.0 | 32.35% | 10.9 | 6.71% | motif file (matrix) | svg |
| 33 | G T A C G A T C C A G T A G T C A G T C A G T C T G C A G A T C C T G A A T G C G T C A A C G T | WT1(Zf)/Kidney-WT1-ChIP-Seq(GSE90016)/Homer | 1e-4 | -9.231e+00 | 0.0031 | 15.0 | 44.12% | 21.8 | 13.44% | motif file (matrix) | svg |
| 34 | T C A G A C T G C A G T A G T C A G T C G T C A C G T A C G T A A C T G C A G T A G T C A G T C C T G A T G C A A G C T | dHNF4(NR)/Fly-HNF4-ChIP-Seq(GSE73675)/Homer | 1e-3 | -9.084e+00 | 0.0035 | 8.0 | 23.53% | 4.2 | 2.61% | motif file (matrix) | svg |
| 35 | T G A C C T G A C T A G C T G A C G T A A G T C C T G A A C G T G C A T T A G C G C A T A T C G G A C T G A C T G A T C | GRE(NR),IR3/RAW264.7-GRE-ChIP-Seq(Unpublished)/Homer | 1e-3 | -8.776e+00 | 0.0046 | 19.0 | 55.88% | 36.6 | 22.49% | motif file (matrix) | svg |
| 36 | G T C A G C A T G C T A C A G T C T A G G A T C C G T A C T G A C G T A C G A T | Oct2(POU,Homeobox)/Bcell-Oct2-ChIP-Seq(GSE21512)/Homer | 1e-3 | -8.773e+00 | 0.0046 | 24.0 | 70.59% | 57.5 | 35.41% | motif file (matrix) | svg |
| 37 | T C A G A C G T T C G A T A G C A G T C C G T A A C T G G T A C A C G T A C T G A T C G A G T C | Atoh1(bHLH)/Cerebellum-Atoh1-ChIP-Seq(GSE22111)/Homer | 1e-3 | -8.750e+00 | 0.0046 | 27.0 | 79.41% | 72.7 | 44.77% | motif file (matrix) | svg |
| 38 | T C G A T A G C G T C A A C T G A C T G C G T A C G T A C T A G A G C T T C A G | ERG(ETS)/VCaP-ERG-ChIP-Seq(GSE14097)/Homer | 1e-3 | -8.749e+00 | 0.0046 | 31.0 | 91.18% | 96.7 | 59.50% | motif file (matrix) | svg |
| 39 | G C T A G C A T G A C T G C A T T C A G G T A C G C T A G C A T C T G A G C T A T A G C G C T A C T G A C G A T C T A G | OCT4-SOX2-TCF-NANOG(POU,Homeobox,HMG)/mES-Oct4-ChIP-Seq(GSE11431)/Homer | 1e-3 | -8.465e+00 | 0.0057 | 19.0 | 55.88% | 37.9 | 23.33% | motif file (matrix) | svg |
| 40 | A G T C C G T A A C G T A G T C A C G T A C T G | Tal1 | 1e-3 | -8.373e+00 | 0.0061 | 33.0 | 97.06% | 113.0 | 69.55% | motif file (matrix) | svg |
| 41 | C A G T C T A G T C G A G T C A C G T A G C T A G C T A C G A T A C G T C A G T A G C T G A T C | SFP1/SacCer-Promoters/Homer | 1e-3 | -8.351e+00 | 0.0061 | 17.0 | 50.00% | 31.0 | 19.05% | motif file (matrix) | svg |
| 42 | C T A G T C G A T G A C A G T C C G T A A C T G G T A C A C G T A C T G A C T G | BHLHA15(bHLH)/NIH3T3-BHLHB8.HA-ChIP-Seq(GSE119782)/Homer | 1e-3 | -8.280e+00 | 0.0063 | 31.0 | 91.18% | 98.5 | 60.64% | motif file (matrix) | svg |
| 43 | T G C A C G T A G T C A A G C T A G T C G C T A T A G C C G A T C T A G G A T C | Gfi1b(Zf)/HPC7-Gfi1b-ChIP-Seq(GSE22178)/Homer | 1e-3 | -8.263e+00 | 0.0063 | 27.0 | 79.41% | 74.2 | 45.69% | motif file (matrix) | svg |
| 44 | G C T A G C A T T A C G A G C T C A T G A C T G C A T G C A T G G A T C A G T C A G T C C T G A A G T C G T A C G C T A | At1g69690(TCP)/colamp-At1g69690-DAP-Seq(GSE60143)/Homer | 1e-3 | -8.169e+00 | 0.0068 | 11.0 | 32.35% | 12.1 | 7.45% | motif file (matrix) | svg |
| 45 | C A G T T C A G T C G A A G T C C G T A A C T G T G A C C G A T A C T G A C T G A C G T A T C G | Atoh7(bHLH)/Retina-Atoh7-CutnRun(GSE156756)/Homer | 1e-3 | -8.148e+00 | 0.0068 | 20.0 | 58.82% | 42.5 | 26.16% | motif file (matrix) | svg |
| 46 | T A C G T G C A A G T C C G T A A C G T T G A C A C G T A C T G A C T G G C A T | TCF4(bHLH)/SHSY5Y-TCF4-ChIP-Seq(GSE96915)/Homer | 1e-3 | -8.101e+00 | 0.0069 | 32.0 | 94.12% | 106.7 | 65.66% | motif file (matrix) | svg |
| 47 | C T A G A G T C T A C G T A C G T G A C C G T A A C T G T A G C G C A T C A T G A T G C A G C T | Ascl1(bHLH)/NeuralTubes-Ascl1-ChIP-Seq(GSE55840)/Homer | 1e-3 | -8.099e+00 | 0.0069 | 29.0 | 85.29% | 87.0 | 53.53% | motif file (matrix) | svg |
| 48 | C G A T C T A G C T G A A T G C C T G A T C G A C G T A C T G A T C G A T A G C A G T C C G T A A C T G T C G A A T G C | Hand2(bHLH)/Mesoderm-Hand2-ChIP-Seq(GSE61475)/Homer | 1e-3 | -8.025e+00 | 0.0072 | 17.0 | 50.00% | 31.2 | 19.21% | motif file (matrix) | svg |
| 49 | C A T G T A C G T A G C G A T C G A T C A T G C G T A C G A C T T C A G A T G C C G A T A T C G C A G T A C T G G T A C | Zic3(Zf)/mES-Zic3-ChIP-Seq(GSE37889)/Homer | 1e-3 | -7.895e+00 | 0.0080 | 10.0 | 29.41% | 10.7 | 6.59% | motif file (matrix) | svg |
| 50 | G A C T C A G T G A T C G A T C A C G T G A T C C T G A T A C G C G T A G T C A | STAT6(Stat)/Macrophage-Stat6-ChIP-Seq(GSE38377)/Homer | 1e-3 | -7.794e+00 | 0.0087 | 27.0 | 79.41% | 76.3 | 46.96% | motif file (matrix) | svg |
| 51 | A T G C A G T C A G C T A G C T A C G T A T C G C G T A C G A T T A G C G A C T | LEF1(HMG)/H1-LEF1-ChIP-Seq(GSE64758)/Homer | 1e-3 | -7.683e+00 | 0.0095 | 30.0 | 88.24% | 94.5 | 58.17% | motif file (matrix) | svg |
| 52 | C G T A T A G C T A G C T G C A A C T G C T A G C G T A C G T A T C A G G A C T | EHF(ETS)/LoVo-EHF-ChIP-Seq(GSE49402)/Homer | 1e-3 | -7.645e+00 | 0.0097 | 32.0 | 94.12% | 108.6 | 66.81% | motif file (matrix) | svg |
| 53 | C T G A T C A G G T A C T G C A A G T C C G T A A G T C A C T G A C G T A C T G | MNT(bHLH)/HepG2-MNT-ChIP-Seq(Encode)/Homer | 1e-3 | -7.597e+00 | 0.0099 | 31.0 | 91.18% | 101.8 | 62.63% | motif file (matrix) | svg |
| 54 | G T A C C T G A T A G C C G T A G C T A T C G A T G C A T G A C C T A G G T C A A G T C C G T A C T G A C T G A C G T A | At1g14580(C2H2)/colamp-At1g14580-DAP-Seq(GSE60143)/Homer | 1e-3 | -7.529e+00 | 0.0104 | 16.0 | 47.06% | 29.9 | 18.41% | motif file (matrix) | svg |
| 55 | A C T G A C T G C T A G G A T C A G T C A G T C C T G A A G T C | At5g08330(TCP)/col-At5g08330-DAP-Seq(GSE60143)/Homer | 1e-3 | -7.407e+00 | 0.0116 | 17.0 | 50.00% | 33.8 | 20.82% | motif file (matrix) | svg |
| 56 | C A G T T C A G G A T C A C T G A C G T C T A G A C T G A C T G G A C T C T A G | Egr1(Zf)/K562-Egr1-ChIP-Seq(GSE32465)/Homer | 1e-3 | -7.338e+00 | 0.0120 | 18.0 | 52.94% | 37.2 | 22.91% | motif file (matrix) | svg |
| 57 | C A T G A C T G A G C T A T G C C G T A A T G C G T A C G A C T T A C G C T G A A C T G A C T G G C A T A T G C C T G A | THRb(NR)/HepG2-THRb.Flag-ChIP-Seq(Encode)/Homer | 1e-3 | -7.338e+00 | 0.0120 | 18.0 | 52.94% | 37.2 | 22.89% | motif file (matrix) | svg |
| 58 | G C A T A G C T A C G T A C G T A C T G A C G T G A T C A C G T A C G T A G C T C G A T G C A T A G T C G A C T C A G T | IDD5(C2H2)/colamp-IDD5-DAP-Seq(GSE60143)/Homer | 1e-3 | -7.242e+00 | 0.0129 | 30.0 | 88.24% | 97.0 | 59.69% | motif file (matrix) | svg |
| 59 | A T G C T G C A A G T C C G T A A G T C A G T C A C G T A C T G A T G C G T C A | E2A(bHLH),near\_PU.1/Bcell-PU.1-ChIP-Seq(GSE21512)/Homer | 1e-3 | -7.204e+00 | 0.0132 | 29.0 | 85.29% | 90.9 | 55.95% | motif file (matrix) | svg |
| 60 | T A G C G T A C A G T C G T A C C G A T A G T C A G T C A G T C A G T C A G T C C G T A G A T C | Zfp281(Zf)/ES-Zfp281-ChIP-Seq(GSE81042)/Homer | 1e-3 | -7.159e+00 | 0.0136 | 4.0 | 11.76% | 0.0 | 0.00% | motif file (matrix) | svg |
| 61 | A G T C G A C T A G C T C G A T A T C G G C T A C G A T A T C G C G A T A C T G T A C G A C G T | Tcf7(HMG)/GM12878-TCF7-ChIP-Seq(Encode)/Homer | 1e-3 | -7.133e+00 | 0.0137 | 21.0 | 61.76% | 50.6 | 31.14% | motif file (matrix) | svg |
| 62 | A C G T C T A G G T C A G C T A C G A T G C T A G C T A G C A T C A G T G A C T T G C A C A G T | POU4F3(POU,Homeobox)/MEF-Pou4f3-ChIP-Seq(GSE150279)/Homer | 1e-3 | -7.073e+00 | 0.0143 | 20.0 | 58.82% | 46.3 | 28.49% | motif file (matrix) | svg |
| 63 | C T G A T C A G A G T C C G T A A T C G A T G C C G A T A C T G A G T C G A C T A T C G A G T C | MyoD(bHLH)/Myotube-MyoD-ChIP-Seq(GSE21614)/Homer | 1e-3 | -7.060e+00 | 0.0143 | 18.0 | 52.94% | 38.2 | 23.49% | motif file (matrix) | svg |
| 64 | A C T G T G A C G T A C C G T A A G T C T A C G A C G T A C T G G T C A A G T C | NPAS2(bHLH)/Liver-NPAS2-ChIP-Seq(GSE39860)/Homer | 1e-3 | -7.027e+00 | 0.0146 | 30.0 | 88.24% | 97.6 | 60.07% | motif file (matrix) | svg |
| 65 | T C A G C A T G C A T G A C T G A C T G A G C T A C T G A C G T A C T G C A G T A T G C A G T C | KLF10(Zf)/HEK293-KLF10.GFP-ChIP-Seq(GSE58341)/Homer | 1e-3 | -6.931e+00 | 0.0158 | 14.0 | 41.18% | 24.7 | 15.18% | motif file (matrix) | svg |
| 66 | C A G T A C T G T C A G T G C A G C T A A T G C T C G A A T C G G T C A T G C A | ZNF189(Zf)/HEK293-ZNF189.GFP-ChIP-Seq(GSE58341)/Homer | 1e-2 | -6.886e+00 | 0.0162 | 23.0 | 67.65% | 60.4 | 37.17% | motif file (matrix) | svg |
| 67 | C T G A C G A T C A T G A T C G G C A T C A T G G C T A A G T C | ASHR1(ND)/col-ASHR1-DAP-Seq(GSE60143)/Homer | 1e-2 | -6.814e+00 | 0.0172 | 30.0 | 88.24% | 98.6 | 60.71% | motif file (matrix) | svg |
| 68 | T C G A T C G A C T G A C G T A A C T G A T G C A C G T A G T C | Lola-I(Zf)/Embryo-LolaI-ChIP-Seq(GSE200870)/Homer | 1e-2 | -6.810e+00 | 0.0172 | 25.0 | 73.53% | 70.2 | 43.21% | motif file (matrix) | svg |
| 69 | G T C A T G C A T G C A G C T A C G T A G C T A G C T A G C T A | REM19(REM)/colamp-REM19-DAP-Seq(GSE60143)/Homer | 1e-2 | -6.729e+00 | 0.0182 | 31.0 | 91.18% | 105.3 | 64.81% | motif file (matrix) | svg |
| 70 | C T A G T G A C G A C T A T C G T C G A A G T C C T A G C A G T C T A G A T C G G T A C T C G A | O2(bZIP)/Corn-O2-ChIP-Seq(GSE63991)/Homer | 1e-2 | -6.608e+00 | 0.0202 | 9.0 | 26.47% | 10.8 | 6.67% | motif file (matrix) | svg |
| 71 | C T A G C A T G A C G T A G T C G C T A A G C T A G T C A G C T T C A G C T G A A C T G C A T G G C A T A T G C C G T A | THRa(NR)/C17.2-THRa-ChIP-Seq(GSE38347)/Homer | 1e-2 | -6.608e+00 | 0.0202 | 14.0 | 41.18% | 25.1 | 15.42% | motif file (matrix) | svg |
| 72 | C T A G T C G A C G A T A C T G C G T A T A C G A G C T T G A C G C T A A C G T G A T C T A G C | Fosl2(bZIP)/3T3L1-Fosl2-ChIP-Seq(GSE56872)/Homer | 1e-2 | -6.559e+00 | 0.0207 | 17.0 | 50.00% | 36.0 | 22.16% | motif file (matrix) | svg |
| 73 | C A T G A C G T A G T C G A T C G A T C G A T C G C T A C T A G C T A G C T A G T C A G T C G A | EBF1(EBF)/Near-E2A-ChIP-Seq(GSE21512)/Homer | 1e-2 | -6.538e+00 | 0.0207 | 19.0 | 55.88% | 44.8 | 27.54% | motif file (matrix) | svg |
| 74 | A C G T G A C T A T G C G C T A C T G A C T A G A C T G G A C T A G T C C G T A | Nr5a2(NR)/mES-Nr5a2-ChIP-Seq(GSE19019)/Homer | 1e-2 | -6.538e+00 | 0.0207 | 19.0 | 55.88% | 44.7 | 27.52% | motif file (matrix) | svg |
| 75 | C T G A T C G A C G T A A T G C C G T A C G T A C G A T C T A G T C A G G A T C | Sox15(HMG)/CPA-Sox15-ChIP-Seq(GSE62909)/Homer | 1e-2 | -6.470e+00 | 0.0217 | 34.0 | 100.00% | 131.3 | 80.78% | motif file (matrix) | svg |
| 76 | A C T G T C A G A G C T G A C T C A T G A G T C A G T C G C T A C G A T C T A G T C A G G T A C C T G A T C G A | Rfx1(HTH)/NPC-H3K4me1-ChIP-Seq(GSE16256)/Homer | 1e-2 | -6.415e+00 | 0.0226 | 11.0 | 32.35% | 16.5 | 10.16% | motif file (matrix) | svg |
| 77 | A G T C C T A G C T A G A G T C G A T C G T A C A G T C C T A G A G T C A G T C A G T C G T A C | Sp2(Zf)/HEK293-Sp2.eGFP-ChIP-Seq(Encode)/Homer | 1e-2 | -6.392e+00 | 0.0228 | 24.0 | 70.59% | 67.7 | 41.68% | motif file (matrix) | svg |
| 78 | C A T G A C T G C T A G T C G A T C G A T C G A T C G A T C A G T C A G T C A G T G A C T G A C C G T A A C T G T G C A C G A T A C T G | RBPJ:Ebox(?,bHLH)/Panc1-Rbpj1-ChIP-Seq(GSE47459)/Homer | 1e-2 | -6.359e+00 | 0.0230 | 8.0 | 23.53% | 8.9 | 5.46% | motif file (matrix) | svg |
| 79 | G A T C G A T C A G T C G T A C C G A T G T A C G T A C A G T C A G T C A G T C G C T A G A T C | ZNF148(Zf)/MDAMB231-ZNF148-ChIP-Seq(GSE147020)/Homer | 1e-2 | -6.359e+00 | 0.0230 | 8.0 | 23.53% | 8.9 | 5.47% | motif file (matrix) | svg |
| 80 | C G T A C T G A C T A G C G T A C G T A A G T C C G T A C A G T G C A T G T C A C G A T A C T G A C G T G C A T G A T C | PGR(NR)/EndoStromal-PGR-ChIP-Seq(GSE69539)/Homer | 1e-2 | -6.296e+00 | 0.0242 | 17.0 | 50.00% | 37.5 | 23.08% | motif file (matrix) | svg |
| 81 | T A G C G C A T A G T C G A T C A T G C G A C T C T A G A C T G A C T G C T G A A C T G C T A G A G T C T G A C C G A T | GLIS3(Zf)/Thyroid-Glis3.GFP-ChIP-Seq(GSE103297)/Homer | 1e-2 | -6.223e+00 | 0.0257 | 23.0 | 67.65% | 63.2 | 38.89% | motif file (matrix) | svg |
| 82 | A G C T A G T C C G T A A G T C C T A G A C G T A C T G T C A G C G A T C A T G | PIF5ox(bHLH)/Arabidopsis-PIF5ox-ChIP-Seq(GSE35062)/Homer | 1e-2 | -6.161e+00 | 0.0270 | 29.0 | 85.29% | 96.0 | 59.05% | motif file (matrix) | svg |
| 83 | T G A C G C T A T G A C C G T A T C A G G A T C C G T A C A T G C A T G C T A G C T A G C T A G | Unknown-ESC-element(?)/mES-Nanog-ChIP-Seq(GSE11724)/Homer | 1e-2 | -6.091e+00 | 0.0286 | 12.0 | 35.29% | 20.1 | 12.35% | motif file (matrix) | svg |
| 84 | C T A G A G C T C A T G C A T G C T A G A T G C G A T C G T A C G T A C C T G A A G T C G C T A G C T A G C A T A T C G | TCP7(TCP)/col-TCP7-DAP-Seq(GSE60143)/Homer | 1e-2 | -6.083e+00 | 0.0286 | 16.0 | 47.06% | 34.1 | 20.96% | motif file (matrix) | svg |
| 85 | T C G A T G A C G C A T A G C T C A G T G A T C G C T A G A T C G A C T A C G T G C A T A G T C | PRDM1(Zf)/Hela-PRDM1-ChIP-Seq(GSE31477)/Homer | 1e-2 | -5.974e+00 | 0.0314 | 24.0 | 70.59% | 69.5 | 42.79% | motif file (matrix) | svg |
| 86 | C G T A G C T A C G A T A C G T A G C T A G C T C T G A G T C A C G T A G C T A | Unknown6/Drosophila-Promoters/Homer | 1e-2 | -5.902e+00 | 0.0333 | 33.0 | 97.06% | 124.2 | 76.45% | motif file (matrix) | svg |
| 87 | G A C T C G T A A C G T A C T G A G T C C G T A C T G A C G T A C A G T A C G T G T C A T C A G | Brn1(POU,Homeobox)/NPC-Brn1-ChIP-Seq(GSE35496)/Homer | 1e-2 | -5.806e+00 | 0.0359 | 26.0 | 76.47% | 80.8 | 49.71% | motif file (matrix) | svg |
| 88 | T C G A T A G C T G C A A C T G A C T G C G T A C G T A C T A G G A C T T A C G | ETS1(ETS)/Jurkat-ETS1-ChIP-Seq(GSE17954)/Homer | 1e-2 | -5.806e+00 | 0.0359 | 26.0 | 76.47% | 80.8 | 49.75% | motif file (matrix) | svg |
| 89 | C G T A T G A C T A G C T G C A A C T G A C T G C G T A C G T A T C A G G A C T | ELF3(ETS)/PDAC-ELF3-ChIP-Seq(GSE64557)/Homer | 1e-2 | -5.806e+00 | 0.0359 | 23.0 | 67.65% | 65.8 | 40.47% | motif file (matrix) | svg |
| 90 | C T A G A C T G C T A G T C A G T C A G T A C G C T A G A C T G | Maz(Zf)/HepG2-Maz-ChIP-Seq(GSE31477)/Homer | 1e-2 | -5.629e+00 | 0.0419 | 15.0 | 44.12% | 32.6 | 20.06% | motif file (matrix) | svg |
| 91 | A T G C T C G A A G T C A G C T A C G T G T A C A G T C G C T A C T A G C A T G G T C A C T G A T C A G A G T C | Stat3+il21(Stat)/CD4-Stat3-ChIP-Seq(GSE19198)/Homer | 1e-2 | -5.575e+00 | 0.0432 | 24.0 | 70.59% | 71.7 | 44.14% | motif file (matrix) | svg |
| 92 | A G T C G A C T A C T G G A T C G T A C C G T A T G A C A G T C C G A T A G C T A C G T A C G T C T A G G A C T C T G A | ZNF7(Zf)/HepG2-ZNF7.Flag-ChIP-Seq(Encode)/Homer | 1e-2 | -5.575e+00 | 0.0432 | 24.0 | 70.59% | 71.7 | 44.15% | motif file (matrix) | svg |
| 93 | C G A T C A G T G T C A G C T A C A G T A G C T A C G T A C T G A G T C C G T A A C G T A C T G A C G T C T G A T C G A | FUS3(ABI3VP1)/col-FUS3-DAP-Seq(GSE60143)/Homer | 1e-2 | -5.511e+00 | 0.0456 | 31.0 | 91.18% | 112.0 | 68.91% | motif file (matrix) | svg |
| 94 | A C T G G T C A A C G T G C A T C G A T T C A G G T A C G T C A A C G T C T G A | Oct11(POU,Homeobox)/NCIH1048-POU2F3-ChIP-seq(GSE115123)/Homer | 1e-2 | -5.422e+00 | 0.0488 | 26.0 | 76.47% | 82.1 | 50.51% | motif file (matrix) | svg |
| 95 | C T G A A G C T A C G T A C G T A G T C G A C T G A C T C T G A C T G A C T A G C G T A C G T A | STAT6(Stat)/CD4-Stat6-ChIP-Seq(GSE22104)/Homer | 1e-2 | -5.422e+00 | 0.0488 | 26.0 | 76.47% | 82.9 | 51.01% | motif file (matrix) | svg |
| 96 | T C G A T A G C G T C A A C T G C T A G C G T A C G A T A C T G A C G T A C T G A C T G A C G T | ETS:RUNX(ETS,Runt)/Jurkat-RUNX1-ChIP-Seq(GSE17954)/Homer | 1e-2 | -5.330e+00 | 0.0529 | 3.0 | 8.82% | 0.6 | 0.35% | motif file (matrix) | svg |
| 97 | A C G T A T G C A C G T A C G T A G C T A G T C A G C T A G C T A G C T A G C T A G C T | hTCT(CPE) | 1e-2 | -5.275e+00 | 0.0553 | 33.0 | 97.06% | 127.2 | 78.29% | motif file (matrix) | svg |
| 98 | C T G A A G T C C G A T A G C T A T G C G T A C A C G T A T C G C A G T G C A T | Elf4(ETS)/BMDM-Elf4-ChIP-Seq(GSE88699)/Homer | 1e-2 | -5.236e+00 | 0.0569 | 26.0 | 76.47% | 83.0 | 51.10% | motif file (matrix) | svg |
| 99 | A G T C A G T C C G A T A C G T A C G T A C T G A C G T A G C T A G T C A G T C | Sox4(HMG)/proB-Sox4-ChIP-Seq(GSE50066)/Homer | 1e-2 | -5.200e+00 | 0.0584 | 29.0 | 85.29% | 100.0 | 61.57% | motif file (matrix) | svg |
| 100 | G A C T C T A G G A T C C A G T A C T G C T G A A T G C G C A T A T G C C T G A | MafA(bZIP)/Islet-MafA-ChIP-Seq(GSE30298)/Homer | 1e-2 | -5.200e+00 | 0.0584 | 25.0 | 73.53% | 78.5 | 48.33% | motif file (matrix) | svg |
| 101 | A G T C G A C T C A G T A C T G C T A G T G A C G C T A A T G C G C A T A T C G C G A T A C T G G A T C G T A C G T C A C T G A | NF1(CTF)/LNCAP-NF1-ChIP-Seq(Unpublished)/Homer | 1e-2 | -5.169e+00 | 0.0591 | 12.0 | 35.29% | 23.5 | 14.45% | motif file (matrix) | svg |
| 102 | G T A C G T A C G T C A G C T A C G T A C G T A C G T A C T A G C T A G C T A G | SEP3(MADS)/Arabidoposis-Flower-Sep3-ChIP-Seq/Homer | 1e-2 | -5.117e+00 | 0.0616 | 34.0 | 100.00% | 137.9 | 84.86% | motif file (matrix) | svg |
| 103 | C T G A A C G T A C G T A C G T A G T C G A C T C G A T C T G A A C T G C G T A C G T A T C G A | STAT5(Stat)/mCD4+-Stat5-ChIP-Seq(GSE12346)/Homer | 1e-2 | -5.107e+00 | 0.0617 | 17.0 | 50.00% | 42.7 | 26.30% | motif file (matrix) | svg |
| 104 | T C A G A G C T A T G C C G T A A G C T T C A G C A G T A C T G C T G A A G T C | MITF(bHLH)/MastCells-MITF-ChIP-Seq(GSE48085)/Homer | 1e-2 | -5.018e+00 | 0.0661 | 29.0 | 85.29% | 101.8 | 62.64% | motif file (matrix) | svg |
| 105 | A G T C C G T A C G A T A G T C G T C A A G T C A C G T C T G A | Unknown2/Drosophila-Promoters/Homer | 1e-2 | -5.018e+00 | 0.0661 | 29.0 | 85.29% | 101.1 | 62.24% | motif file (matrix) | svg |
| 106 | C T A G T C G A A C G T A C T G C G T A T A G C C G A T G T A C C G T A A G C T G A T C G T A C | Jun-AP1(bZIP)/K562-cJun-ChIP-Seq(GSE31477)/Homer | 1e-2 | -4.992e+00 | 0.0672 | 10.0 | 29.41% | 17.8 | 10.94% | motif file (matrix) | svg |
| 107 | A C T G A G C T A G T C G T C A A G C T T C A G A T G C G A T C G C A T A T C G T C G A T A G C C G A T C A T G T A G C | Pax8(Paired,Homeobox)/Thyroid-Pax8-ChIP-Seq(GSE26938)/Homer | 1e-2 | -4.885e+00 | 0.0741 | 6.0 | 17.65% | 6.2 | 3.84% | motif file (matrix) | svg |
| 108 | C T A G C T A G T C G A C G T A A T G C C G T A A T C G T C G A T A C G G C A T A C T G C A G T T A G C G A T C G A C T | MRE(NR)/Neuro2A-NR3C2-ChIPnexus(GSE115417)/Homer | 1e-2 | -4.755e+00 | 0.0829 | 31.0 | 91.18% | 115.3 | 70.98% | motif file (matrix) | svg |
| 109 | A T G C A G T C G A T C C G T A A C G T A C G T A C T G A C G T A G C T G A T C | Sox2(HMG)/mES-Sox2-ChIP-Seq(GSE11431)/Homer | 1e-2 | -4.755e+00 | 0.0829 | 31.0 | 91.18% | 115.4 | 71.03% | motif file (matrix) | svg |
| 110 | C T A G C A T G C A T G A G C T G A C T C G T A G A T C A G C T G T C A A T G C T C G A C T A G C A T G A C G T A G T C C T G A | LXRE(NR),DR4/RAW-LXRb.biotin-ChIP-Seq(GSE21512)/Homer | 1e-2 | -4.720e+00 | 0.0850 | 5.0 | 14.71% | 4.4 | 2.69% | motif file (matrix) | svg |
| 111 | T A C G T C G A C A G T A C T G G C T A A T G C C G A T G T A C C G T A A C T G T A G C C G T A | NF-E2(bZIP)/K562-NFE2-ChIP-Seq(GSE31477)/Homer | 1e-2 | -4.718e+00 | 0.0850 | 4.0 | 11.76% | 2.4 | 1.46% | motif file (matrix) | svg |
| 112 | C T A G G A C T G A T C A C G T A T C G A G C T C T G A A T C G C G A T C T A G G A T C G A C T C A T G A T C G G T A C G A C T A G T C G C A T A G C T C G A T | ZNF382(Zf)/HEK293-ZNF382.GFP-ChIP-Seq(GSE58341)/Homer | 1e-2 | -4.718e+00 | 0.0850 | 4.0 | 11.76% | 2.0 | 1.24% | motif file (matrix) | svg |
