## Supplementary Data 11 for "Evolutionary insights into glucose production in vertebrate development: new findings from Arctic lamprey (*Lethenteron camtschaticum*)": Suplementary Data11_C_g6pc_motif.html

lampreyg6pc2\_ex\_motif/ - Homer de novo Motif Results


### Homer *de novo* Motif Results (lampreyg6pc2\_ex\_motif/)

Non-redundant Motif File of Results  
Known Motif Enrichment Results  
Gene Ontology Enrichment Results  
If Homer is having trouble matching a motif to a known motif, try copy/pasting the matrix file into
STAMP  
More information on motif finding results: HOMER
| Description of Results
| Tips
  
Total target sequences = 8  
Total background sequences = 37  
\* - possible false positive  

|  |  |  |  |  |  |  |  |  |
| --- | --- | --- | --- | --- | --- | --- | --- | --- |
| Rank | Motif | P-value | log P-pvalue | % of Targets | % of Background | STD(Bg STD) | Best Match/Details | Motif File |
| 1 \* | A C T G G A T C A C G T A C G T C G A T A G T C A T C G A G C T A C T G C T G A | 1e-7 | -1.699e+01 | 100.00% | 3.59% | 635.2bp (446.4bp) | SWI4(MacIsaac)/Yeast(0.703) More Information \| Similar Motifs Found | motif file (matrix) |
| 2 \* | A C T G C G T A C G T A C G T A A G T C A C T G A C G T A G T C A C T G A G T C | 1e-6 | -1.555e+01 | 87.50% | 0.00% | 783.2bp (0.0bp) | HSFA1B/MA2019.2/Jaspar(0.715) More Information \| Similar Motifs Found | motif file (matrix) |
| 3 \* | A G T C A C G T A G T C A G T C A G T C A G T C G A C T A G T C A G T C A T G C | 1e-6 | -1.555e+01 | 87.50% | 1.30% | 855.5bp (0.0bp) | MAZ/MA1522.2/Jaspar(0.865) More Information \| Similar Motifs Found | motif file (matrix) |
| 4 \* | A C G T A G T C C G T A A C G T C G T A A G C T A C G T A T G C A C G T A C G T | 1e-6 | -1.555e+01 | 87.50% | 1.79% | 503.0bp (0.0bp) | KAN2(G2like)/colamp-KAN2-DAP-Seq(GSE60143)/Homer(0.808) More Information \| Similar Motifs Found | motif file (matrix) |
| 5 \* | G T C A C G T A G T A C G T C A C G T A A G T C G T C A G T C A A G T C C G T A | 1e-6 | -1.555e+01 | 87.50% | 1.30% | 925.8bp (0.0bp) | PB0122.1\_Foxk1\_2/Jaspar(0.896) More Information \| Similar Motifs Found | motif file (matrix) |
| 6 \* | C G T A C G T A C G T A A C T G A C G T C G T A C G T A C G T A C G T A A C G T C G T A C G T A | 1e-6 | -1.555e+01 | 87.50% | 0.00% | 509.5bp (0.0bp) | D1/MA2306.1/Jaspar(0.779) More Information \| Similar Motifs Found | motif file (matrix) |
| 7 \* | C A T G C G T A C G T A G T C A C A T G G A T C G C A T C G A T A T G C G T C A C G T A C G T A | 1e-6 | -1.555e+01 | 87.50% | 0.00% | 700.9bp (0.0bp) | TCX6/MA1380.1/Jaspar(0.758) More Information \| Similar Motifs Found | motif file (matrix) |
| 8 \* | C T G A A G T C A G T C A G C T C G T A A G T C A C T G A C G T A C T G C G T A A G T C A C G T | 1e-6 | -1.555e+01 | 87.50% | 0.00% | 611.8bp (0.0bp) | hif-1/MA2168.1/Jaspar(0.771) More Information \| Similar Motifs Found | motif file (matrix) |
| 9 \* | C A T G A G C T C T A G C G T A T A G C C G A T A C T G T G A C T C G A C A T G T C A G T A C G | 1e-6 | -1.555e+01 | 87.50% | 0.00% | 681.9bp (0.0bp) | MSANTD3/MA1523.2/Jaspar(0.669) More Information \| Similar Motifs Found | motif file (matrix) |
| 10 \* | C G T A C G A T A G C T C G T A C G T A G C T A C G T A A C G T C G A T C G T A C G T A A C G T | 1e-6 | -1.555e+01 | 87.50% | 0.00% | 576.4bp (0.0bp) | retn/MA2265.1/Jaspar(0.785) More Information \| Similar Motifs Found | motif file (matrix) |
| 11 \* | A T G C C T A G A G T C C G T A C G T A A G T C C T G A A G T C T C G A C G T A A C T G A G T C | 1e-6 | -1.555e+01 | 87.50% | 0.00% | 777.2bp (0.0bp) | RAV1/MA0582.2/Jaspar(0.731) More Information \| Similar Motifs Found | motif file (matrix) |
| 12 \* | A T C G A G T C C G A T A G T C C G T A A G T C C G T A C G T A | 1e-6 | -1.555e+01 | 87.50% | 2.48% | 784.9bp (0.0bp) | PB0208.1\_Zscan4\_2/Jaspar(0.794) More Information \| Similar Motifs Found | motif file (matrix) |
| 13 \* | A G T C C G A T A G T C A G C T A G T C A C G T A G T C G T A C A G C T A G T C C G A T A C G T | 1e-5 | -1.349e+01 | 87.50% | 4.79% | 446.6bp (184.4bp) | BPC1(BBRBPC)/colamp-BPC1-DAP-Seq(GSE60143)/Homer(0.733) More Information \| Similar Motifs Found | motif file (matrix) |
| 14 \* | A G T C A C T G A G T C A C G T A C G T A C T G A G T C C G T A | 1e-5 | -1.349e+01 | 87.50% | 2.90% | 564.1bp (134.1bp) | MBNL1(Znf)/Homo\_sapiens-RNCMPT00038-PBM/HughesRNA(0.841) More Information \| Similar Motifs Found | motif file (matrix) |
| 15 \* | A C G T A C G T A G T C A C T G A C T G A C G T C G T A C G T A C G T A A C T G | 1e-5 | -1.258e+01 | 75.00% | 0.00% | 559.7bp (0.0bp) | ARO80/MA0273.2/Jaspar(0.698) More Information \| Similar Motifs Found | motif file (matrix) |
| 16 \* | C G A T A C G T A T C G C G T A C G T A A C G T C T G A C G T A A G T C C G T A | 1e-5 | -1.258e+01 | 75.00% | 0.00% | 472.9bp (0.0bp) | ara/dmmpmm(Noyes\_hd)/fly(0.739) More Information \| Similar Motifs Found | motif file (matrix) |
| 17 \* | A C T G A C G T C A T G A G C T A C T G A C T G A C G T A C G T A C G T A C T G | 1e-5 | -1.258e+01 | 75.00% | 2.13% | 769.5bp (385.4bp) | RUNX(Runt)/HPC7-Runx1-ChIP-Seq(GSE22178)/Homer(0.785) More Information \| Similar Motifs Found | motif file (matrix) |
| 18 \* | A G T C A G T C A G T C A G T C A G T C A G T C C G T A A G T C A G T C C G T A A G T C A G T C | 1e-5 | -1.258e+01 | 75.00% | 0.00% | 948.0bp (0.0bp) | ZNF740/MA0753.3/Jaspar(0.762) More Information \| Similar Motifs Found | motif file (matrix) |
| 19 \* | C G T A G T C A C G T A G A C T A G T C C G T A C T G A C G A T C G T A A C G T | 1e-4 | -9.990e+00 | 62.50% | 1.79% | 793.3bp (0.0bp) | ONECUT1/MA0679.3/Jaspar(0.884) More Information \| Similar Motifs Found | motif file (matrix) |
| 20 \* | C G A T G T C A A C G T C G T A A C G T C G T A A G C T C T G A A G C T C T G A | 1e-4 | -9.990e+00 | 62.50% | 1.23% | 549.4bp (0.7bp) | SeqBias: TA-repeat(0.985) More Information \| Similar Motifs Found | motif file (matrix) |
| 21 \* | C G A T G C A T A C G T A C G T A C G T C G A T C T G A C G A T A C G T G T C A A C G T G A C T | 1e-4 | -9.990e+00 | 62.50% | 0.00% | 323.8bp (0.0bp) | PB0148.1\_Mtf1\_2/Jaspar(0.831) More Information \| Similar Motifs Found | motif file (matrix) |
| 22 \* | A C G T C T A G C G T A A C G T A C T G C G T A A C G T A T C G C T G A C G A T A T C G C T G A | 1e-4 | -9.990e+00 | 62.50% | 0.00% | 967.3bp (0.0bp) | ZML2(C2C2gata)/col-ZML2-DAP-Seq(GSE60143)/Homer(0.900) More Information \| Similar Motifs Found | motif file (matrix) |
| 23 \* | T A G C A C T G A C G T A C T G A C T G A T C G A C G T A C T G C G T A C T A G | 1e-4 | -9.849e+00 | 87.50% | 13.13% | 840.3bp (774.8bp) | CRZ1(MacIsaac)/Yeast(0.803) More Information \| Similar Motifs Found | motif file (matrix) |
| 24 \* | A C T G A C T G C G T A A C G T A C G T C G T A A C G T A C G T | 1e-4 | -9.339e+00 | 75.00% | 7.66% | 804.6bp (814.0bp) | dve/MA0915.2/Jaspar(0.866) More Information \| Similar Motifs Found | motif file (matrix) |
| 25 \* | C G T A C G T A C G T A A C T G C G T A C G T A A G T C A G T C | 1e-4 | -9.339e+00 | 75.00% | 6.28% | 655.5bp (362.9bp) | OPI1/MA0349.2/Jaspar(0.791) More Information \| Similar Motifs Found | motif file (matrix) |
| 26 \* | C G T A A C T G A C T G A C G T A C G T A C G T A C G T A C T G A C T G A G T C A C T G A C T G | 1e-3 | -7.663e+00 | 50.00% | 0.00% | 222.5bp (0.0bp) | E2F7/MA0758.1/Jaspar(0.699) More Information \| Similar Motifs Found | motif file (matrix) |
