## Supplementary figures and images for "Evolutionary insights into glucose production in vertebrate development: new findings from Arctic lamprey (*Lethenteron camtschaticum*)"

### Supplementary Figure 1

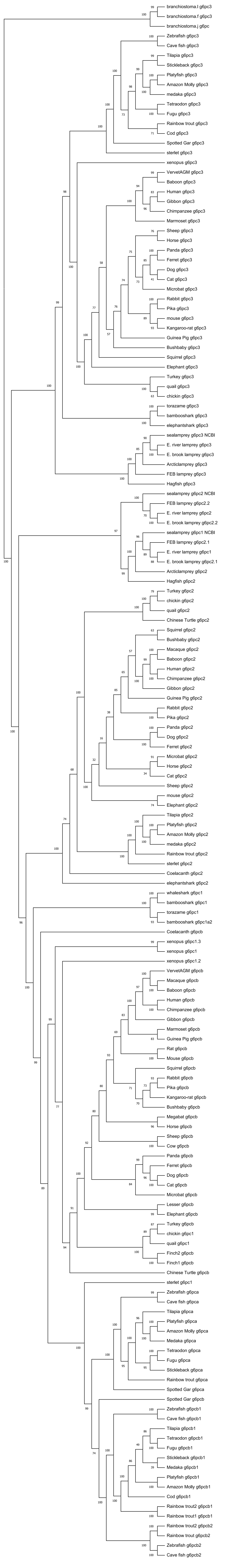

### Supplementary Figure 2

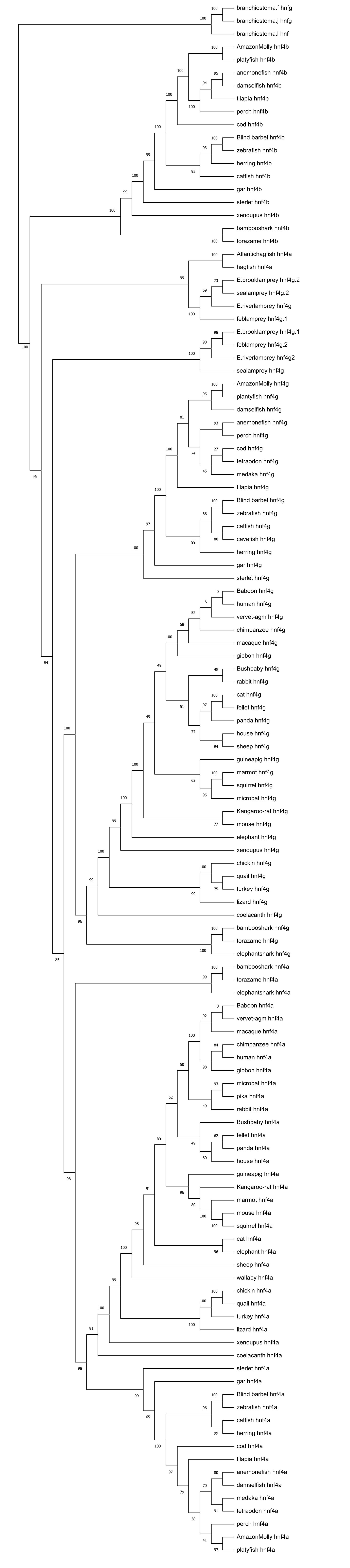
